# Adeno-associated viral delivery of Env-specific antibodies prevents SIV rebound after discontinuing antiretroviral therapy

**DOI:** 10.1101/2024.05.30.593694

**Authors:** Vadim A. Klenchin, Natasha M. Clark, Nida K. Keles, Saverio Capuano, Rosemarie Mason, Guangping Gao, Aimee Broman, Emek Kose, Taina T. Immonen, Christine M. Fennessey, Brandon F. Keele, Jeffrey D. Lifson, Mario Roederer, Matthew R. Gardner, David T. Evans

## Abstract

An alternative to lifelong antiretroviral therapy (ART) is needed to achieve durable control of HIV-1. Here we show that adeno-associated virus (AAV)-delivery of two rhesus macaque antibodies to the SIV envelope glycoprotein (Env) with potent neutralization and antibody-dependent cellular cytotoxicity can prevent viral rebound in macaques infected with barcoded SIV_mac_239M after discontinuing suppressive ART. Following AAV administration, sustained antibody expression with minimal anti-drug antibody responses was achieved in all but one animal. After ART withdrawal, SIV replication rebounded within two weeks in all of the control animals but remained below the threshold of detection in plasma (<15 copies/mL) for more than a year in four of the eight animals that received AAV vectors encoding Env-specific antibodies. Viral sequences from animals with delayed rebound exhibited restricted barcode diversity and antibody escape. Thus, sustained expression of antibodies with potent antiviral activity can afford durable, ART-free containment of pathogenic SIV infection.

## Introduction

Advances in antiretroviral therapy (ART) have narrowed but not eliminated the life expectancy gap between people living with HIV-1 and the general population^1,2^. Treatment challenges such as access to drugs, non-adherence to treatment regimens and side effects still exist. Long-lasting, gene therapy-based approaches employing broadly neutralizing antibodies (bnAbs) offer an attractive alternative because there is now ample evidence that passive infusion of potent bnAbs can protect against immunodeficiency virus infection^3–6^ and suppress virus replication in humans and nonhuman primates^7–10^. bnAbs can also eliminate HIV-1-infected cells by Fc-mediated effector functions such as antibody-dependent cellular cytotoxicity (ADCC)^11–15^.

Transduction of long-lived muscle cells with adeno-associated virus (AAV) vectors encoding bnAbs can afford high-level, long-term expression of bnAbs that protect against HIV-1 in humanized mice^16,17^ and simian-human immunodeficiency virus (SHIV) in rhesus macaques^18–22^. AAV-delivery of HIV-1 bnAbs has been demonstrated to be safe and well tolerated in humans^23,24^. However, the antiviral activity of AAV-delivered antibodies is typically limited by antibodies to the vectored antibodies, termed anti-drug antibody (ADA) responses, that severely impair bnAb expression and the ability to control virus replication^25,26^. In this study, we demonstrate that sustained expression of natural, species-matched antibodies with potent neutralization and ADCC can be achieved in most animals by AAV delivery, and that these antibodies can contain the replication of SIV_mac_239, which is considered to be a tier 3 neutralization-resistant virus, below the limit of detection in 50% of treated macaques for more than a year after cessation of ART. This unprecedented, ART-free containment of a highly pathogenic strain of SIV in macaques illustrates the potential for AAV-delivery of bnAbs to delay, perhaps indefinitely, the resurgence of HIV-1 replication in humans after discontinuing antiretroviral drug therapy.

## Results

### Antibodies and AAV vector design

Two rhesus macaque antibodies specific for the SIV envelope glycoprotein, ITS61.01 and ITS103.01^27^, and a control antibody to the respiratory syncytial virus fusion protein, 17-HD9^28^, were selected for AAV delivery. ITS61.01 and ITS103.01 both direct efficient ADCC against SIV_mac_239-infected cells (Extended Data Fig. 1a) and ITS103.01 also potently neutralizes SIV_mac_239 infectivity (Extended Data Fig. 1b)^29^. For optimal expression in long-lived muscle cells, the sequences for these antibodies were cloned into an AAV vector downstream of a cytomegalovirus CMV enhancer, chicken β-actin promoter and an SV40 intron, and upstream of a WPRE and SV40 polyadenylation site (Extended Data Fig. 2). The reading frames for the IgG1 heavy and light chains were separated by a P2A ribosomal skip sequence for expression from the same mRNA transcript and a furin cleavage site for proteolytic removal of the P2A peptide. To minimize off-target expression in professional antigen-presenting cells, three tandem repeats of the miRNA binding site miRNA-142T were included in the 3’UTR^30,31^. Each of these antibodies contain heavy chain M428L and N434S (LS) substitutions to extend their *in vivo* half-lives through increased affinity for the neonatal Fc receptor^32,33^. A rhodopsin tag (C9) was also appended to the C-terminus of ITS61.01 and 17-HD9 for quantification by ELISA^25^.

### Prevention of SIV rebound after discontinuing ART

Fourteen adult rhesus macaques of Indian ancestry were selected for this study after pre-screening to exclude animals with neutralizing antibody titers to AAV serotype 9 (AAV9)^34^ and MHC class I alleles associated with spontaneous control of SIV replication (*Mamu-B*008* and *-B*017*^35,36^). All fourteen animals were infected intravenously with 5,000 IU of barcoded SIV_mac_239M, a dose sufficient to seed the rebound-competent viral reservoir (RCVR) with ∼100-1,000 unique barcodes^37–39^. On day 9 post-infection (PI), a daily subcutaneous ART regimen of tenofovir disoproxil fumarate, emtricitabine and dolutegravir was initiated and maintained until week 60 PI (Fig. 1a). At week 34 PI, when viremia was well controlled, eight animals in the treatment group received intramuscular injections of AAV9 packaged vectors encoding ITS61.01 and ITS103.01 (3.3×10^12^ gc/kg each) and six control animals received an AAV9 vector encoding 17-HD9. ART was maintained for an additional 26 weeks to allow antibody expression to stabilize. During this period, there was a general decline in cell-associated viral RNA and DNA in PBMCs and lymph nodes, consistent with a gradual reduction in the size of the viral reservoir (Extended Data Fig. 3). However, we did not observe a greater reduction or a more rapid rate of decline in the animals treated with ITS61.01 and ITS103.01 compared to the control animals, which would have been consistent with targeted depletion of the viral reservoir.

**Fig. 1.**
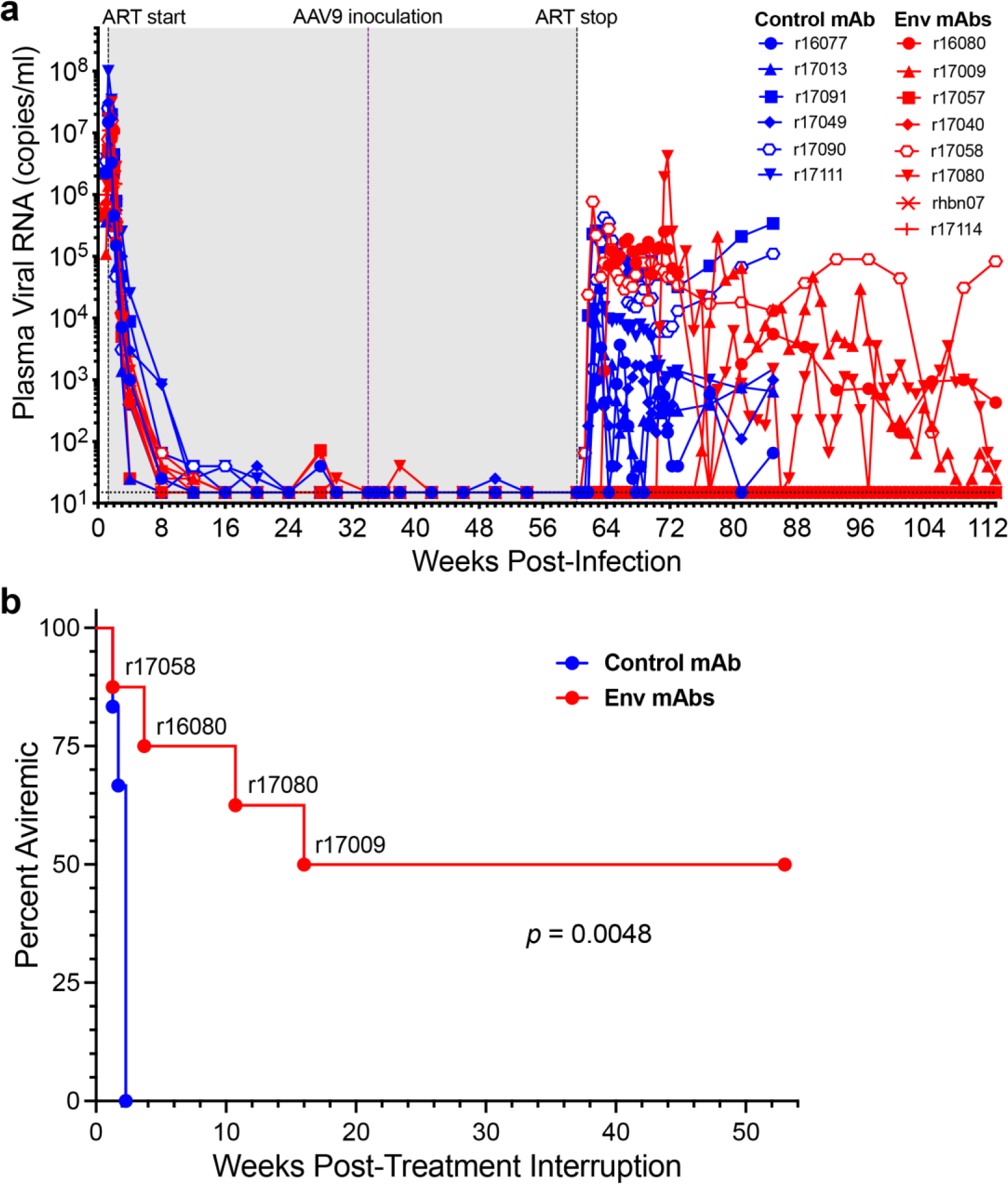
AAV-delivery of antibodies to the SIV envelope glycoprotein delays viral rebound in SIV_mac_239M-infected rhesus macaques. **a**, Fourteen rhesus macaques were infected intravenously with barcoded SIV_mac_239M (5,000 IU)^37^. From day 9 to week 60 PI, the animals were maintained on a daily subcutaneous ART regimen consisting of dolutegravir (2.5 mg/kg), tenofovir disoproxil fumarate (5.1 mg/kg) and emtricitabine (40 mg/kg). At week 34 PI, eight animals were inoculated with AAV9 vectors encoding the SIV Env-specific antibodies ITS61.01 and ITS103.01 (red), and six animals were inoculated with a vector encoding the control antibody 17-HD9 (blue). Viral RNA loads in plasma were measured using a qRT-PCR assay with a detection threshold of 15 copies/ml (dotted line)^47^. The shaded region indicates the period of ART (gray). **b**, The percentage of animals with viral loads below the threshold of detection (< 15 copies/ml) are shown after treatment interruption. Differences in the percentage of aviremic animals between the treatment and control groups were compared (p<0.005, Mantel-Cox test).

At week 60 PI, ART was discontinued and plasma SIV RNA was monitored every 2-3 days for 12 weeks, and weekly thereafter, for potential viral rebound. Plasma viral loads rebounded to >300 RNA copies/ml in all six of the control animals within two weeks of treatment interruption (TI) (Fig. 1a). Viral loads also rebounded within one week of TI in animal r17058, which had the highest ADA responses and lowest concentrations of ITS61.01 and ITS103.01 in serum (see Fig. 2). In three other animals that received ITS61.01 and ITS103.01, viral rebound was delayed for 3-17 weeks (Fig. 1a). However, plasma viral loads remained below the assay threshold (<15 RNA copies/ml) for more than a year after TI in four of the eight animals that received vectors expressing Env-specific antibodies (Fig. 1a). The difference in the rate of viral rebound between the treatment and control groups was significant (*p*=0.0048, Mantel-Cox test, Fig. 1b).

**Fig. 2.**
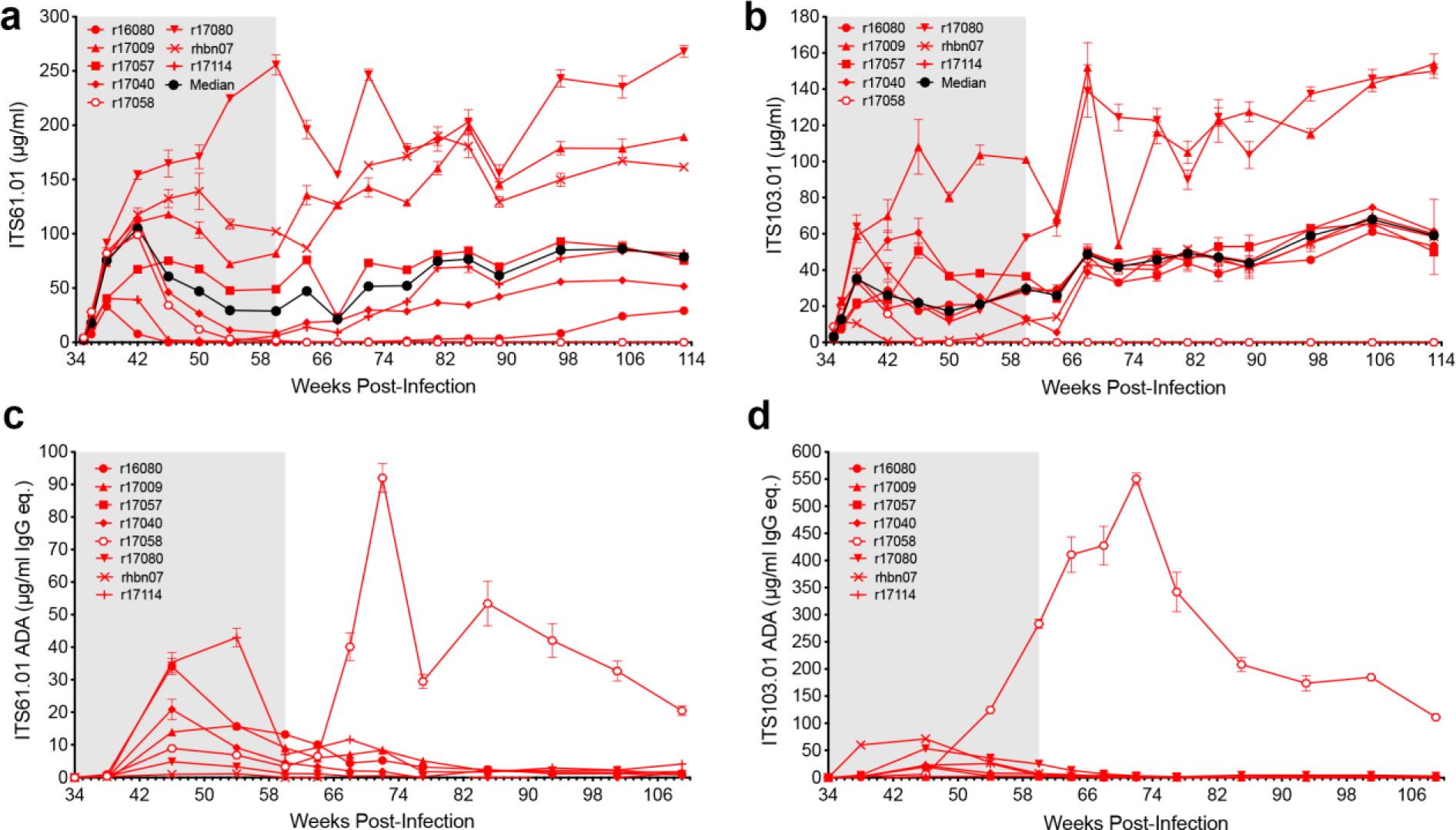
Serum antibody concentrations and anti-drug antibody responses. **a,** ITS61.01 and **b**, ITS103.01 concentrations in serum were measured by ELISA on plates coated with an antibody to the rhodopsin tag appended to the C-terminus of ITS61.01 or with an anti-idiotype antibody to ITS103.01. Anti-drug antibody responses to **c**, ITS61.01 and **d**, ITS103.01 were measured by probing ITS61.01- or ITS103.01-coated plates with biotinylated IgG purified from serum followed by streptavidin-HRP and development in TMB substrate.

### Env-specific antibody concentrations and anti-antibody responses

The serum concentrations of both Env-specific antibodies were highly variable. At the time of TI (week 60 PI), ITS61.01 ranged from 0.13 to 256 (median 28.8) µg/ml and ITS103.01 ranged from 0.1 to 101 (median 29.6) µg/ml. Both antibodies exhibited a trend toward increasing expression levels at later time points, with median serum concentrations rising to 78.8 µg/ml for ITS61.01 and to 59.1 µg/ml for ITS103.01 by week 113 PI (Fig. 2a, b and Extended Data Table 1).

ADA responses were low and generally reflected an inverse relationship with ITS61.01 and ITS103.01 levels. The only animal with runaway ADA responses (r17058) had the lowest concentrations of both antibodies in serum at the time of ART release (Fig. 2c, d and Extended Data Table 1). Thus, rapid viral rebound in this animal can be explained by negligible concentrations of the vectored antibodies as a result of strong ADA responses. Consistent with gradually increasing levels of both Env-specific antibodies, progressively waning ADA responses were observed in other animals. This stands in marked contrast to previous studies using AAV to deliver “simianized” versions of human bnAbs, which elicited strong ADA responses that severely limited bnAb expression in most animals^25,26^.

Serum samples from the treated animals were also tested for neutralization and ADCC shortly after ART release to assess antiviral antibody responses. Sera collected 24 days after TI were tested for the ability to neutralize SIV_mac_239 and to direct ADCC against SIV_mac_239-infected cells. All of the samples neutralized SIV_mac_239 with titers that generally corresponded to serum concentrations of ITS103.01 (Extended Data Fig. 4a). All of the samples also mediated ADCC against SIV_mac_239-infected cells (Extended Data Fig. 4b). To determine how much of these antiviral activities could be attributed to ITS103.01, we took advantage of the unusual sensitivity of this antibody to heat-inactivation in the presence of serum (Extended Data Fig. 4c, d). Heat-inactivation for 20 minutes at 55°C completely eliminated serum neutralization of SIV_mac_239 for all of the animals except r17058 (Extended Data Fig. 4e). However, a low level of ADCC reflecting the activity of ITS61.01 remained after heat-treatment (Extended Data Fig. 4f). Thus, except r17058, for which viral rebound two weeks earlier may have stimulated recall responses, endogenous neutralizing antibody titers were not detectable after suspending ART.

### Restricted barcode diversity of rebounding SIV

To determine if the vectored antibodies limited the diversity of SIV clonotypes contributing to viral rebound, the number of distinct barcodes identified in plasma at the time of ART initiation (day 9 PI) and immediately after viral rebound were analyzed. By day 9 PI, 92-770 (median 373) unique barcodes were identified in plasma (Fig. 3). At the earliest detectable time point after TI, there were 2-16 (median 3.5) barcodes identified in the plasma of control animals compared to 1-2 barcodes (median 1.0) in the rebounding animals that received vectors expressing Env-specific antibodies (p=0.019, Mann-Whitney U test) (Fig. 3 and Extended Data Table 2). In each case, the rebounding clonotypes were among the highest frequency barcodes in plasma prior to ART. At peak rebound, the number of detectable clonotypes in the control group increased to 5-30 (median 8). Excluding r17058, which had the highest ADA responses and the lowest serum concentrations of both ITS61.01 and ITS103.01, only 1-2 barcodes (median 1) were detected in the three animals with viral rebound despite high levels of Env-specific antibodies (p=0.012, Mann-Whitney U test) (Fig. 3 and Extended Data Table 2).

**Fig. 3.**
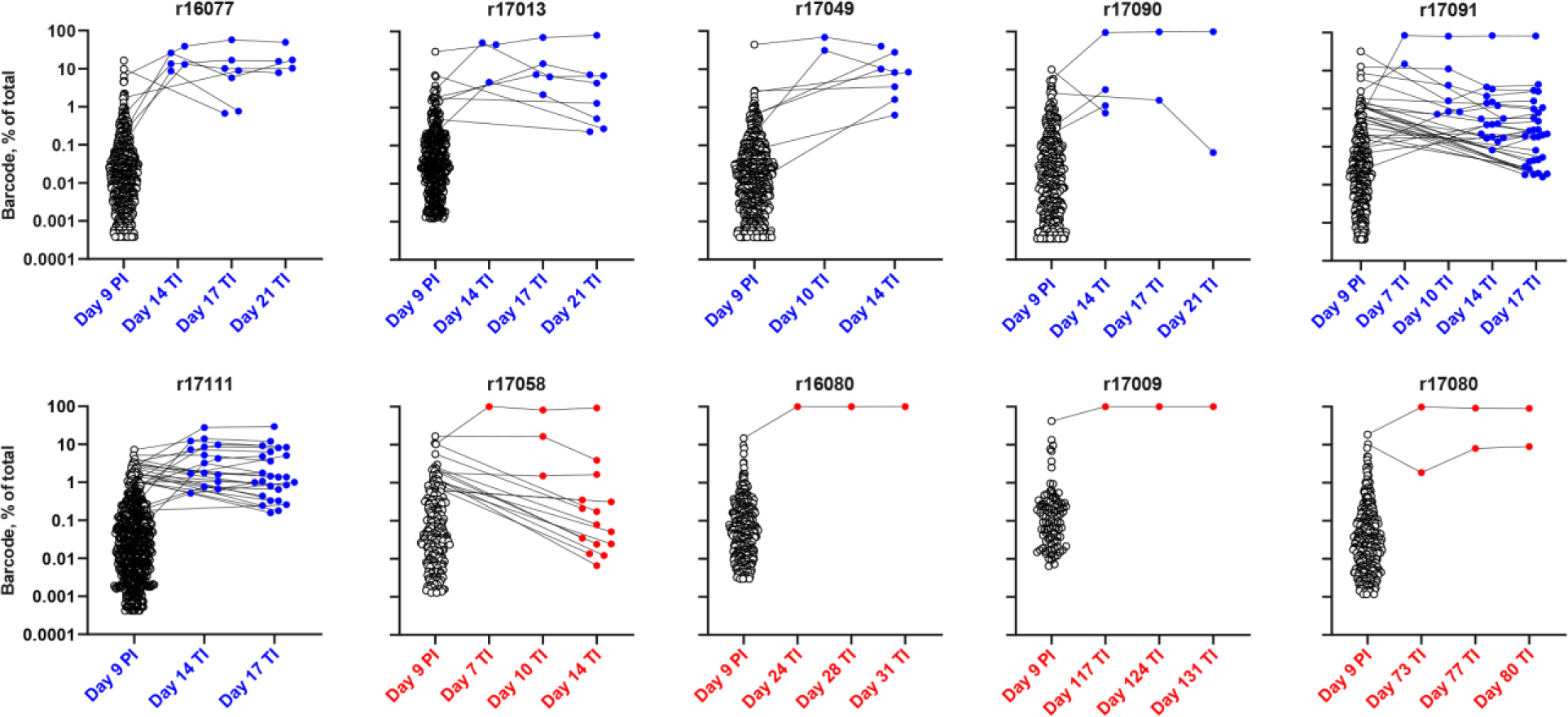
Restricted barcode diversity of the rebounding virus in animals with Env-specific antibodies. The virus population in plasma was sequenced on day 9 post-infection (PI) and at the indicated time points after treatment interruption (TI) to determine the number of unique SIV_mac_239M barcodes^37^. The frequency of each of the barcodes detected in the animals that received vectors expressing 17-HD9 (blue) or ITS61.01 and ITS103.01 (red) is shown.

Using the rebound growth rate and the relative proportion of identified barcodes in plasma, the average time between viral reactivation events (reactivation rate, RR) leading to rebound viremia was calculated^37–39^. In the control animals, the reactivation rate averaged 2.3 events per day (range 1.1-5.5, median 1.4) (Extended Data Table 2). Importantly, r17058 had 15 detectable clonotypes at rebound with a reactivation rate of 1.54, which is in line with the control group. Animals with delayed rebound or that failed to rebound within a year off ART were therefore exposed to an estimated 2.3 reactivation events per day (∼400-2,000 events per year). Together with the absence of viral rebound in four of the animals that received ITS61.01 and ITS103.01 more than one year after TI, these results indicate that AAV delivery of the Env-specific antibodies restricted the number of distinct SIV clonotypes emerging from the reservoir.

### Selection of antibody escape variants

To determine if the resurgence of SIV replication in animals that received AAV9 vectors encoding ITS61.01 and ITS103.01 was a result of antibody escape mutations in Env, the *env* gene of virus circulating in plasma was sequenced at two time points shortly after viral rebound. In addition to low frequency amino acid changes unique to each animal, three of the four animals acquired high frequency substitutions in Env exceeding 90% of the virus population, which are predicted to eliminate an N-linked glycan attached to residue N479 (Fig. 4a, Extended Data Fig. 5 and Supplemental Fig. 1). These include N479D, N479K and T481A in animals r16080, r17009 and r17080 (Fig. 4a). Additional substitutions predicted to eliminate another N-linked glycan on residue N295 (N295D and N297V) were observed in r16080 (Fig. 4a, Extended Data Fig. 5 and Supplemental Fig. 1). However, none of these changes occurred in r17058 (Fig. 4a), which had negligible serum concentrations of the vectored antibodies at the time of ART withdrawal. The N295 and N479 glycans are located on the periphery of the CD4 binding site (Fig. 4b)^40,41^, consistent with their participation in ITS103.01 binding to this region^42^.

**Fig. 4.**
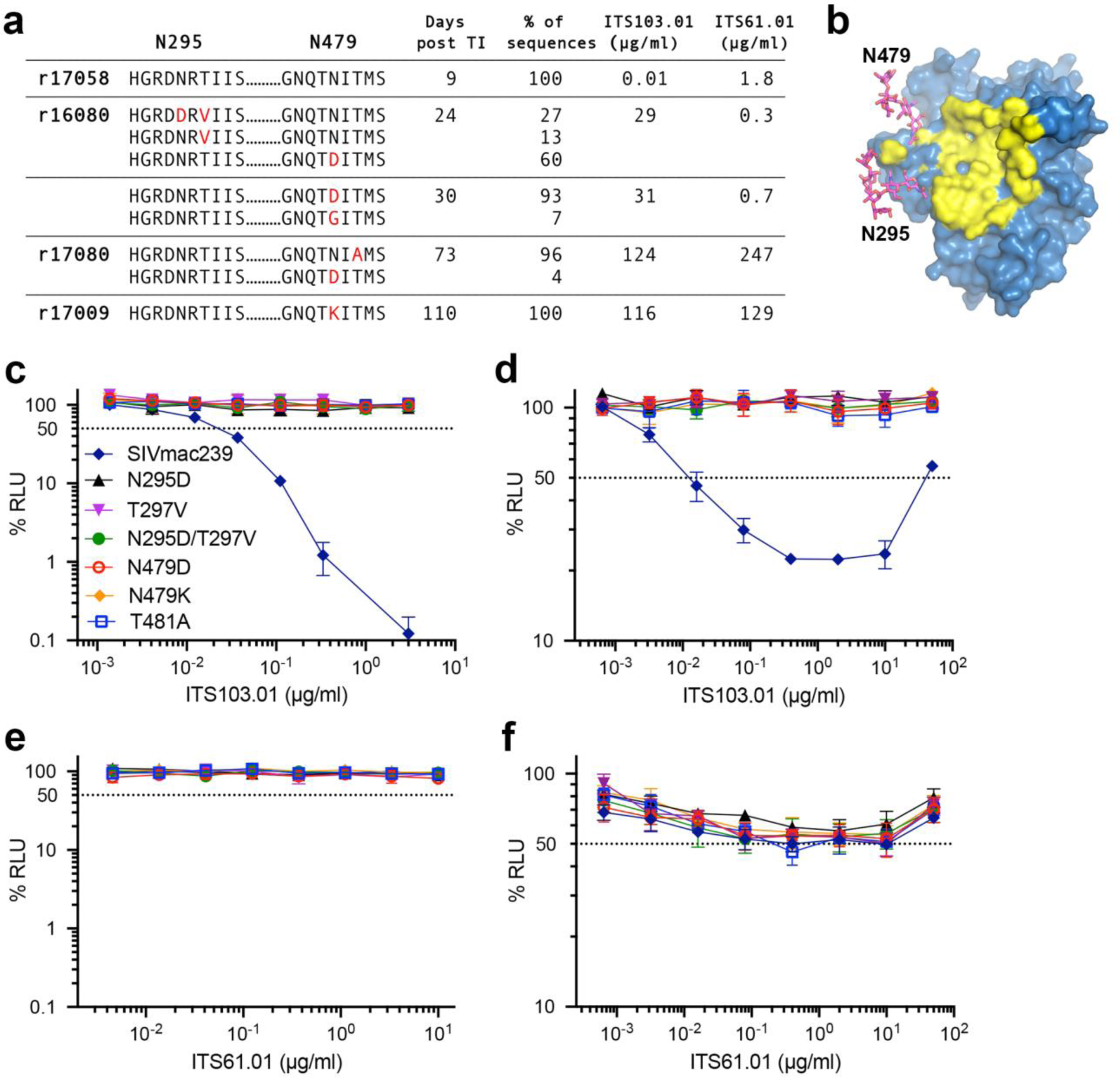
Substitutions predicted to eliminate glycosylation of Env residues N295 and N479 confer resistance to ITS103.01. **a**, SIV RNA was isolated from plasma after viral rebound and sequenced to look for evidence of antibody escape. Amino acid changes in Env (red) surrounding residues N295 and N479, their frequency, and serum concentrations of ITS103.01 and ITS61.01 are shown for each animal at the indicated time points after treatment interruption. **b**, N-linked glycans attached to N295 and N479 (magenta) are located on the periphery of the CD4-binding site (yellow)^40,41^. **c-f**, The introduction of substitutions into SIV_mac_239 predicted to abrogate glycosylation of N295 or N479 confer resistance to **c**, neutralization and **d**, ADCC by ITS103.01, but not does not alter sensitivity to **e**, neutralization or **f**, ADCC by ITS61.01.

Accordingly, the introduction of each of the Env substitutions into SIV_mac_239 predicted to prevent glycosylation of N295 or N479 provided complete resistance to ITS103.01 neutralization and ADCC (Fig. 4c, d) but did not confer sensitivity to neutralization by ITS61.01 (Fig. 4e) or alter the susceptibility of infected cells to ADCC in the presence of this antibody (Fig. 4f). Viral rebound despite high levels of vectored antibodies can therefore be explained by the emergence of antibody escape variants that confer resistance to ITS103.01.

## Discussion

A single round of intramuscular injection with AAV9-packaged vectors encoding natural rhesus macaque antibodies to the SIV envelope glycoprotein resulted in sustained antibody expression with no signs of abatement for more than one-and-a-half years. After discontinuing ART, SIV_mac_239M replication was maintained below the threshold of detection for over a year in 50% of the animals that received these vectors. SIV_mac_239 is considered to be a tier 3, neutralization-resistant virus with especially dense Env glycosylation that makes it highly refractory to host antibody responses^40^. Moreover, the viral reservoir in SIV-infected macaques is estimated to be an order of magnitude higher than in HIV-1-infected people^43^. This study therefore provides a rigorous demonstration of the potential for AAV-delivery of antibodies to contain viral rebound from an established infection after discontinuing ART.

In contrast to AAV-delivery of “simianized” versions of human bnAbs, which typically elicit strong ADA responses that severely impair expression in most animals^25,26^, ADA responses to rhesus macaque antibodies were generally low. Only one animal (r17058) developed high ADA titers that interfered with the expression of both Env-specific antibodies. ADA responses were also detectable in other animals for a few weeks after AAV9 administration. However, after an initial peak these responses progressively declined. The explanation for this decline is presently unclear but may reflect a tolerizing effect of continuous antibody expression reminiscent of the high zone tolerance that can occur with frequent or high dose administration of certain biologics^44–46^. Thus, although the delivery of natural rhesus macaque antibodies did not completely eliminate ADA, in most cases these responses were low, transient, and did not prevent vectored antibodies from reaching effective concentrations *in vivo*. This suggests that ADA may not be an insurmountable barrier to the delivery of natural, species-matched antibodies, which portends well for the potential identification of HIV-1 bnAbs for AAV-delivery in humans.

Env substitutions that confer resistance to ITS103.01 emerged in each of the animals with delayed viral rebound despite detectable levels of this antibody in serum. However, there was no evidence of escape from ITS61.01, which suggests that ITS103.01 was largely, if not entirely, responsible for containing virus replication after treatment interruption. ITS103.01 mediates potent neutralization and ADCC against SIV_mac_239, whereas ITS61.01 only mediates ADCC against this virus. Although the relative contribution of ADCC and other Fc-mediated effector functions to virus control is uncertain, these results suggest that potent neutralization is a critical determinant of antibody-mediated containment of virus replication.

Although cell-associated viral loads in peripheral blood and lymph nodes gradually declined while the animals were maintained on ART, we did not detect a greater reduction or a more rapid rate of decline in the animals that received Env-specific antibodies than in the control animals. Based on the reactivation rate in the control animals, we can infer that the treated animals needed to contain 1-5 reactivation events per day after stopping ART. At this continuous rate of challenge, the vectored antibodies were sufficient to block viral rebound in half of the treated animals and to delay viral rebound in the remaining treated animals with eventual loss of virus suppression due to the emergence of antibody-escape variants. These data are consistent with the hypothesis that ITS61.01 and ITS103.01 did not significantly reduce the rebound-competent viral reservoir. Rather, the antiviral activity of these antibodies prevented or delayed viral rebound by targeting spontaneously reactivating cells and/or by blocking virus transmission to uninfected cells.

The timing of ART initiation for this study was selected on the basis of extensive data indicating that starting ART at this time allows for the establishment of SIV_mac_239M infection with near peak seeding of the rebound-competent viral reservoir with 10^2^-10^3^ distinct barcode variants and universal post-ART rebound^39^. Due to early initiation of ART, it is unlikely that the viral reservoir contained a high frequency of antibody-escape mutations. Rather, antibody-escape likely occurred after the reactivation of latently infected cells but prior to the detection of virus in plasma. Considering the ability of rebounding virus to generate escape mutations in this model, reliable, long-term suppression of HIV-1 in a majority of individuals will probably require the delivery of two or more bnAbs, which the present results suggest should be feasible.

The unprecedented containment of highly pathogenic SIV in half of the treated animals in this study for more than a year after discontinuing ART provides a rigorous demonstration of the potential for AAV-delivery of antibodies to achieve durable control of virus replication. A key to the success of this approach was the delivery of natural, species-matched antibodies that elicit minimal ADA responses allowing antibody expression to be maintained indefinitely at effective concentrations after a single round of AAV administration. Dozens of potent, broadly active neutralizing antibodies targeting diverse epitopes of the HIV-1 Env trimer are now available. This raises the intriguing possibility that similar ART-free control of HIV-1 may be possible through the judicious selection of some of these bnAbs for AAV-delivery.

## Methods

### Ethics statement

Rhesus macaques (*Macaca mulatta*) were housed at the Wisconsin National Primate Research Center (WNPRC) in accordance with the standards of AAALAC International and the University of Wisconsin Research Animal Resources Center and Compliance unit (UWRARC). Animal experiments were approved by the University of Wisconsin College of Letters and Sciences and the Vice Chancellor for Research and Graduate Education Centers Institutional Animal Care and Use Committee (protocol number G005141) and performed in compliance with the principles described in the Guide for the Care and Use of Laboratory Animals^48^. Fresh water was always available, commercial monkey chow was provided twice a day, and fresh produce was supplied daily. To minimize any distress ketamine HCL alone or ketamine HCL in combination with dexamedetomidine were used to sedate animals prior to experimental procedures (e.g., blood collection, SIV infection, lymph node biopsy) and animals were monitored twice a day by animal care and veterinary staff. Analgesics and anti-inflammatories (e.g., buprenorphine, lidocaine, meloxicam) were administered to alleviate pain associated with the experimental procedures. The animals were socially housed in pairs or groups of compatible animals whenever possible.

### Animals

Fourteen rhesus macaques of Indian ancestry and free of simian retrovirus type D (SRV), simian T-lymphotropic virus type 1 (STLV-1), SIV and macacine herpesvirus 1 (herpesvirus B) were used for these studies. Animals were pre-screened for AAV9 neutralizing antibodies and were considered negative with no detectable neutralizing activity at a 1:10 dilution of serum^34^. MHC class I genotyping was performed using genomic DNA isolated from PBMCs by sequencing a 150 bp region of exon 2 (Illumina MiSeq system). Sequences were analyzed by comparison to an inhouse database as previously described^49^. Animals positive for AAV9 neutralizing antibodies or MHC class I alleles associated with spontaneous control of SIV infection (*Mamu-B*008* and *-B*017*^35,36^) were excluded and experimental and control groups were structured with a similar distribution of male and female animals. The sex, age, weight and MHC class I genotype of the animals are shown in table S3.

### AAV vector design and preparation

The overall design of the AAV transfer plasmids containing AAV2 inverted terminal repeats was similar to previous reports^21,34^. Codon-optimized (Genscript) sequences for the heavy and light chains of the SIV Env-specific rhesus macaque antibodies ITS61.01 and ITS103.01^27,42^ and the RSV F-specific control antibody 17-HD9^28^ were cloned into an AAV vector downstream of a CMV enhancer, chicken β-actin promoter and an SV40 intron and upstream of a WPRE and SV40 polyadenylation site (Extended Data Fig. 2). The heavy and light chain reading frames were separated by a P2A ribosomal skip sequence and a furin cleavage site for proteolytic removal of the P2A peptide. Three tandem repeats of the miRNA binding site miRNA-142T were included in the 3’ UTR to minimize off-target expression in professional antigen presenting cells^30,31^. M428L and N434S (LS) substitutions were introduced into the heavy chain of each antibody to extend their *in vivo* half-lives through increased affinity for the neonatal Fc receptor^32,33^. The 17-HD9 construct lacked the miRNA-142 binding sites and contained S239D, A330L and I332E (DLE) substitutions to enhance binding to Fcγ receptors^50^.

Recombinant AAV9 vectors were produced as described previously^51,52^. HEK293T cells were co-transfected with the AAV transfer plasmid encoding antibody, the pAAV2/9n packaging plasmid expression Rep2/Cap9 (Addgene, cat. 112865) and the pAdDeltaF6 plasmid expressing adenoviral helper proteins E2a, E4 and VA (Addgene, cat. 112867). AAV9 particles were purified from lysates of transfected cells on CsCl density gradients followed by extensive dialysis against buffer consisting of 5% sorbitol in PBS, after which Pluronic F-68 was added to 0.001% final concentration. Vector concentration in genome copies per ml (gc/ml) was determined by droplet digital PCR for DNAse-resistant templates^53^. Purity of the final AAV9 preparations was evaluated from a silver-stained SDS-PAGE. For long-term storage, 0.5 ml aliquots of the AAV9 stocks were frozen and stored at -80°C.

### SIV infection and antiretroviral therapy

The barcoded SIV_mac_239M used for this study is the same virus stock described previously^37^. After dilution of this virus to 5,000 IU/ml in RPMI 640 medium without serum, all fourteen animals were infected by intravenous inoculation with 1 ml of the virus preparation.

A solution of the antiretroviral drugs dolutegravir (DTG, 2.5 mg/ml), tenofovir disoproxil fumarate (TDF, 5.1 mg/ml) and emtricitabine (FTC, 40 mg/ml) was prepared in 15% w/v Kleptose (HPB Biopharma, cat. 346113105E), filter-sterilized through a 0.2 μm nylon membrane and stored at -20°C. Beginning on day 9 PI and ending on week 60 PI, each animal received daily subcutaneous injections of ART at doses of 1 ml per kg body weight^54^.

### AAV inoculations

Eight rhesus macaques received AAV9 vectors encoding the SIV Env-specific antibodies ITS61.01 and ITS103.01 and six control animals received an AAV9 vector encoding the RSV F-specific antibody 17-HD9. The AAV9 vectors were administered by intramuscular injection at doses of 3.3×10^12^ gc/kg for ITS61.01 and ITS103.01 and 2.7×10^12^ gc/kg for 17-HD9. To prevent heavy and light chain mixing in coinfected cells, vectors encoding ITS61.01 and ITS103.01 were given on opposite sides of the body. To maximize muscle transduction, the AAV9 inoculum was distributed over six separate injection sites on the same side of the body. Each dose of a given vector was suspended in 3 ml of sterile PBS, and 0.5 ml volumes drawn into tuberculin syringes (Excelint International, cat. 26046) were injected into two sites in the quadriceps, two in the deltoids and two in the biceps.

### SIV viral detection assays

Plasma SIV RNA levels were determined using a *gag*-targeted quantitative real time/digital RT-PCR format assay, essentially as previously described with 6 replicate reactions analyzed per extracted sample for an assay threshold of 15 SIV RNA copies/ml^47^. Samples that did not yield any positive results across the replicate reactions were reported as a value of “less than” the value that would apply for one positive reaction out of six^47^. Quantitative assessment of SIV DNA and RNA in cells and tissues was performed using *gag*-targeted nested quantitative hybrid real-time/digital RT-PCR and PCR assays, as previously described^55,56^. SIV RNA or DNA copy numbers were normalized based on quantitation of a single copy rhesus genomic DNA sequence from the *CCR5* locus from the same specimen to allow normalization of SIV RNA or DNA copy numbers per 10^6^ diploid genome cell equivalents, as described^57^. Ten replicate reactions were performed with aliquots of extracted DNA or RNA from each sample, with two additional spiked internal control reactions performed with each sample to assess potential reaction inhibition. Samples that did not yield any positive results across the replicate reactions were reported as a value of “less than” the value that would apply for one positive reaction out of 10. Threshold sensitivities for individual specimens varied as a function of the number of cells or amount of tissue available and analyzed.

### Sequencing

Barcode sequencing was performed as previously described^37,58^. Briefly, total RNA was isolated from plasma using the QIAamp Viral RNA mini kit (Qiagen, cat. 52904) or QIASymphony DSP virus/pathogen Midi kit (Qiagen, cat. 937055) on a QIASymphony SP laboratory automation instrument platform according to manufacturer’s instructions. Complementary DNA (cDNA) was generated with Superscript III reverse transcriptase (ThermoFisher, cat. 18080093) and an SIV-specific reverse primer (Vpr.cDNA3: 5′-CAG GTT GGC CGA TTC TGG AGT GGA TGC). The cDNA was quantified via qRT-PCR using the primers VpxF1 5′-CTA GGG GAA GGA CAT GGG GCA GG and VprR1 5′-CCA GAA CCT CCA CTA CCC ATT CATC with labeled probe (ACC TCC AGA AAA TGA AGG ACC ACA AAG GG). Prior to sequencing, PCR was performed with VpxF1 and VprR1primers combined with either the F5 or F7 Illumina adaptors containing unique 8-nucleotide index sequences for multiplexing. PCR was performed using High Fidelity Platinum Taq (ThermoFisher, cat. 11304011) under the following conditions: 94°C, 2 min; 40 cycles of (94°C, 15 s; 60°C, 1.5 min; 68°C, 30 s); 68°C, 5 min. The multiplexed samples and Phi X 174 library were prepared via standard protocols and sequenced on a MiSeq instrument (Illumina).

Single genome analysis (SGA) amplification and Sanger sequencing of the SIV *env* gene was performed as described previously^59^.

### Reactivation rate estimation

The viral reactivation rate is defined as the viral growth rate over the average log-ratio of the number of copies of the distinct barcodes detected in rebound plasma. The reactivation rate was estimated using a previously developed mathematical model based on the ratio of viral growth rate to viral loads of different barcoded clonotypes as previously described^37^.

### ELISA for monitoring monoclonal antibody concentration in serum

ITS61.01 with a C-terminal rhodopsin tag (C9 tag) and untagged ITS103.01 were produced in transfected Expi293 cells (ThermoFisher, cat. A14527) and purified on rProtein A Gravitrap columns (Cytiva Life Sciences, cat. 28985254) according to manufacturer’s suggestions. These antibodies were used to build a calibration curve in a standard sandwich ELISA at final concentrations ranging from 0.47 to 15 ng/ml.

Reacti-Bind plates (Thermo Scientific, cat. 15041) were coated overnight at 4°C with either 0.75 μg/ml anti-rhodopsin antibody (clone 1D4, MilliporeSigma, cat. MAB5356) or 1.1 μg/ml mouse ITS103.01 anti-idiotype antibody in PBS^42^. All washes were done with PBS plus 0.07% Tween 20 (PBST) and all incubation steps (blocking, primary and secondary antibody) used PBST containing 5% non-fat dry milk. Experimental sera samples were not heat-inactivated and were applied to the plate as serial two-fold dilutions. Blocking and primary sample incubations were one hour at 37°C and secondary antibody incubations with a 1:6,000 dilution of mouse anti-monkey HRP conjugate (clone SB108a, Southern Biotech, cat. 4700-05) were 45 min at 37°C. Bound HRP activity was developed with SureBlue TMB substrate (LGC Clinical Diagnostics, cat. 5120-0075) for 10-15 min, the reaction was stopped with an equal volume of 1 M H_2_S0_4_ and absorbance at 450 nm was quantified on a Victor X4 plate reader (PerkinElmer). Values reported are A_450_ means ± standard deviations across all dilutions falling within a valid calibration curve range that are at least two-fold above background and no more than 2.8.

### Anti-drug antibody assay

Anti-drug antibodies were measured by microscale on-column purification and biotinylation of total IgG from rhesus macaque serum followed by quantification of the amount of biotinylated IgG bound to immobilized ITS61.01 or ITS103.01. Heat-inactivated sera (250 µl) were mixed in a 1.5 ml microcentrifuge tube with 100 μl of Binding Buffer (460 mM K_2_HPO_4_, 40 mM NaH_2_PO, 2 M NaCl), 650 μl water and 50 μl of 50% suspension of rProtein A Sepharose Fast Flow (Cytiva Life Sciences, cat. 17127902) equilibrated in PBS with 6 mM sodium azide. These mixtures were incubated at room temperature with end-to-end rotation for one hour. After washing once with 1 ml of Wash Buffer (WB, consisting of 23 mM K_2_HPO_4_, 2 mM NaH_2_PO_4_, 200 mM NaCl), the Sepharose pellets were mixed with 1 ml WB, transferred to 1 ml columns with pre-wetted bottom frit (Bio-Rad, cat. 7326008) and further washed three times with 1 ml WB by gravity flow. The columns were then inserted into 5 ml tissue culture tubes and were spun at 60 *g* momentarily to remove excess liquid without drying the sorbent, and outlets were then capped with rubber caps (BD, cat. 308341). Working as quickly as possible with batches of four columns to minimize non-productive hydrolysis of NHS group, 100 μl of freshly prepared 0.2 mg/ml of ChromaLINK biotin (Vector Laboratories, cat. B-1001-105) in WB was added, mixed with Sepharose by tapping the column, capped to prevent evaporation, inserted into a 5 ml tissue culture tube and rocked vertically at maximum speed for one hour at room temperature. Typically, an aliquot of 1 mg of ChromaLINK biotin was solubilized in 50 μl of anhydrous DMSO and either used immediately or stored in a screw cap tube at -80°C. A working concentration of 0.2 mg/ml reagent was prepared in batches of 450 μl by diluting the DMSO stock solution 100-fold with WB. After incubation, the columns were washed three times with 1 ml WB by gravity flow, flicked by hand to remove excess liquid from the tip. Bound IgG was eluted with two subsequent additions of 100 μl of elution buffer to the column and placed into 5 ml culture tube containing 20 μl of sodium bicarbonate pH 9.3. The extent of biotinylation was quantified by absorbance at 280 and 354 nm on a Nanodrop 2000 as suggested by the ChromaLINK manufacturer’s instructions, which assume IgG average molecular mass of 150 kDa, absorbance at 280 nm of 1 mg/ml IgG equals 1.38 optical units, absorbance of 1 mM ChromaLINK biotin at 354 nm equals 29 optical units and absorbance of 1 mM ChromaLINK biotin at 280 nm equals 6.67 optical units. Under the described conditions, the extent of biotinylation typically ranged from 1.7 to 2.0 biotins per IgG molecule. All samples were subsequently diluted to 1 mg/ml IgG with PBS containing 10 mg/ml bovine serum albumin (BSA; Fisher Scientific, cat. BP1605-100), aliquoted and stored at -80°C. For a known negative standard, a sample consisting of equal volumes of pre-immune sera from all experimental animals was used. For calibration standard, 40 μg/ml polyclonal rabbit anti-mouse antibody (Jackson ImmunoResearch, cat. 315-005-045) was spiked into pre-immune serum. Both standards were processed identically to the individual experimental samples. Up to 16 samples can be conveniently processed in parallel by this approach.

ELISA conditions for the detection of biotinylated IgG binding to immobilized monoclonal antibodies were as follows. Wells of Reacti-Bind plates (Thermo Scientific, cat. 15041) were coated at 4°C overnight with 2 μg/ml of ITS61.01, ITS103.01 or polyclonal mouse IgG (Jackson ImmunoResearch, cat. 015-000-002). All plate washes were done with ADA wash buffer (ADA-WB) consisting of 25 mM Tris, 140 mM NaCl, 0.5% Tween 20, pH 8.2. Coated plates were blocked with 10 mg/ml BSA in ADA-WB for one hour at 37°C. Biotinylated samples were serially diluted two-fold with 20% FBS (VWR, cat. 76419-584) in ADA-WB starting at 10 μg/ml, added to the wells and incubated for one hour at 37°C. Biotinylated IgG retained on plates was reacted with a 1:10,000 dilution Streptavidin-HRP conjugate (Thermo Scientific, cat. N504) in ADA-WB containing 10 mg/ml BSA for one hour at 37°C. HRP activity was developed with SureBlue TMB substrate for 30 min as described for above. There is an appreciable background observed in this assay that stems from minute amounts of some non-specific IgG-binding. The negative control values were therefore subtracted from the corresponding experimental and calibration values before further analysis. The colorimetric signals were converted to the equivalents of polyclonal IgG concentrations using the calibration curve obtained with the biotinylated rabbit anti-mouse IgG standard assayed against total mouse IgG. Reported values are means and standard deviations of all dilutions falling within a valid calibration curve range.

### Neutralization assay

SIV neutralization was measured as described using a standard TZM-bl assay^60^ with a few modifications. Serum samples, without heat-inactivation were serially diluted in D10 medium (DMEM containing 10% heat-inactivated FBS) and incubated with SIV_mac_239Δ*vif* in white 96-well plates (Greiner, cat. 82050-736) at 6 ng of SIV p27 per well in a final volume of 100 μl per well. After one hour at 37°C, 10,000 of TZM-bl cells (NIH HIV Reagent Program, cat. ARP-8129) were added in 100 μl of D10. After 72 hours, the medium was carefully aspirated and 100 μl of BriteLite Plus luciferase substrate diluted two-fold in PBS (Revvity, cat. 6066761) was added to each well. The relative light units (RLU) of luciferase activity in each well was measured within 5 min after the addition of substrate using a Victor X4 plate reader (PerkinElmer). After subtracting background luciferase activity in uninfected cells, the luciferase activity was normalized to 100% RLU relative to wells with infected cells without serum or neutralizing antibody. All reported values are means and standard deviations of technical triplicates.

### ADCC assay

ADCC was measured as previously described^61^ with the following modifications. Two to five million CEM.NKR-_CCR5_-sLTR-Luc target cells were infected with VSV-G pseudotyped SIVmac239Δ*vif* (125-250 ng p27) in 15 ml conical tubes by spinoculation at 1200 *g* in the presence of 40 μg/ml Polybrene for 90 min at 25°C. Three days after infection, the target cells were washed three times with R10 (RPMI 640 medium supplemented with 10% heat-inactivated FBS) and resuspended in assay medium (R10 containing 10 U/ml IL-2). CEM.NKR-_CCR5_-sLTR-Luc target cells (1×10^4^ cells) were mixed with an NK cell line (KHYG-1 cells) transduced with rhesus macaque CD16 (1×10^5^ cells) in the presence of serial dilutions of serum in a round-bottom 96-well plates (200 μl/well). After an 8-hour incubation at 37°C, 150 μl of the cell suspension from each well was transferred to white 96-well plates (Greiner, cat. 82050-736) containing 50 μl per well of BriteLite Plus luciferase substrate. The luciferase activity (RLU) in each well was measured using a Victor X4 plate reader. After correction for the background luciferase activity in wells containing uninfected cells without serum, the measured the luciferase activity was normalized to 100% RLU relative to wells with infected cells but without serum. All reported values are means and standard deviations of technical triplicates.

### Statistics and data presentation

All statistical comparisons were done with GraphPad Prism version 8.4.3.

## Acknowledgements

We thank ViiV Healthcare Ltd. for providing dolutegravir and Romas Geleziunas, Gilead Sciences Inc., for providing tenofovir disoproxil fumarate and emtricitabine for antiretroviral treatment of the animals in this study. This work was supported in part by the intramural Research Program of the National Institute of Allergy and Infectious Diseases. The content of this publication does not necessarily reflect the views or policies of the Department of Health and Human Services, nor does mention of trade names, commercial products, or organizations imply endorsement by the U.S. Government.

## Author contributions

Conceptualization: JDL, MR, MRG, DTE

Methodology: VAK, BFK

Formal analysis: VAK, AB

Investigation: VAK, NMC, NKK, CMF, BFK

Resources: RM, GG, MR, MRG

Writing – original draft: VAK, DTE

Supervision: SC, BFK, JDL, DTE

Funding acquisition: DTE

## Data and materials availability

All data are available in the main text or the supplementary materials. Nucleotide sequences of SIV *env* variants are available at GenBank accession numbers PP098761 - PP098961. Nucleotide sequences of the AAV vectors encoding antibodies ITS61.01, 17-HD9 and ITS103.01 are available at GenBank accession numbers PP481455, PP481456 and PP498839, respectively.

---

**Extended Data Fig. 1.**
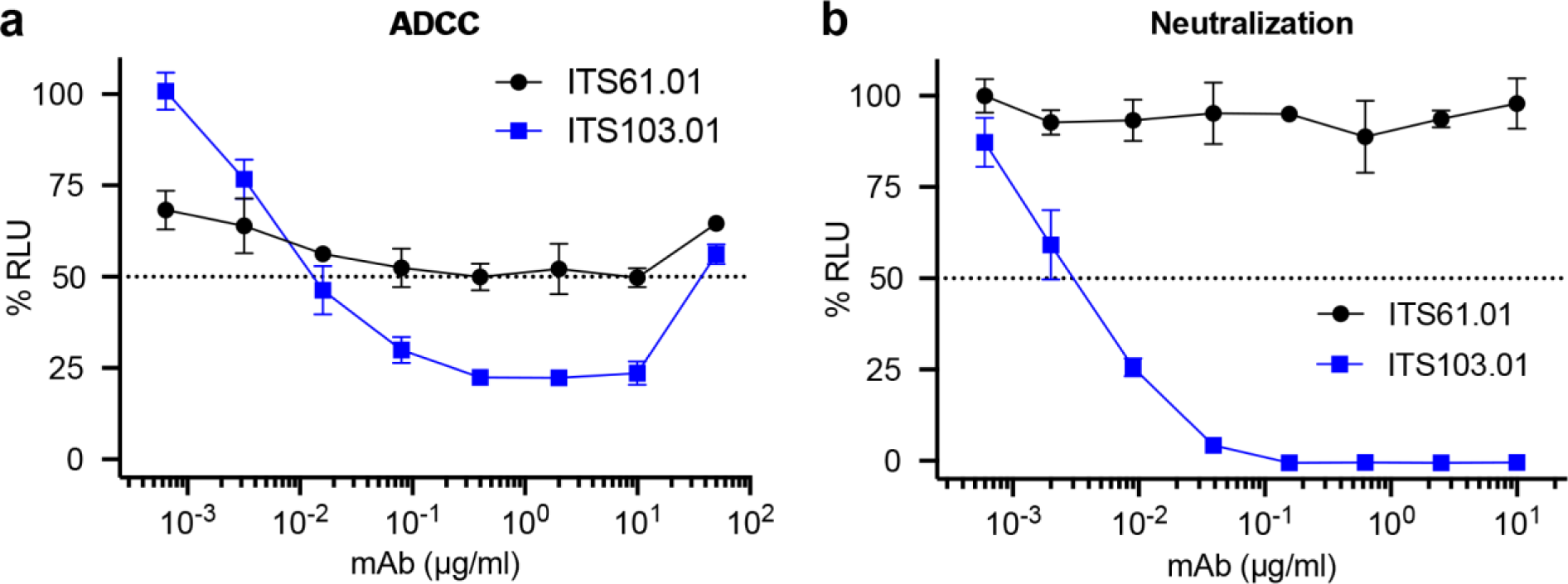
ADCC and neutralization activity of SIV Env-specific antibodies. The SIV Env-specific antibodies ITS61.01 and ITS103.01 were tested for **a**, ADCC and **b**, neutralization activity against SIV_mac_239. **a**, ADCC responses were measured by infecting CEM.NKR-_CCR5_-sLTR-Luc cells with SIV_mac_239 and incubating with an NK cell line (KHYG-1 cells) expressing rhesus macaque CD16 in the presence of the indicated concentrations of antibody^61^. ADCC responses were calculated as the remaining luciferase activity (% RLU) after an 8-hour incubation at a 10:1 effector-to-target ratio. The values indicate the mean and standard deviation (error bars) for triplicate wells at each antibody concentration and the dotted line indicates half-maximal killing of SIV-infected cells. **b**, SIV neutralization was measured by incubating virus dilutions with the indicated antibody concentrations for one hour before addition to TZM-bl cells. Three days post-infection, neutralization was calculated as the reduction of luciferase activity (% RLU) in cells inoculated with virus plus antibody relative to cells inoculated with virus without antibody. The values indicate the mean and standard deviation (error bars) for triplicate wells and the dotted line indicates 50% neutralization.

**Extended Data Fig. 2.**
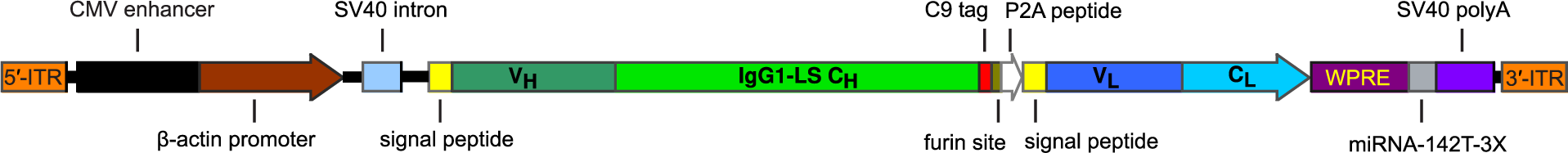
Adeno-associated virus vector for antibody expression. ITS61.01, ITS103.01 and 17-HD9 sequences were cloned into an AAV vector downstream of a CMV enhancer, chicken β-actin promoter and an SV40 intron and upstream of a WPRE and SV40 polyadenylation site. The heavy and light chain reading frames were separated by a P2A ribosomal skip sequence and a furin cleavage site for proteolytic removal of the P2A peptide. Three tandem repeats of the miRNA binding site miRNA-142T were included in the 3’UTR to minimize off-target expression in professional antigen presenting cells^30,31^. M428L and N434S (LS) substitutions were introduced into the heavy chain of each antibody to extend their *in vivo* half-lives through increased affinity for the neonatal Fc receptor^32,33^. A rhodopsin (C9) tag was also appended to the C-terminus of ITS61.01 and 17-HD9 for quantification by ELISA^25^.

**Extended Data Fig. 3.**
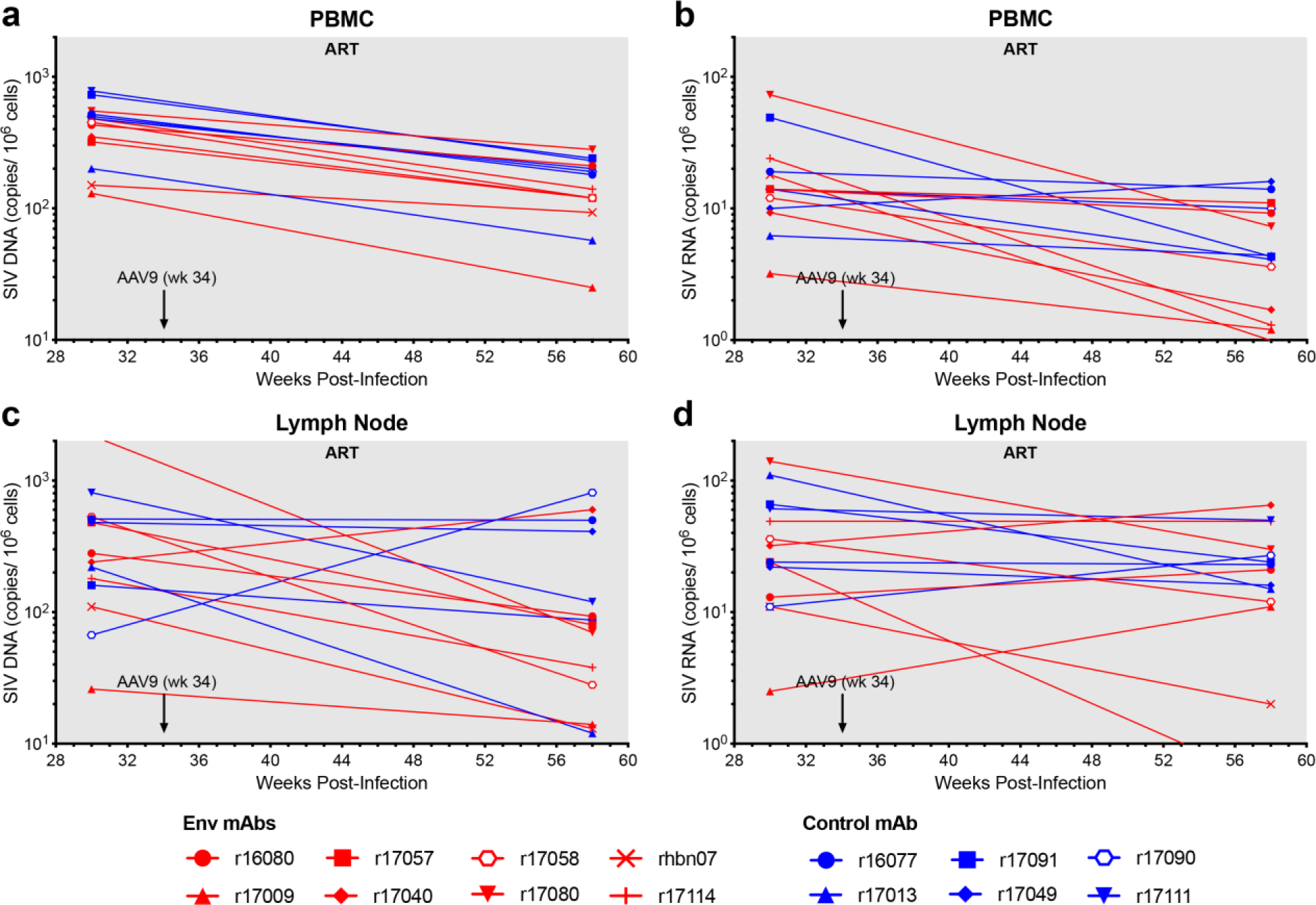
Cell-associated SIV loads in PBMCs and lymph nodes. **a**, Viral DNA in PBMCs **b**, viral RNA in PBMCs **c,** viral DNA in lymph nodes and **d**, viral RNA in lymph nodes were measured before (week 30 PI) and after (week 58 PI) the administration of AAV9 vectors to assess the impact of Env-specific antibodies on the size of the viral reservoir. Cell-associated SIV DNA and RNA were measured by quantitative hybrid real-time/digital RT-PCR and PCR assays as previously described^55^.

**Extended Data Fig. 4.**
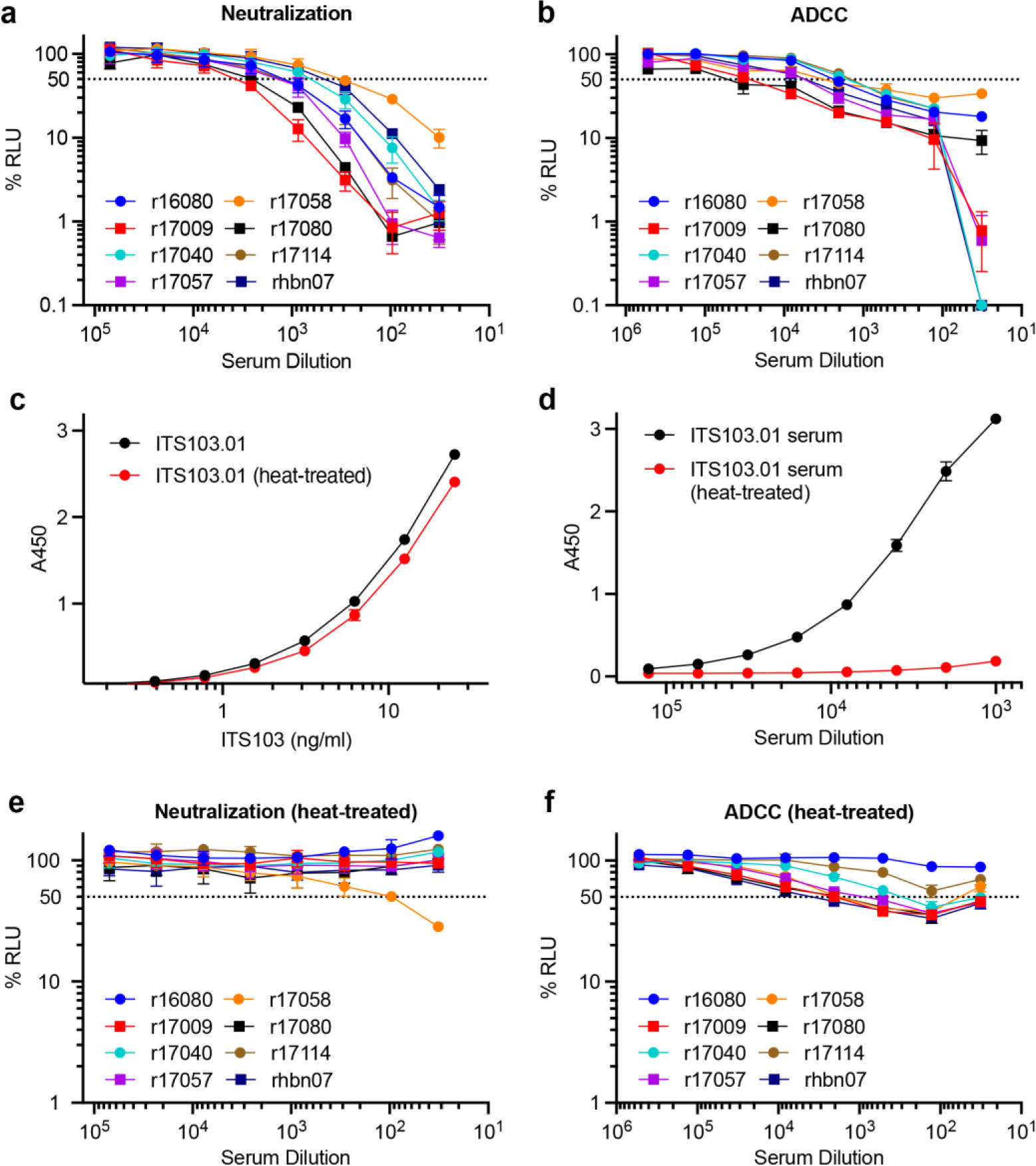
Neutralization and ADCC responses after treatment interruption. a, Neutralization and **b**, ADCC responses to SIV_mac_239 were measured in serum collected 4 weeks after TI (week 64 PI). The binding of **c**, purified ITS103.01 in PBS-BSA and **d**, ITS103.01 in serum from an AAV9-ITS103.01-inoculated animal to ELISA plates coated with a mouse anti-ITS103.01 idiotype antibody was determined with and without heat treatment at 55°C for 20 minutes. **e**, Neutralization and **f**, ADCC were measured after heat treatment of serum for 20 minutes at 55°C. Neutralization was assessed by the ability to block infection of TZM-bl cells with replication-competent SIV_mac_239. ADCC was measured as the ability to direct NK cell killing of SIV_mac_239-infected CEM._NKR-_CCR5-sLTR-Luc cells. Neutralization and ADCC were calculated as the dose-dependent reduction in luciferase activity for virus or virus-infected cells incubated with antibody relative to virus or infected cells without antibody after subtracting the background luciferase signal in uninfected cells (% RLU). Values reflect the mean and standard deviation (error bars) for triplicate wells at each dilution. The dotted lines indicate half-maximal responses.

**Extended Data Fig. 5.**
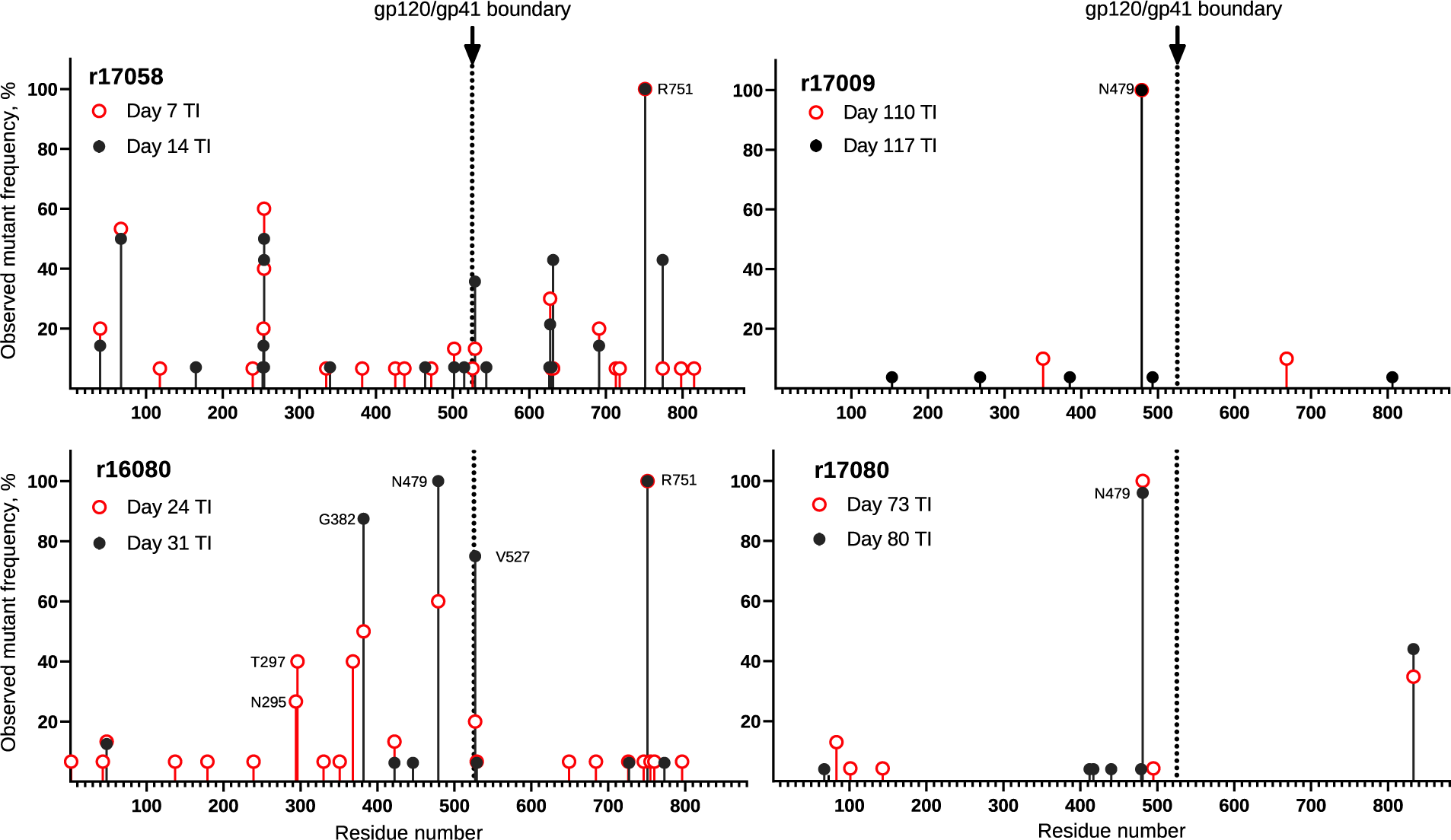
Overview of SIV Env substitutions. SIV RNA was isolated from plasma at the indicated timepoints after treatment interruption. The SIV *env* gene was amplified by RT-PCR at limiting dilution favoring amplification from a single viral genome and at least fifteen independent cDNA products were sequenced at each time point. The frequency and position of predicted amino acid changes in Env are shown.

**Extended Data Table 1.** Serum concentrations of SIV Env-specific antibodies.

| Macaque | Week 60 PI |  | Week 113 PI |  |
| --- | --- | --- | --- | --- |
| | ITS61.01<br>($\mu\text{g/ml}$) | ITS103.01<br>($\mu\text{g/ml}$) | ITS61.01<br>($\mu\text{g/ml}$) | ITS103.01<br>($\mu\text{g/ml}$) |
| r16080 | 0.13 | 28.3 | 29.2 | 53.2 |
| r17009 | 82.0 | 101.1 | 189.3 | 153.9 |
| r17040 | 8.53 | 13.4 | 51.7 | 61.4 |
| r17057 | 49.0 | 36.5 | 75.5 | 50.3 |
| r17058 | 1.80 | 0.11 | 0.02 | 0.02 |
| r17080 | 255.7 | 57.8 | 268.0 | 149.9 |
| r17114 | 6.97 | 30.9 | 82.0 | 59.9 |
| rhbn07 | 102.1 | 11.8 | 161.6 | 58.3 |

Serum concentrations of ITS61.01 and ITS103.01 at the time of ART withdrawal (week 60 PI) and one year later (week 113 PI).

**Extended Data Table 2.** Number of unique barcodes and rebound reactivation rates.

| Antibody | Macaque | Number of Unique Barcodes |  |  | Rebound Growth Rate<sup>2</sup> | Rebound Reactivation Rate<sup>2</sup> |
| --- | --- | --- | --- | --- | --- | --- |
|  |  | ART Initiation<sup>1</sup> | Primary Rebound<sup>1</sup> | Peak Rebound<sup>1</sup> |  |  |
| 17-HD9 | r16077 | 770 | 5 | 5 | 0.61 | 1.31 |
|  | r17013 | 292 | 3 | 8 | 0.91 | 1.09 |
|  | r17049 | 717 | 2 | 8 | 0.90 | 1.51 |
|  | r17090 | 453 | 3 | 8 | 1.13 | 1.36 |
|  | r17091 | 383 | 2 | 30 | 0.93 | 3.15 |
|  | r17111 | 460 | 16 | 26 | 1.15 | 5.50 |
| ITS61.01 & ITS103.01 | r16080 | 424 | 1 | 1 | 1.20 | - |
|  | r17009 | 92 | 1 | 1 | 0.68 | - |
|  | r17040 | 362 | - | - | - | - |
|  | r17057 | 266 | - | - | - | - |
|  | r17058 | 241 | 1 | 15 | 1.05 | 1.54 |
|  | r17080 | 356 | 2 | 2 | 1.40 | - |
|  | r17114 | 402 | - | - | - | - |
|  | rhbn07 | 228 | - | - | - | - |

<sup>1</sup>Number of unique SIV<sub>mac</sub>239M barcodes in plasma at the time of ART initiation (day 9 PI) and at the first and last timepoints analyzed after viral rebound in control animals (17-HD9) and animals that received vectors encoding Env-specific antibodies (ITS61.01 and ITS103.01).
<sup>2</sup>Rebound growth rates and reactivation rates (events per day) of individual control animals and 17058 were calculated using barcode proportions in plasma from the last timepoints analyzed. The rebound reactivation rates of the remaining treated animals could not be calculated by standard methods due to the limited number of rebounding clonotypes.

**Extended Data Table 3.**
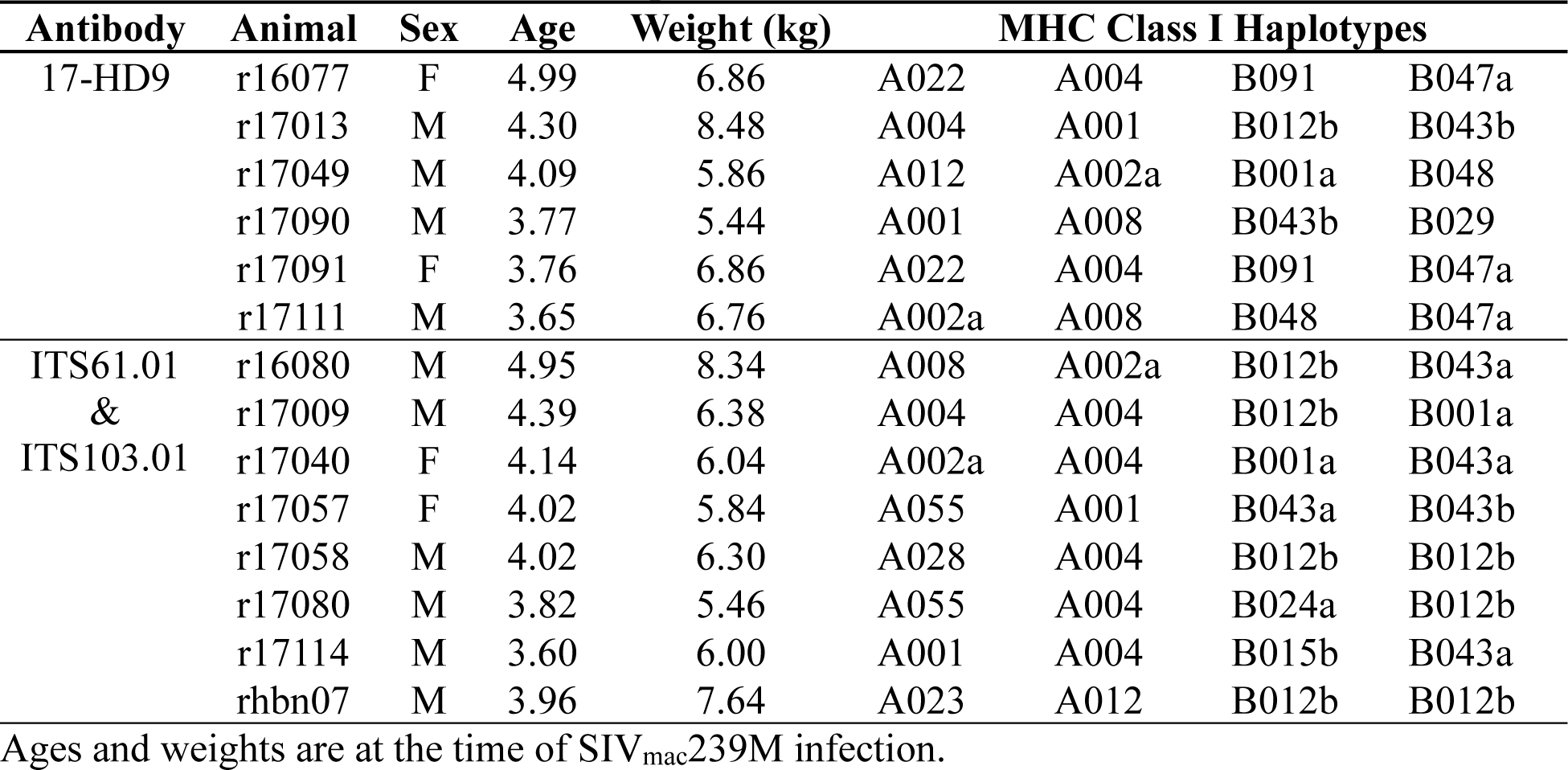
Rhesus macaques.

| **Antibody** | **Animal** | **Sex** | **Age** | **Weight (kg)** | **MHC Class I Haplotypes** |  |  |  |
| --- | --- | --- | --- | --- | --- | --- | --- | --- |
| 17-HD9 | r16077 | F | 4.99 | 6.86 | A022 | A004 | B091 | B047a |
|  | r17013 | M | 4.30 | 8.48 | A004 | A001 | B012b | B043b |
|  | r17049 | M | 4.09 | 5.86 | A012 | A002a | B001a | B048 |
|  | r17090 | M | 3.77 | 5.44 | A001 | A008 | B043b | B029 |
|  | r17091 | F | 3.76 | 6.86 | A022 | A004 | B091 | B047a |
|  | r17111 | M | 3.65 | 6.76 | A002a | A008 | B048 | B047a |
| ITS61.01<br>&<br>ITS103.01 | r16080 | M | 4.95 | 8.34 | A008 | A002a | B012b | B043a |
|  | r17009 | M | 4.39 | 6.38 | A004 | A004 | B012b | B001a |
|  | r17040 | F | 4.14 | 6.04 | A002a | A004 | B001a | B043a |
|  | r17057 | F | 4.02 | 5.84 | A055 | A001 | B043a | B043b |
|  | r17058 | M | 4.02 | 6.30 | A028 | A004 | B012b | B012b |
|  | r17080 | M | 3.82 | 5.46 | A055 | A004 | B024a | B012b |
|  | r17114 | M | 3.60 | 6.00 | A001 | A004 | B015b | B043a |
|  | rhbn07 | M | 3.96 | 7.64 | A023 | A012 | B012b | B012b |

Ages and weights are at the time of SIV<sub>mac</sub>239M infection.

**Supplemental File 1. SIV Env sequences in each animal after viral rebound (separate file).** Predicted amino acid sequences of the Env variants in plasma at time points after viral rebound are shown aligned to SIV_mac_239 Env. Positions of amino acid identity are indicated by periods, deletions are indicated by dashes and amino acid differences are identified by their single letter code.

