## Supplemental File 1 for "Adeno-associated viral delivery of Env-specific antibodies prevents SIV rebound after discontinuing antiretroviral therapy"

```
             MAC239_Env MGCLGNQLLIAILLLSVYGI YCTLYVTVFYGVPAWRNATI PLFCATKNRDTWGTTQCLPD NGDYSEVALNVTESFDAWNN TVTEQAIEDVWQLFETSIKP  100
     r16080_Day84_1_Env -------------------- -------------------- -------------------- -------------------- --------------------
     r16080_Day84_2_Env -------------------- -------------------- -------------------- -------------------- --------------------
     r16080_Day84_3_Env -------------------- -------------------- -------------------- -------------------- --------------------
     r16080_Day84_4_Env -------------------- -------------------- -------------------- -------------------- --------------------
     r16080_Day84_5_Env -------------------- -------------------- -------------------- -------------------- --------------------
     r16080_Day84_6_Env -------------------- -------------------- -------------------- -------------------- --------------------
     r16080_Day84_7_Env -------------------- -------------------- -------------------- -------------------- --------------------
     r16080_Day84_8_Env -------------------- -------------------- -------------------- -------------------- --------------------
     r16080_Day84_9_Env -------------------- -------------------- -------D------------ -------------------- --------------------
    r16080_Day84_10_Env -R------------------ -------------------- -------------------- -------------------- --------------------
    r16080_Day84_11_Env -------------------- -------------------- --L----------------- -------------------- --------------------
    r16080_Day84_12_Env -------------------- -------------------- -------------------- -------------------- --------------------
    r16080_Day84_13_Env -------------------- -------------------- -------------------- -------------------- --------------------
    r16080_Day84_14_Env -------------------- -------------------- -------------------- -------------------- --------------------
    r16080_Day84_15_Env -------------------- -------------------- -------D------------ -------------------- --------------------
 r17009_Day124TI_32_Env -------------------- -------------------- -------------------- -------------------- --------------------
 r17009_Day124TI_33_Env -------------------- -------------------- -------------------- -------------------- --------------------
 r17009_Day124TI_34_Env -------------------- -------------------- -------------------- -------------------- --------------------
 r17009_Day124TI_35_Env -------------------- -------------------- -------------------- -------------------- --------------------
 r17009_Day124TI_36_Env -------------------- -------------------- -------------------- -------------------- --------------------
 r17009_Day124TI_37_Env -------------------- -------------------- -------------------- -------------------- --------------------
 r17009_Day124TI_38_Env -------------------- -------------------- -------------------- -------------------- --------------------
 r17009_Day124TI_39_Env -------------------- -------------------- -------------------- -------------------- --------------------
 r17009_Day124TI_40_Env -------------------- -------------------- -------------------- -------------------- --------------------
 r17009_Day124TI_41_Env -------------------- -------------------- -------------------- -------------------- --------------------
 r17009_Day124TI_42_Env -------------------- -------------------- -------------------- -------------------- --------------------
 r17009_Day124TI_43_Env -------------------- -------------------- -------------------- -------------------- --------------------
 r17009_Day124TI_44_Env -------------------- -------------------- -------------------- -------------------- --------------------
 r17009_Day124TI_45_Env -------------------- -------------------- -------------------- -------------------- --------------------
 r17009_Day124TI_46_Env -------------------- -------------------- -------------------- -------------------- --------------------
 r17009_Day124TI_47_Env -------------------- -------------------- -------------------- -------------------- --------------------
 r17009_Day124TI_48_Env -------------------- -------------------- -------------------- -------------------- --------------------
 r17009_Day124TI_49_Env -------------------- -------------------- -------------------- -------------------- --------------------
 r17009_Day124TI_50_Env -------------------- -------------------- -------------------- -------------------- --------------------
 r17009_Day124TI_51_Env -------------------- -------------------- -------------------- -------------------- --------------------
 r17009_Day124TI_52_Env -------------------- -------------------- -------------------- -------------------- --------------------
 r17009_Day124TI_53_Env -------------------- -------------------- -------------------- -------------------- --------------------
 r17009_Day124TI_54_Env -------------------- -------------------- -------------------- -------------------- --------------------
 r17009_Day124TI_55_Env -------------------- -------------------- -------------------- -------------------- --------------------
 r17009_Day124TI_56_Env -------------------- -------------------- -------------------- -------------------- --------------------
 r17009_Day124TI_57_Env -------------------- -------------------- -------------------- -------------------- --------------------
r17009_Day118TI_165_Env -------------------- -------------------- -------------------- -------------------- --------------------
r17009_Day118TI_166_Env -------------------- -------------------- -------------------- -------------------- --------------------
r17009_Day118TI_167_Env -------------------- -------------------- -------------------- -------------------- --------------------
r17009_Day118TI_168_Env -------------------- -------------------- -------------------- -------------------- --------------------
r17009_Day118TI_169_Env -------------------- -------------------- -------------------- -------------------- --------------------
r17009_Day118TI_170_Env -------------------- -------------------- -------------------- -------------------- --------------------
r17009_Day118TI_171_Env -------------------- -------------------- -------------------- -------------------- --------------------
r17009_Day118TI_172_Env -------------------- -------------------- -------------------- -------------------- --------------------
r17009_Day118TI_173_Env -------------------- -------------------- -------------------- -------------------- --------------------
  r17058_Day84TI_59_Env -------------------- -------------------V -------------------- -------------------- --------------------
  r17058_Day84TI_60_Env -------------------- -------------------- -------------------- ------M------------- --------------------
  r17058_Day84TI_61_Env -------------------- -------------------- -------------------- ------M------------- --------------------
  r17058_Day84TI_62_Env -------------------- -------------------V -------------------- -------------------- --------------------
  r17058_Day84TI_63_Env -------------------- -------------------V -------------------- -------------------- --------------------
  r17058_Day84TI_64_Env -------------------- -------------------- -------------------- -------------------- --------------------
  r17058_Day84TI_65_Env -------------------- -------------------- -------------------- -------------------- --------------------
  r17058_Day84TI_66_Env -------------------- -------------------- -------------------- ------M------------- --------------------
  r17058_Day84TI_67_Env -------------------- -------------------- -------------------- -------------------- --------------------
  r17058_Day84TI_68_Env -------------------- -------------------- -------------------- ------M------------- --------------------
  r17058_Day84TI_69_Env -------------------- -------------------- -------------------- ------M------------- --------------------
  r17058_Day84TI_70_Env -------------------- -------------------- -------------------- -------------------- --------------------
  r17058_Day84TI_71_Env -------------------- -------------------- -------------------- ------M------------- --------------------
  r17058_Day84TI_72_Env -------------------- -------------------- -------------------- ------M------------- --------------------
  r17058_Day84TI_73_Env -------------------- -------------------- -------------------- ------M------------- --------------------
  r17080_Day73TI_88_Env -------------------- -------------------- -------------------- -------------------- --------------------
  r17080_Day73TI_89_Env -------------------- -------------------- -------------------- -------------------- --------------------
  r17080_Day73TI_90_Env -------------------- -------------------- -------------------- -------------------- --------------------
  r17080_Day73TI_91_Env -------------------- -------------------- -------------------- -------------------- --------------------
  r17080_Day73TI_92_Env -------------------- -------------------- -------------------- -------------------- --I-----------------
  r17080_Day73TI_93_Env -------------------- -------------------- -------------------- -------------------- --------------------
  r17080_Day73TI_94_Env -------------------- -------------------- -------------------- -------------------- --------------------
  r17080_Day73TI_95_Env -------------------- -------------------- -------------------- -------------------- --------------------
  r17080_Day73TI_96_Env -------------------- -------------------- -------------------- -------------------- --------------------
  r17080_Day73TI_97_Env -------------------- -------------------- -------------------- -------------------- --------------------
  r17080_Day73TI_98_Env -------------------- -------------------- -------------------- -------------------- --------------------
  r17080_Day73TI_99_Env -------------------- -------------------- -------------------- -------------------- --------------------
 r17080_Day73TI_100_Env -------------------- -------------------- -------------------- -------------------- --------------------
 r17080_Day73TI_101_Env -------------------- -------------------- -------------------- -------------------- --------------------
 r17080_Day73TI_102_Env -------------------- -------------------- -------------------- -------------------- --------------------
 r17080_Day73TI_103_Env -------------------- -------------------- -------------------- -------------------- --------------------
 r17080_Day73TI_104_Env -------------------- -------------------- -------------------- -------------------- --------------------
 r17080_Day73TI_105_Env -------------------- -------------------- -------------------- -------------------- --------------------
 r17080_Day73TI_106_Env -------------------- -------------------- -------------------- -------------------- --------------------
 r17080_Day73TI_107_Env -------------------- -------------------- -------------------- -------------------- --------------------
 r17080_Day73TI_108_Env -------------------- -------------------- -------------------- -------------------- --------------------
 r17080_Day73TI_109_Env -------------------- -------------------- -------------------- -------------------- --------------------
 r17080_Day73TI_110_Env -------------------- -------------------- -------------------- -------------------- --------------------
 r17080_Day77TI_111_Env -------------------- -------------------- -------------------- -------------------- --------------------
 r17080_Day77TI_112_Env -------------------- -------------------- -------------------- -------------------- --------------------
 r17080_Day77TI_113_Env -------------------- -------------------- -------------------- -------------------- --------------------
 r17080_Day77TI_114_Env -------------------- -------------------- -------------------- -------------------- --------------------
 r17080_Day77TI_115_Env -------------------- -------------------- -------------------- -------------------- -----E--------------
 r17080_Day77TI_116_Env -------------------- -------------------- -------------------- -------------------- --------------------
 r17080_Day77TI_117_Env -------------------- -------------------- -------------------- -------------------- --------------------
 r17080_Day77TI_118_Env -------------------- -------------------- -------------------- -------------------- --------------------
 r17080_Day77TI_119_Env -------------------- -------------------- -------------------- -------------------- --------------------
 r17080_Day77TI_120_Env -------------------- -------------------- -------------------- -------------------- --------------------
 r17080_Day77TI_121_Env -------------------- -------------------- -------------------- -------------------- --------------------
 r17080_Day77TI_122_Env -------------------- -------------------- ---------E---------- -------------------- --------------------
 r17080_Day77TI_123_Env -------------------- -------------------- -------------------- -------------------- --------------------
 r17080_Day77TI_124_Env -------------------- -------------------- -------------------- -------------------- --------------------
 r17080_Day77TI_125_Env -------------------- -------------------- -------------------- -------------------- --------------------
 r17080_Day77TI_126_Env -------------------- -------------------- -------------------- -------------------- --------------------
 r17080_Day77TI_127_Env -------------------- -------------------- -------------------- -------------------- --------------------
 r17080_Day77TI_128_Env -------------------- -------------------- -------------------- -------------------- --------------------
 r17080_Day77TI_129_Env -------------------- -------------------- -------------------- -------------------- --------------------
 r17080_Day77TI_130_Env -------------------- -------------------- -------------------- -------------------- --------------------
 r17080_Day77TI_131_Env -------------------- -------------------- -------------------- -------------------- --------------------
 r17080_Day77TI_132_Env -------------------- -------------------- -------------------- -------------------- --------------------
 r17080_Day77TI_133_Env -------------------- -------------------- -------------------- -------------------- --------------------
 r17080_Day77TI_134_Env -------------------- -------------------- -------------------- -------------------- --------------------
 r17080_Day77TI_135_Env -------------------- -------------------- ------R------------- -------------------- --------------------
 r17080_Day77TI_136_Env -------------------- -------------------- -------------------- -------------------- --------------------
 r17080_Day77TI_137_Env -------------------- -------------------- -------------------- -------------------- --------------------
 r17080_Day77TI_138_Env -------------------- -------------------- -------------------- -------------------- --------------------
 r17080_Day77TI_139_Env -------------------- -------------------- -------------------- -------------------- --------------------
    r16080_Day89_16_Env -------------------- -------------------- -------------------- -------------------- --------------------
    r16080_Day89_17_Env -------------------- -------------------- -------------------- -------------------- --------------------
    r16080_Day89_18_Env -------------------- -------------------- -------------------- -------------------- --------------------
    r16080_Day89_19_Env -------------------- -------------------- -------------------- -------------------- --------------------
    r16080_Day89_20_Env -------------------- -------------------- -------------------- -------------------- --------------------
    r16080_Day89_21_Env -------------------- -------------------- -------------------- -------------------- --------------------
    r16080_Day89_22_Env -------------------- -------------------- -------------------- -------------------- --------------------
    r16080_Day89_23_Env -------------------- -------------------- -------------------- -------------------- --------------------
    r16080_Day89_24_Env -------------------- -------------------- -------D------------ -------------------- --------------------
    r16080_Day89_25_Env -------------------- -------------------- -------------------- -------------------- --------------------
    r16080_Day89_26_Env -------------------- -------------------- -------------------- -------------------- --------------------
    r16080_Day89_27_Env -------------------- -------------------- -------------------- -------------------- --------------------
    r16080_Day89_28_Env -------------------- -------------------- -------D------------ -------------------- --------------------
    r16080_Day89_29_Env -------------------- -------------------- -------------------- -------------------- --------------------
    r16080_Day89_30_Env -------------------- -------------------- -------------------- -------------------- --------------------
    r16080_Day89_31_Env -------------------- -------------------- -------------------- -------------------- --------------------
r17009_Day131TI_175_Env -------------------- -------------------- -------------------- -------------------- --------------------
r17009_Day131TI_176_Env -------------------- -------------------- -------------------- -------------------- --------------------
r17009_Day131TI_177_Env -------------------- -------------------- -------------------- -------------------- --------------------
r17009_Day131TI_178_Env -------------------- -------------------- -------------------- -------------------- --------------------
r17009_Day131TI_179_Env -------------------- -------------------- -------------------- -------------------- --------------------
r17009_Day131TI_180_Env -------------------- -------------------- -------------------- -------------------- --------------------
r17009_Day131TI_181_Env -------------------- -------------------- -------------------- -------------------- --------------------
r17009_Day131TI_182_Env -------------------- -------------------- -------------------- -------------------- --------------------
r17009_Day131TI_183_Env -------------------- -------------------- -------------------- -------------------- --------------------
r17009_Day131TI_184_Env -------------------- -------------------- -------------------- -------------------- --------------------
r17009_Day131TI_185_Env -------------------- -------------------- -------------------- -------------------- --------------------
r17009_Day131TI_186_Env -------------------- -------------------- -------------------- -------------------- --------------------
r17009_Day131TI_187_Env -------------------- -------------------S -------------------- -------------------- --------------------
r17009_Day131TI_188_Env -------------------- -------------------- -------------------- -------------------- --------------------
r17009_Day131TI_189_Env -------------------- -------------------- -------------------- -------------------- --------------------
r17009_Day131TI_190_Env -------------------- -------------------- -------------------- -------------------- --------------------
r17009_Day131TI_191_Env -------------------- -------------------- -------------------- -------------------- --------------------
r17009_Day131TI_192_Env -------------------- -------------------- -------------------- -------------------- --------------------
r17009_Day131TI_193_Env -------------------- -------------------- -------------------- -------------------- --------------------
r17009_Day131TI_194_Env -------------------- -------------------- -------------------- -------------------- --------------------
r17009_Day131TI_195_Env -------------------- -------------------- -------------------- -------------------- --------------------
r17009_Day131TI_196_Env -------------------- -------------------- -------------------- -------------------- --------------------
r17009_Day131TI_197_Env -------------------- -------------------- -------------------- -------------------- --------------------
r17009_Day131TI_198_Env -------------------- -------------------- -------------------- -------------------- --------------------
r17009_Day131TI_199_Env -------------------- -------------------- -------------------- -------------------- --------------------
r17009_Day131TI_200_Env -------------------- -------------------- -------------------- -------------------- --------------------
r17009_Day131TI_201_Env -------------------- -------------------- -------------------- -------------------- --------------------
r17009_Day131TI_202_Env -------------------- -------------------- -------------------- -------------------- --------------------
 r17058__Day89TI_74_Env -------------------- -------------------- -------------------- ------M------------- --------------------
 r17058__Day89TI_75_Env -------------------- -------------------- -------------------- -------------------- --------------------
 r17058__Day89TI_76_Env -------------------- -------------------V -------------------- -------------------- --------------------
 r17058__Day89TI_77_Env -------------------- -------------------- -------------------- -------------------- --------------------
 r17058__Day89TI_78_Env -------------------- -------------------- -------------------- -------------------- --------------------
 r17058__Day89TI_79_Env -------------------- -------------------- -------------------- ------M------------- --------------------
 r17058__Day89TI_80_Env -------------------- -------------------- -------------------- ------M------------- --------------------
 r17058__Day89TI_81_Env -------------------- -------------------- -------------------- ------M------------- --------------------
 r17058__Day89TI_82_Env -------------------- -------------------- -------------------- -------------------- --------------------
 r17058__Day89TI_83_Env -------------------- -------------------- -------------------- -------------------- --------------------
 r17058__Day89TI_84_Env -------------------- -------------------- -------------------- ------M------------- --------------------
 r17058__Day89TI_85_Env -------------------- -------------------- -------------------- ------M------------- --------------------
 r17058__Day89TI_86_Env -------------------- -------------------V -------------------- -------------------- --------------------
 r17058__Day89TI_87_Env -------------------- -------------------- -------------------- ------M------------- --------------------
 r17080_Day80TI_140_Env -------------------- -------------------- -------------------- -------------------- --------------------
 r17080_Day80TI_141_Env -------------------- -------------------- -------------------- -------------------- --------------------
 r17080_Day80TI_142_Env -------------------- -------------------- -------------------- -------------------- --------------------
 r17080_Day80TI_143_Env -------------------- -------------------- -------------------- -------------------- --------------------
 r17080_Day80TI_144_Env -------------------- -------------------- -------------------- -------------------- --------------------
 r17080_Day80TI_145_Env -------------------- -------------------- -------------------- -------------------- --------------------
 r17080_Day80TI_146_Env -------------------- -------------------- -------------------- -------------------- --------------------
 r17080_Day80TI_147_Env -------------------- -------------------- -------------------- -------------------- --------------------
 r17080_Day80TI_148_Env -------------------- -------------------- -------------------- -------------------- --------------------
 r17080_Day80TI_149_Env -------------------- -------------------- -------------------- -------------------- --------------------
 r17080_Day80TI_150_Env -------------------- -------------------- -------------------- -------------------- --------------------
 r17080_Day80TI_151_Env -------------------- -------------------- -------------------- -------------------- --------------------
 r17080_Day80TI_152_Env -------------------- -------------------- -------------------- -------------------- --------------------
 r17080_Day80TI_153_Env -------------------- -------------------- -------------------- -------------------- --------------------
 r17080_Day80TI_154_Env -------------------- -------------------- -------------------- -------------------- --------------------
 r17080_Day80TI_155_Env -------------------- -------------------- -------------------- -------------------- --------------------
 r17080_Day80TI_156_Env -------------------- -------------------- -------------------- -------------------- --------------------
 r17080_Day80TI_157_Env -------------------- -------------------- -------------------- -------------------- --------------------
 r17080_Day80TI_158_Env -------------------- -------------------- -------------------- -------------------- --------------------
 r17080_Day80TI_159_Env -------------------- -------------------- -------------------- -------------------- --------------------
 r17080_Day80TI_160_Env -------------------- -------------------- -------------------- -------------------- --------------------
 r17080_Day80TI_161_Env -------------------- -------------------- -------------------- -------------------- --------------------
 r17080_Day80TI_162_Env -------------------- -------------------- -------------------- -------------------- --------------------
 r17080_Day80TI_163_Env -------------------- -------------------- -------------------- -------------------- --------------------
 r17080_Day80TI_164_Env -------------------- -------------------- -------------------- ------M------------- --------------------

             MAC239_Env CVKLSPLCITMRCNKSETDR WGLTKSITTTASTTSTTASA KVDMVNETSSCIAQDNCTGL EQEQMISCKFNMTGLKRDKK KEYNETWYSADLVCEQGNNT  200
     r16080_Day84_1_Env -------------------- -------------------- -------------------- -------------------- --------------------
     r16080_Day84_2_Env -------------------- -------------------- -------------------- ------------------E- --------------------
     r16080_Day84_3_Env -------------------- -------------------- -------------------- -------------------- --------------------
     r16080_Day84_4_Env -------------------- -------------------- -------------------- -------------------- --------------------
     r16080_Day84_5_Env -------------------- -------------------- -------------------- -------------------- --------------------
     r16080_Day84_6_Env -------------------- -------------------- -------------------- -------------------- --------------------
     r16080_Day84_7_Env -------------------- -------------------- -------------------- -------------------- --------------------
     r16080_Day84_8_Env -------------------- -------------------- -------------------- -------------------- --------------------
     r16080_Day84_9_Env -------------------- -------------------- -------------------- -------------------- --------------------
    r16080_Day84_10_Env -------------------- -------------------- -------------------- -------------------- --------------------
    r16080_Day84_11_Env -------------------- -------------------- -------------------- -------------------- --------------------
    r16080_Day84_12_Env -------------------- -------------------- -------------------- -------------------- --------------------
    r16080_Day84_13_Env -------------------- -------------------- -------------------- -------------------- --------------------
    r16080_Day84_14_Env -------------------- ----------------K--- -------------------- -------------------- --------------------
    r16080_Day84_15_Env -------------------- -------------------- -------------------- -------------------- --------------------
 r17009_Day124TI_32_Env -------------------- -------------------- -------------------- -------------------- --------------------
 r17009_Day124TI_33_Env -------------------- -------------------- -------------------- -------------------- --------------------
 r17009_Day124TI_34_Env -------------------- -------------------- -------------------- -------------------- --------------------
 r17009_Day124TI_35_Env -------------------- -------------------- -------------------- -------------------- --------------------
 r17009_Day124TI_36_Env -------------------- -------------------- -------------------- -------------------- --------------------
 r17009_Day124TI_37_Env -------------------- -------------------- -------------------- -------------------- --------------------
 r17009_Day124TI_38_Env -------------------- -------------------- -------------------- -------------------- --------------------
 r17009_Day124TI_39_Env -------------------- -------------------- -------------------- -------------------- --------------------
 r17009_Day124TI_40_Env -------------------- -------------------- -------------------- -------------------- --------------------
 r17009_Day124TI_41_Env -------------------- -------------------- -------------------- -------------------- --------------------
 r17009_Day124TI_42_Env -------------------- -------------------- -------------------- -------------------- --------------------
 r17009_Day124TI_43_Env -------------------- -------------------- -------------------- -------------------- --------------------
 r17009_Day124TI_44_Env -------------------- -------------------- -------------------- -------------------- --------------------
 r17009_Day124TI_45_Env -------------------- -------------------- -------------------- -------------------- --------------------
 r17009_Day124TI_46_Env -------------------- -------------------- ------------T------- -------------------- --------------------
 r17009_Day124TI_47_Env -------------------- -------------------- -------------------- -------------------- --------------------
 r17009_Day124TI_48_Env -------------------- -------------------- -------------------- -------------------- --------------------
 r17009_Day124TI_49_Env -------------------- -------------------- -------------------- -------------------- --------------------
 r17009_Day124TI_50_Env -------------------- -------------------- -------------------- -------------------- --------------------
 r17009_Day124TI_51_Env -------------------- -------------------- -------------------- -------------------- --------------------
 r17009_Day124TI_52_Env -------------------- -------------------- -------------------- -------------------- --------------------
 r17009_Day124TI_53_Env -------------------- -------------------- -------------------- -------------------- --------------------
 r17009_Day124TI_54_Env -------------------- -------------------- -------------------- -------------------- --------------------
 r17009_Day124TI_55_Env -------------------- -------------------- -------------------- -------------------- --------------------
 r17009_Day124TI_56_Env -------------------- -------------------- -------------------- -------------------- --------------------
 r17009_Day124TI_57_Env -------------------- -------------------- -------------------- -------------------- --------------------
r17009_Day118TI_165_Env -------------------- -------------------- -------------------- -------------------- --------------------
r17009_Day118TI_166_Env -------------------- -------------------- -------------------- -------------------- --------------------
r17009_Day118TI_167_Env -------------------- -------------------- -------------------- -------------------- --------------------
r17009_Day118TI_168_Env -------------------- -------------------- -------------------- -------------------- --------------------
r17009_Day118TI_169_Env -------------------- -------------------- -------------------- -------------------- --------------------
r17009_Day118TI_170_Env -------------------- -------------------- -------------------- -------------------- --------------------
r17009_Day118TI_171_Env -------------------- -------------------- -------------------- -------------------- --------------------
r17009_Day118TI_172_Env -------------------- -------------------- -------------------- -------------------- --------------------
r17009_Day118TI_173_Env -------------------- -------------------- -------------------- -------------------- --------------------
  r17058_Day84TI_59_Env -------------------- -------------------- -------------------- -------------------- --------------------
  r17058_Day84TI_60_Env -------------------- -------------------- -------------------- -------------------- --------------------
  r17058_Day84TI_61_Env -----------------A-- -------------------- -------------------- -------------------- --------------------
  r17058_Day84TI_62_Env -------------------- -------------------- -------------------- -------------------- --------------------
  r17058_Day84TI_63_Env -------------------- -------------------- -------------------- -------------------- --------------------
  r17058_Day84TI_64_Env -------------------- -------------------- -------------------- -------------------- --------------------
  r17058_Day84TI_65_Env -------------------- -------------------- -------------------- -------------------- --------------------
  r17058_Day84TI_66_Env -------------------- -------------------- -------------------- -------------------- --------------------
  r17058_Day84TI_67_Env -------------------- -------------------- -------------------- -------------------- --------------------
  r17058_Day84TI_68_Env -------------------- -------------------- -------------------- -------------------- --------------------
  r17058_Day84TI_69_Env -------------------- -------------------- -------------------- -------------------- --------------------
  r17058_Day84TI_70_Env -------------------- -------------------- -------------------- -------------------- --------------------
  r17058_Day84TI_71_Env -------------------- -------------------- -------------------- -------------------- --------------------
  r17058_Day84TI_72_Env -------------------- -------------------- -------------------- -------------------- --------------------
  r17058_Day84TI_73_Env -------------------- -------------------- -------------------- -------------------- --------------------
  r17080_Day73TI_88_Env -------------------- -------------------- -------------------- -------------------- --------------------
  r17080_Day73TI_89_Env -------------------- -------------------- -------------------- -------------------- --------------------
  r17080_Day73TI_90_Env -------------------- -------------------- -------------------- -------------------- --------------------
  r17080_Day73TI_91_Env -------------------- -------------------- -------------------- -------------------- --------------------
  r17080_Day73TI_92_Env -------------------- -------------------- -------------------- -------------------- --------------------
  r17080_Day73TI_93_Env -------------------- -------------------- -------------------- -------------------- --------------------
  r17080_Day73TI_94_Env -------------------- -------------------- -------------------- -------------------- --------------------
  r17080_Day73TI_95_Env -------------------- -------------------- -------------------- -------------------- --------------------
  r17080_Day73TI_96_Env -------------------- -------------------- -------------------- -------------------- --------------------
  r17080_Day73TI_97_Env -------------------- -------------------- -------------------- -------------------- --------------------
  r17080_Day73TI_98_Env -------------------- -------------------- -------------------- -------------------- --------------------
  r17080_Day73TI_99_Env -------------------- -------------------- -------------------- -------------------- --------------------
 r17080_Day73TI_100_Env -------------------- -------------------- -------------------- -------------------- --------------------
 r17080_Day73TI_101_Env -------------------- -------------------- --G----------------- -------------------- --------------------
 r17080_Day73TI_102_Env Y------------------- -------------------- -------------------- -------------------- --------------------
 r17080_Day73TI_103_Env -------------------- -------------------- -------------------- -------------------- --------------------
 r17080_Day73TI_104_Env -------------------- -------------------- -------------------- -------------------- --------------------
 r17080_Day73TI_105_Env -------------------- -------------------- -------------------- -------------------- --------------------
 r17080_Day73TI_106_Env -------------------- -------------------- -------------------- -------------------- --------------------
 r17080_Day73TI_107_Env -------------------- -------------------- -------------------- -------------------- --------------------
 r17080_Day73TI_108_Env -------------------- -------------------- -------------------- -------------------- --------------------
 r17080_Day73TI_109_Env -------------------- -------------------- -------------------- -------------------- --------------------
 r17080_Day73TI_110_Env -------------------- -------------------- -------------------- -------------------- --------------------
 r17080_Day77TI_111_Env -------------------- -------------------- -------------------- -------------------- --------------------
 r17080_Day77TI_112_Env -------------------- -------------------- -------------------- -------------------- --------------------
 r17080_Day77TI_113_Env -------------------- -------------------- -------------------- -------------------- --------------------
 r17080_Day77TI_114_Env -------------------- -------------------- -------------------- -------------------- --------------------
 r17080_Day77TI_115_Env -------------------- -------------------- -------------------- -------------------- --------------------
 r17080_Day77TI_116_Env -------------------- -------------------- -------------------- -------------------- --------------------
 r17080_Day77TI_117_Env -------------------- -------------------- -------------------- -------------------- --------------------
 r17080_Day77TI_118_Env -------------------- -------------------- -------------------- -------------------- --------------------
 r17080_Day77TI_119_Env -------------------- -------------------- -------------------- -------------------- --------------------
 r17080_Day77TI_120_Env -------------------- -------------------- -------------------- -------------------- --------------------
 r17080_Day77TI_121_Env -------------------- -------------------- -------------------- -------------------- --------------------
 r17080_Day77TI_122_Env -------------------- -------------------- -------------------- -------------------- --------------------
 r17080_Day77TI_123_Env -------------------- -------------------- -------------------- -------------------- --------------------
 r17080_Day77TI_124_Env -------------------- -------------------- -------------------- -------------------- --------------------
 r17080_Day77TI_125_Env -------------------- -------------------- -------------------- -------------------- --------------------
 r17080_Day77TI_126_Env -------------------- -------------------- -------------------- -------------------- --------------------
 r17080_Day77TI_127_Env -------------------- -------------------- -------------------- -------------------- --------------------
 r17080_Day77TI_128_Env -------------------- -------------------- -------------------- -------------------- --------------------
 r17080_Day77TI_129_Env -------------------- -------------------- -------------------- -------------------- --------------------
 r17080_Day77TI_130_Env -------------------- -------------------- -------------------- -------------------- --------------------
 r17080_Day77TI_131_Env -------------------- -------------------- -------------------- -------------------- --------------------
 r17080_Day77TI_132_Env -------------------- -------------------- -------------------- -------------------- --------------------
 r17080_Day77TI_133_Env -------------------- -------------------- -------------------- -------------------- --------------------
 r17080_Day77TI_134_Env -------------------- -------------------- -------------------- -------------------- --------------------
 r17080_Day77TI_135_Env -------------------- -------------------- -------------------- -------------------- --------------------
 r17080_Day77TI_136_Env -------------------- -------------------- -------------------- -------------------- --------------------
 r17080_Day77TI_137_Env -------------------- -------------------- -------------------- -------------------- --------------------
 r17080_Day77TI_138_Env -------------------- -------------------- -------------------- -------------------- --------------------
 r17080_Day77TI_139_Env -------------------- -------------------- -------------------- -------------------- --------------------
    r16080_Day89_16_Env -------------------- -------------------- -------------------- -------------------- --------------------
    r16080_Day89_17_Env -------------------- -------------------- -------------------- -------------------- --------------------
    r16080_Day89_18_Env -------------------- -------------------- -------------------- -------------------- --------------------
    r16080_Day89_19_Env -------------------- -------------------- -------------------- -------------------- --------------------
    r16080_Day89_20_Env -------------------- -------------------- -------------------- -------------------- --------------------
    r16080_Day89_21_Env -------------------- -------------------- -------------------- -------------------- --------------------
    r16080_Day89_22_Env -------------------- -------------------- -------------------- -------------------- --------------------
    r16080_Day89_23_Env -------------------- -------------------- -------------------- -------------------- --------------------
    r16080_Day89_24_Env -------------------- -------------------- -------------------- -------------------- --------------------
    r16080_Day89_25_Env -------------------- -------------------- -------------------- -------------------- --------------------
    r16080_Day89_26_Env -------------------- -------------------- -------------------- -------------------- --------------------
    r16080_Day89_27_Env -------------------- -------------------- -------------------- -------------------- --------------------
    r16080_Day89_28_Env -------------------- -------------------- -------------------- -------------------- --------------------
    r16080_Day89_29_Env -------------------- -------------------- -------------------- -------------------- --------------------
    r16080_Day89_30_Env -------------------- -------------------- -------------------- -------------------- --------------------
    r16080_Day89_31_Env -------------------- -------------------- -------------------- -------------------- --------------------
r17009_Day131TI_175_Env -------------------- -------------------- -------------------- -------------------- --------------------
r17009_Day131TI_176_Env -------------------- -------------------- -------------------- -------------------- --------------------
r17009_Day131TI_177_Env -------------------- -------------------- -------------------- -------------------- --------------------
r17009_Day131TI_178_Env -------------------- -------------------- -------------------- -------------------- --------------------
r17009_Day131TI_179_Env -------------------- -------------------- -------------------- -------------------- --------------------
r17009_Day131TI_180_Env -------------------- -------------------- -------------------- -------------------- --------------------
r17009_Day131TI_181_Env -------------------- -------------------- -------------------- -------------------- --------------------
r17009_Day131TI_182_Env -------------------- -------------------- -------------------- -------------------- --------------------
r17009_Day131TI_183_Env -------------------- -------------------- -------------------- -------------------- --------------------
r17009_Day131TI_184_Env -------------------- -------------------- -------------------- -------------------- --------------------
r17009_Day131TI_185_Env -------------------- -------------------- -------------------- -------------------- --------------------
r17009_Day131TI_186_Env -------------------- -------------------- -------------------- -------------------- --------------------
r17009_Day131TI_187_Env -------------------- -------------------- -------------------- -------------------- --------------------
r17009_Day131TI_188_Env -------------------- -------------------- -------------------- -------------------- --------------------
r17009_Day131TI_189_Env -------------------- -------------------- -------------------- -------------------- --------------------
r17009_Day131TI_190_Env -------------------- -------------------- -------------------- -------------------- --------------------
r17009_Day131TI_191_Env -------------------- -------------------- -------------------- -------------------- --------------------
r17009_Day131TI_192_Env -------------------- -------------------- -------------------- -------------------- --------------------
r17009_Day131TI_193_Env -------------------- -------------------- -------------------- -------------------- --------------------
r17009_Day131TI_194_Env -------------------- -------------------- -------------------- -------------------- --------------------
r17009_Day131TI_195_Env -------------------- -------------------- -------------------- -------------------- --------------------
r17009_Day131TI_196_Env -------------------- -------------------- -------------------- -------------------- --------------------
r17009_Day131TI_197_Env -------------------- -------------------- -------------------- -------------------- --------------------
r17009_Day131TI_198_Env -------------------- -------------------- -------------------- -------------------- --------------------
r17009_Day131TI_199_Env -------------------- -------------------- -------------------- -------------------- --------------------
r17009_Day131TI_200_Env -------------------- -------------------- -------------------- -------------------- --------------------
r17009_Day131TI_201_Env -------------------- -------------------- -------------------- -------------------- --------------------
r17009_Day131TI_202_Env -------------------- -------------------- -------------------- -------------------- --------------------
 r17058__Day89TI_74_Env -------------------- -------------------- -------------------- -------------------- --------------------
 r17058__Day89TI_75_Env -------------------- -------------------- -------------------- -------------------- --------------------
 r17058__Day89TI_76_Env -------------------- -------------------- -------------------- -------------------- --------------------
 r17058__Day89TI_77_Env -------------------- -------------------- -------------------- -------------------- --------------------
 r17058__Day89TI_78_Env -------------------- -------------------- -------------------- -------------------- --------------------
 r17058__Day89TI_79_Env -------------------- -------------------- -------------------- -------------------- --------------------
 r17058__Day89TI_80_Env -------------------- -------------------- -------------------- ----T--------------- --------------------
 r17058__Day89TI_81_Env -------------------- -------------------- -------------------- -------------------- --------------------
 r17058__Day89TI_82_Env -------------------- -------------------- -------------------- -------------------- --------------------
 r17058__Day89TI_83_Env -------------------- -------------------- -------------------- -------------------- --------------------
 r17058__Day89TI_84_Env -------------------- -------------------- -------------------- -------------------- --------------------
 r17058__Day89TI_85_Env -------------------- -------------------- -------------------- -------------------- --------------------
 r17058__Day89TI_86_Env -------------------- -------------------- -------------------- -------------------- --------------------
 r17058__Day89TI_87_Env -------------------- -------------------- -------------------- -------------------- --------------------
 r17080_Day80TI_140_Env -------------------- -------------------- -------------------- -------------------- --------------------
 r17080_Day80TI_141_Env -------------------- -------------------- -------------------- -------------------- --------------------
 r17080_Day80TI_142_Env -------------------- -------------------- -------------------- -------------------- --------------------
 r17080_Day80TI_143_Env -------------------- -------------------- -------------------- -------------------- --------------------
 r17080_Day80TI_144_Env -------------------- -------------------- -------------------- -------------------- --------------------
 r17080_Day80TI_145_Env -------------------- -------------------- -------------------- -------------------- --------------------
 r17080_Day80TI_146_Env -------------------- -------------------- -------------------- -------------------- --------------------
 r17080_Day80TI_147_Env -------------------- -------------------- -------------------- -------------------- --------------------
 r17080_Day80TI_148_Env -------------------- -------------------- -------------------- -------------------- --------------------
 r17080_Day80TI_149_Env -------------------- -------------------- -------------------- -------------------- --------------------
 r17080_Day80TI_150_Env -------------------- -------------------- -------------------- -------------------- --------------------
 r17080_Day80TI_151_Env -------------------- -------------------- -------------------- -------------------- --------------------
 r17080_Day80TI_152_Env -------------------- -------------------- -------------------- -------------------- --------------------
 r17080_Day80TI_153_Env -------------------- -------------------- -------------------- -------------------- --------------------
 r17080_Day80TI_154_Env -------------------- -------------------- -------------------- -------------------- --------------------
 r17080_Day80TI_155_Env -------------------- -------------------- -------------------- -------------------- --------------------
 r17080_Day80TI_156_Env -------------------- -------------------- -------------------- -------------------- --------------------
 r17080_Day80TI_157_Env -------------------- -------------------- -------------------- -------------------- --------------------
 r17080_Day80TI_158_Env -------------------- -------------------- -------------------- -------------------- --------------------
 r17080_Day80TI_159_Env -------------------- -------------------- -------------------- -------------------- --------------------
 r17080_Day80TI_160_Env -------------------- -------------------- -------------------- -------------------- --------------------
 r17080_Day80TI_161_Env -------------------- -------------------- -------------------- -------------------- --------------------
 r17080_Day80TI_162_Env -------------------- -------------------- -------------------- -------------------- --------------------
 r17080_Day80TI_163_Env -------------------- -------------------- -------------------- -------------------- --------------------
 r17080_Day80TI_164_Env -------------------- -------------------- -------------------- -------------------- --------------------

             MAC239_Env GNESRCYMNHCNTSVIQESC DKHYWDAIRFRYCAPPGYAL LRCNDTNYSGFMPKCSKVVV SSCTRMMETQTSTWFGFNGT RAENRTYIYWHGRDNRTIIS  300
     r16080_Day84_1_Env -------------------- -------------------- -------------------- -------------------- --------------------
     r16080_Day84_2_Env -------------------- -------------------- -------------------- -------------------- --------------------
     r16080_Day84_3_Env -------------------- -------------------- -------------------- -------------------- --------------------
     r16080_Day84_4_Env -------------------- ------------------V- -------------------- -------------------- --------------------
     r16080_Day84_5_Env -------------------- -------------------- -------------------- -------------------- --------------------
     r16080_Day84_6_Env -------------------- -------------------- -------------------- -------------------- --------------D-V---
     r16080_Day84_7_Env -------------------- -------------------- -------------------- -------------------- --------------D-V---
     r16080_Day84_8_Env -------------------- -------------------- -------------------- -------------------- --------------------
     r16080_Day84_9_Env -------------------- -------------------- -------------------- -------------------- ----------------V---
    r16080_Day84_10_Env -------------------- -------------------- -------------------- -------------------- --------------------
    r16080_Day84_11_Env -------------------- -------------------- -------------------- -------------------- --------------------
    r16080_Day84_12_Env -------------------- -------------------- -------------------- -------------------- --------------D-V---
    r16080_Day84_13_Env -------------------- -------------------- -------------------- -------------------- --------------------
    r16080_Day84_14_Env -------------------- -------------------- -------------------- -------------------- --------------D-V---
    r16080_Day84_15_Env -------------------- -------------------- -------------------- -------------------- ----------------V---
 r17009_Day124TI_32_Env -------------------- -------------------- -------------------- -------------------- --------------------
 r17009_Day124TI_33_Env -------------------- -------------------- -------------------- -------------------- --------------------
 r17009_Day124TI_34_Env -------------------- -------------------- -------------------- -------------------- --------------------
 r17009_Day124TI_35_Env -------------------- -------------------- -------------------- -------------------- --------------------
 r17009_Day124TI_36_Env -------------------- -------------------- -------------------- -------------------- --------------------
 r17009_Day124TI_37_Env -------------------- -------------------- -------------------- -------------------- --------------------
 r17009_Day124TI_38_Env -------------------- -------------------- -------------------- -------------------- --------------------
 r17009_Day124TI_39_Env -------------------- -------------------- -------------------- -------------------- --------------------
 r17009_Day124TI_40_Env -------------------- -------------------- -------------------- -------------------- --------------------
 r17009_Day124TI_41_Env -------------------- -------------------- -------------------- -------------------- --------------------
 r17009_Day124TI_42_Env -------------------- -------------------- -------------------- -------------------- --------------------
 r17009_Day124TI_43_Env -------------------- -------------------- -------------------- -------------------- --------------------
 r17009_Day124TI_44_Env -------------------- -------------------- -------------------- -------------------- --------------------
 r17009_Day124TI_45_Env -------------------- -------------------- -------------------- -------------------- --------------------
 r17009_Day124TI_46_Env -------------------- -------------------- -------------------- -------------------- --------------------
 r17009_Day124TI_47_Env -------------------- -------------------- -------------------- -------------------- --------------------
 r17009_Day124TI_48_Env -------------------- -------------------- -------------------- -------------------- --------------------
 r17009_Day124TI_49_Env -------------------- -------------------- -------------------- -------------------- --------------------
 r17009_Day124TI_50_Env -------------------- -------------------- -------------------- -------------------- --------------------
 r17009_Day124TI_51_Env -------------------- -------------------- -------------------- -------------------- --------------------
 r17009_Day124TI_52_Env -------------------- -------------------- -------------------- -------------------- --------------------
 r17009_Day124TI_53_Env -------------------- -------------------- -------------------- -------------------- --------------------
 r17009_Day124TI_54_Env -------------------- -------------------- -------------------- -------------------- --------------------
 r17009_Day124TI_55_Env -------------------- -------------------- -------------------- -------------------- --------------------
 r17009_Day124TI_56_Env -------------------- -------------------- -------------------- -------------------- --------------------
 r17009_Day124TI_57_Env -------------------- -------------------- -------------------- -------K------------ --------------------
r17009_Day118TI_165_Env -------------------- -------------------- -------------------- -------------------- --------------------
r17009_Day118TI_166_Env -------------------- -------------------- -------------------- -------------------- --------------------
r17009_Day118TI_167_Env -------------------- -------------------- -------------------- -------------------- --------------------
r17009_Day118TI_168_Env -------------------- -------------------- -------------------- -------------------- --------------------
r17009_Day118TI_169_Env -------------------- -------------------- -------------------- -------------------- --------------------
r17009_Day118TI_170_Env -------------------- -------------------- -------------------- -------------------- --------------------
r17009_Day118TI_171_Env -------------------- -------------------- -------------------- -------------------- --------------------
r17009_Day118TI_172_Env -------------------- -------------------- -------------------- -------------------- --------------------
r17009_Day118TI_173_Env -------------------- -------------------- -------------------- -------------------- --------------------
  r17058_Day84TI_59_Env -------------------- -------------------- -------------E------ -------------------- --------------------
  r17058_Day84TI_60_Env -------------------- -------------------- ------------SN------ -------------------- --------------------
  r17058_Day84TI_61_Env -------------------- -------------------- -------------N------ -------------------- --------------------
  r17058_Day84TI_62_Env -------------------- -------------------- -------------E------ -------------------- --------------------
  r17058_Day84TI_63_Env -------------------- -------------------- -------------E------ -------------------- --------------------
  r17058_Day84TI_64_Env -------------------- -------------------- -------------E------ -------------------- --------------------
  r17058_Day84TI_65_Env -------------------- ------------------V- -------------E------ -------------------- --------------------
  r17058_Day84TI_66_Env -------------------- -------------------- ------------SN------ -------------------- --------------------
  r17058_Day84TI_67_Env -------------------- -------------------- -------------E------ -------------------- --------------------
  r17058_Day84TI_68_Env -------------------- -------------------- -------------E------ -------------------- --------------------
  r17058_Day84TI_69_Env -------------------- -------------------- -------------N------ -------------------- --------------------
  r17058_Day84TI_70_Env -------------------- -------------------- -------------E------ -------------------- --------------------
  r17058_Day84TI_71_Env -------------------- -------------------- -------------N------ -------------------- --------------------
  r17058_Day84TI_72_Env -------------------- -------------------- -------------E------ -------------------- --------------------
  r17058_Day84TI_73_Env -------------------- -------------------- ------------SN------ -------------------- --------------------
  r17080_Day73TI_88_Env -------------------- -------------------- -------------------- -------------------- --------------------
  r17080_Day73TI_89_Env -------------------- -------------------- -------------------- -------------------- --------------------
  r17080_Day73TI_90_Env -------------------- -------------------- -------------------- -------------------- --------------------
  r17080_Day73TI_91_Env -------------------- -------------------- -------------------- -------------------- --------------------
  r17080_Day73TI_92_Env -------------------- -------------------- -------------------- -------------------- --------------------
  r17080_Day73TI_93_Env -------------------- -------------------- -------------------- -------------------- --------------------
  r17080_Day73TI_94_Env -------------------- -------------------- -------------------- -------------------- --------------------
  r17080_Day73TI_95_Env -------------------- -------------------- -------------------- -------------------- --------------------
  r17080_Day73TI_96_Env -------------------- -------------------- -------------------- -------------------- --------------------
  r17080_Day73TI_97_Env -------------------- -------------------- -------------------- -------------------- --------------------
  r17080_Day73TI_98_Env -------------------- -------------------- -------------------- -------------------- --------------------
  r17080_Day73TI_99_Env -------------------- -------------------- -------------------- -------------------- --------------------
 r17080_Day73TI_100_Env -------------------- -------------------- -------------------- -------------------- --------------------
 r17080_Day73TI_101_Env -------------------- -------------------- -------------------- -------------------- --------------------
 r17080_Day73TI_102_Env -------------------- -------------------- -------------------- -------------------- --------------------
 r17080_Day73TI_103_Env -------------------- -------------------- -------------------- -------------------- --------------------
 r17080_Day73TI_104_Env -------------------- -------------------- -------------------- -------------------- --------------------
 r17080_Day73TI_105_Env -------------------- -------------------- -------------------- -------------------- --------------------
 r17080_Day73TI_106_Env -------------------- -------------------- -------------------- -------------------- --------------------
 r17080_Day73TI_107_Env -------------------- -------------------- -------------------- -------------------- --------------------
 r17080_Day73TI_108_Env -------------------- -------------------- -------------------- -------------------- --------------------
 r17080_Day73TI_109_Env -------------------- -------------------- -------------------- -------------------- --------------------
 r17080_Day73TI_110_Env -------------------- -------------------- -------------------- -------------------- --------------------
 r17080_Day77TI_111_Env -------------------- -------------------- -------------------- -------------------- --------------------
 r17080_Day77TI_112_Env -------------------- -------------------- -------------------- -------------------- --------------------
 r17080_Day77TI_113_Env -------------------- -------------------- -------------------- -------------------- --------------------
 r17080_Day77TI_114_Env -------------------- -------------------- -------------------- -------------------- --------------------
 r17080_Day77TI_115_Env -------------------- -------------------- -------------------- -------------------- --------------------
 r17080_Day77TI_116_Env -------------------- -------------------- -------------------- -------------------- --------------------
 r17080_Day77TI_117_Env -------------------- -------------------- -------------------- -------------------- --------------------
 r17080_Day77TI_118_Env -------------------- -------------------- -------------------- -------------------- --------------------
 r17080_Day77TI_119_Env -------------------- -------------------- -------------------- -------------------- --------------------
 r17080_Day77TI_120_Env -------------------- -------------------- -------------------- -------------------- --------------------
 r17080_Day77TI_121_Env -------------------- -------------------- -------------------- -------------------- --------------------
 r17080_Day77TI_122_Env -------------------- -------------------- -------------------- -------------------- --------------------
 r17080_Day77TI_123_Env -------------------- -------------------- -------------------- -------------------- --------------------
 r17080_Day77TI_124_Env -------------------- -------------------- -------------------- -------------------- --------------------
 r17080_Day77TI_125_Env -------------------- -------------------- -------------------- -------------------- --------------------
 r17080_Day77TI_126_Env -------------------- -------------------- -------------------- -------------------- --------------------
 r17080_Day77TI_127_Env -------------------- -------------------- -------------------- -------------------- --------------------
 r17080_Day77TI_128_Env -------------------- -------------------- -------------------- -------------------- --------------------
 r17080_Day77TI_129_Env -------------------- -------------------- -------------------- -------------------- --------------------
 r17080_Day77TI_130_Env -------------------- -------------------- -------------------- -------------------- --------------------
 r17080_Day77TI_131_Env -------------------- -------------------- -------------------- -------------------- --------------------
 r17080_Day77TI_132_Env -------------------- -------------------- -------------------- -------------------- --------------------
 r17080_Day77TI_133_Env -------------------- -------------------- -------------------- -------------------- --------------------
 r17080_Day77TI_134_Env -------------------- -------------------- -------------------- -------------------- --------------------
 r17080_Day77TI_135_Env -------------------- -------------------- -------------------- -------------------- --------------------
 r17080_Day77TI_136_Env -------------------- -------------------- -------------------- -------------------- --------------------
 r17080_Day77TI_137_Env -------------------- -------------------- -------------------- -------------------- --------------------
 r17080_Day77TI_138_Env -------------------- -------------------- -------------------- -------------------- --------------------
 r17080_Day77TI_139_Env -------------------- -------------------- -------------------- -------------------- --------------------
    r16080_Day89_16_Env -------------------- -------------------- -------------------- -------------------- --------------------
    r16080_Day89_17_Env -------------------- -------------------- -------------------- -------------------- --------------------
    r16080_Day89_18_Env -------------------- -------------------- -------------------- -------------------- --------------------
    r16080_Day89_19_Env -------------------- -------------------- -------------------- -------------------- --------------------
    r16080_Day89_20_Env -------------------- -------------------- -------------------- -------------------- --------------------
    r16080_Day89_21_Env -------------------- -------------------- -------------------- -------------------- --------------------
    r16080_Day89_22_Env -------------------- -------------------- -------------------- -------------------- --------------------
    r16080_Day89_23_Env -------------------- -------------------- -------------------- -------------------- --------------------
    r16080_Day89_24_Env -------------------- -------------------- -------------------- -------------------- --------------------
    r16080_Day89_25_Env -------------------- -------------------- -------------------- -------------------- --------------------
    r16080_Day89_26_Env -------------------- -------------------- -------------------- -------------------- --------------------
    r16080_Day89_27_Env -------------------- -------------------- -------------------- -------------------- --------------------
    r16080_Day89_28_Env -------------------- -------------------- -------------------- -------------------- --------------------
    r16080_Day89_29_Env -------------------- -------------------- -------------------- -------------------- --------------------
    r16080_Day89_30_Env -------------------- -------------------- -------------------- -------------------- --------------------
    r16080_Day89_31_Env -------------------- -------------------- -------------------- -------------------- --------------------
r17009_Day131TI_175_Env -------------------- -------------------- -------------------- -------------------- --------------------
r17009_Day131TI_176_Env -------------------- -------------------- -------------------- -------------------- --------------------
r17009_Day131TI_177_Env -------------------- -------------------- -------------------- -------------------- --------------------
r17009_Day131TI_178_Env -------------------- -------------------- -------------------- -------------------- --------------------
r17009_Day131TI_179_Env -------------------- -------------------- -------------------- -------------------- --------------------
r17009_Day131TI_180_Env -------------------- -------------------- -------------------- -------------------- --------------------
r17009_Day131TI_181_Env -------------------- -------------------- -------------------- -------------------- --------------------
r17009_Day131TI_182_Env -------------------- -------------------- -------------------- -------------------- --------------------
r17009_Day131TI_183_Env -------------------- -------------------- -------------------- -------------------- --------------------
r17009_Day131TI_184_Env -------------------- -------------------- -------------------- -------------------- --------------------
r17009_Day131TI_185_Env -------------------- -------------------- -------------------- -------------------- --------------------
r17009_Day131TI_186_Env -------------------- -------------------- -------------------- -------------------- --------------------
r17009_Day131TI_187_Env -------------------- -------------------- -------------------- -------------------- --------------------
r17009_Day131TI_188_Env -------------------- -------------------- -------------------- -------------------- --------------------
r17009_Day131TI_189_Env -------------------- -------------------- -------------------- -------------------- --------------------
r17009_Day131TI_190_Env -------------------- -------------------- -------------------- -------------------- --------------------
r17009_Day131TI_191_Env -------------------- -------------------- -------------------- -------------------- --------------------
r17009_Day131TI_192_Env -------------------- -------------------- -------------------- -------------------- --------------------
r17009_Day131TI_193_Env -------------------- -------------------- -------------------- -------------------- --------------------
r17009_Day131TI_194_Env -------------------- -------------------- -------------------- -------------------- --------------------
r17009_Day131TI_195_Env -------------------- -------------------- -------------------- -------------------- --------------------
r17009_Day131TI_196_Env -------------------- -------------------- -------------------- -------------------- --------------------
r17009_Day131TI_197_Env -------------------- -------------------- -------------------- -------------------- --------------------
r17009_Day131TI_198_Env -------------------- -------------------- -------------------- -------------------- --------------------
r17009_Day131TI_199_Env -------------------- -------------------- -------------------- -----T-------------- --------------------
r17009_Day131TI_200_Env -------------------- -------------------- -------------------- -------------------- --------------------
r17009_Day131TI_201_Env -------------------- -------------------- -------------------- -------------------- --------------------
r17009_Day131TI_202_Env -------------------- -------------------- -------------------- -------------------- --------------------
 r17058__Day89TI_74_Env -------------------- -------------------- -------------E------ -------------------- --------------------
 r17058__Day89TI_75_Env -------------------- -------------------- -------------E------ -------------------- --------------------
 r17058__Day89TI_76_Env -------------------- -------------------- -------------E------ -------------------- --------------------
 r17058__Day89TI_77_Env -------------------- -------------------- -------------E------ -------------------- --------------------
 r17058__Day89TI_78_Env -------------------- -------------------- -------------N------ -------------------- --------------------
 r17058__Day89TI_79_Env -------------------- -------------------- -----------ISD------ -------------------- --------------------
 r17058__Day89TI_80_Env -------------------- -------------------- -------------N------ -------------------- --------------------
 r17058__Day89TI_81_Env -------------------- -------------------- ------------SN------ -------------------- --------------------
 r17058__Day89TI_82_Env -------------------- -------------------- -------------E------ -------------------- --------------------
 r17058__Day89TI_83_Env -------------------- -------------------- -------------N------ -------------------- --------------------
 r17058__Day89TI_84_Env -------------------- -------------------- -------------E------ -------------------- --------------------
 r17058__Day89TI_85_Env -------------------- -------------------- -------------N------ -------------------- --------------------
 r17058__Day89TI_86_Env -------------------- -------------------- -------------E------ -------------------- --------------------
 r17058__Day89TI_87_Env -------------------- -------------------- -------------N------ -------------------- --------------------
 r17080_Day80TI_140_Env -------------------- -------------------- -------------------- -------------------- --------------------
 r17080_Day80TI_141_Env -------------------- -------------------- -------------------- -------------------- --------------------
 r17080_Day80TI_142_Env -------------------- -------------------- -------------------- -------------------- --------------------
 r17080_Day80TI_143_Env -------------------- -------------------- -------------------- -------------------- --------------------
 r17080_Day80TI_144_Env -------------------- -------------------- -------------------- -------------------- --------------------
 r17080_Day80TI_145_Env -------------------- -------------------- -------------------- -------------------- --------------------
 r17080_Day80TI_146_Env -------------------- -------------------- -------------------- -------------------- --------------------
 r17080_Day80TI_147_Env -------------------- -------------------- -------------------- -------------------- --------------------
 r17080_Day80TI_148_Env -------------------- -------------------- -------------------- -------------------- --------------------
 r17080_Day80TI_149_Env -------------------- -------------------- -------------------- -------------------- --------------------
 r17080_Day80TI_150_Env -------------------- -------------------- -------------------- -------------------- --------------------
 r17080_Day80TI_151_Env -------------------- -------------------- -------------------- -------------------- --------------------
 r17080_Day80TI_152_Env -------------------- -------------------- -------------------- -------------------- --------------------
 r17080_Day80TI_153_Env -------------------- -------------------- -------------------- -------------------- --------------------
 r17080_Day80TI_154_Env -------------------- -------------------- -------------------- -------------------- --------------------
 r17080_Day80TI_155_Env -------------------- -------------------- -------------------- -------------------- --------------------
 r17080_Day80TI_156_Env -------------------- -------------------- -------------------- -------------------- --------------------
 r17080_Day80TI_157_Env -------------------- -------------------- -------------------- -------------------- --------------------
 r17080_Day80TI_158_Env -------------------- -------------------- -------------------- -------------------- --------------------
 r17080_Day80TI_159_Env -------------------- -------------------- -------------------- -------------------- --------------------
 r17080_Day80TI_160_Env -------------------- -------------------- -------------------- -------------------- --------------------
 r17080_Day80TI_161_Env -------------------- -------------------- -------------------- -------------------- --------------------
 r17080_Day80TI_162_Env -------------------- -------------------- -------------------- -------------------- --------------------
 r17080_Day80TI_163_Env -------------------- -------------------- -------------------- -------------------- --------------------
 r17080_Day80TI_164_Env -------------------- -------------------- -------------------- -------------------- --------------------


             MAC239_Env LNKYYNLTMKCRRPGNKTVL PVTIMSGLVFHSQPINDRPK QAWCWFGGKWKDAIKEVKQT IVKHPRYTGTNNTDKINLTA PGGGDPEVTFMWTNCRGEFL  400
     r16080_Day84_1_Env -------------------- -------------------- -------------------- -------------------- -R------------------
     r16080_Day84_2_Env -------------------- -------------------- -------------------- -------------------- --------------------
     r16080_Day84_3_Env -------------------- -------------------- -------------------- -------------------- -R------------------
     r16080_Day84_4_Env -------------------- -------------------- -------------------- -------------------- -R------------------
     r16080_Day84_5_Env -------------------- -------------------- ----------R--------- -------------------- -R------------------
     r16080_Day84_6_Env -------------------- -------------------- -------------------- -------N------------ --------------------
     r16080_Day84_7_Env ---------R---------- -------------------- -------------------- -------N------------ -R------------------
     r16080_Day84_8_Env -------------------- -------------------- -------------------- -------------------- --------------------
     r16080_Day84_9_Env -------------------- -------------------- -------------------- -------N------------ -R------------------
    r16080_Day84_10_Env -------------------- -------------------- -------------------- -------------------- --------------------
    r16080_Day84_11_Env -------------------- -------------------- -------------------- -------------------- --------------------
    r16080_Day84_12_Env -------------------- -------------------- -------------------- -------N------------ --------------------
    r16080_Day84_13_Env -------------------- -------------------- -------------------- -------------------- -R------------------
    r16080_Day84_14_Env -------------------- -------------------- -------------------- -------N------------ --------------------
    r16080_Day84_15_Env -------------------- -------------------- -------------------- -------N------------ -R------------------
 r17009_Day124TI_32_Env -------------------- -------------------- -------------------- -------------------- --------------------
 r17009_Day124TI_33_Env -------------------- -------------------- -------------------- -------------------- --------------------
 r17009_Day124TI_34_Env -------------------- -------------------- -------------------- -------------------- --------------------
 r17009_Day124TI_35_Env -------------------- -------------------- -------------------- -------------------- ----N---------------
 r17009_Day124TI_36_Env -------------------- -------------------- -------------------- -------------------- --------------------
 r17009_Day124TI_37_Env -------------------- -------------------- -------------------- -------------------- --------------------
 r17009_Day124TI_38_Env -------------------- -------------------- -------------------- -------------------- --------------------
 r17009_Day124TI_39_Env -------------------- -------------------- -------------------- -------------------- --------------------
 r17009_Day124TI_40_Env -------------------- -------------------- -------------------- -------------------- --------------------
 r17009_Day124TI_41_Env -------------------- -------------------- -------------------- -------------------- --------------------
 r17009_Day124TI_42_Env -------------------- -------------------- -------------------- -------------------- --------------------
 r17009_Day124TI_43_Env -------------------- -------------------- -------------------- -------------------- --------------------
 r17009_Day124TI_44_Env -------------------- -------------------- -------------------- -------------------- --------------------
 r17009_Day124TI_45_Env -------------------- -------------------- -------------------- -------------------- --------------------
 r17009_Day124TI_46_Env -------------------- -------------------- -------------------- -------------------- --------------------
 r17009_Day124TI_47_Env -------------------- -------------------- -------------------- -------------------- --------------------
 r17009_Day124TI_48_Env -------------------- -------------------- -------------------- -------------------- --------------------
 r17009_Day124TI_49_Env -------------------- -------------------- -------------------- -------------------- --------------------
 r17009_Day124TI_50_Env -------------------- -------------------- -------------------- -------------------- --------------------
 r17009_Day124TI_51_Env -------------------- -------------------- -------------------- -------------------- --------------------
 r17009_Day124TI_52_Env -------------------- -------------------- -------------------- -------------------- --------------------
 r17009_Day124TI_53_Env -------------------- -------------------- -------------------- -------------------- --------------------
 r17009_Day124TI_54_Env -------------------- -------------------- -------------------- -------------------- --------------------
 r17009_Day124TI_55_Env -------------------- -------------------- -------------------- -------------------- --------------------
 r17009_Day124TI_56_Env -------------------- -------------------- -------------------- -------------------- --------------------
 r17009_Day124TI_57_Env -------------------- -------------------- -------------------- -------------------- --------------------
r17009_Day118TI_165_Env -------------------- -------------------- -------------------- -------------------- --------------------
r17009_Day118TI_166_Env -------------------- -------------------- -------------------- -------------------- --------------------
r17009_Day118TI_167_Env -------------------- -------------------- -------------------- -------------------- --------------------
r17009_Day118TI_168_Env -------------------- -------------------- -------------------- -------------------- --------------------
r17009_Day118TI_169_Env -------------------- -------------------- -------------------- -------------------- --------------------
r17009_Day118TI_170_Env -------------------- -------------------- -------------------- -------------------- --------------------
r17009_Day118TI_171_Env -------------------- -------------------- ---------*---------- -------------------- --------------------
r17009_Day118TI_172_Env -------------------- -------------------- -------------------- -------------------- --------------------
r17009_Day118TI_173_Env -------------------- -------------------- -------------------- -------------------- --------------------
  r17058_Day84TI_59_Env -------------------- -------------------- -------------------- -------------------- --------------------
  r17058_Day84TI_60_Env -------------------- -------------------- -------------------- -------------------- --------------------
  r17058_Day84TI_61_Env -------------------- -------------------- -------------------- -------------------- --------------------
  r17058_Day84TI_62_Env -------------------- -------------------- -------------------- -------------------- --------------------
  r17058_Day84TI_63_Env -------------------- -------------------- -------------------- -------------------- --------------------
  r17058_Day84TI_64_Env -------------------- -------------------- -------------------- -------------------- --------------------
  r17058_Day84TI_65_Env -------------------- --------------V----- -------------------- -------------------- --------------------
  r17058_Day84TI_66_Env -------------------- -------------------- -------------------- -------------------- --------------------
  r17058_Day84TI_67_Env -------------------- -------------------- -------------------- -------------------- --------------------
  r17058_Day84TI_68_Env -------------------- -------------------- -------------------- -------------------- -E------------------
  r17058_Day84TI_69_Env -------------------- -------------------- -------------------- -------------------- --------------------
  r17058_Day84TI_70_Env -------------------- -------------------- -------------------- -------------------- --------------------
  r17058_Day84TI_71_Env -------------------- -------------------- -------------------- -------------------- --------------------
  r17058_Day84TI_72_Env -------------------- -------------------- -------------------- -------------------- --------------------
  r17058_Day84TI_73_Env -------------------- -------------------- -------------------- -------------------- --------------------
  r17080_Day73TI_88_Env -------------------- -------------------- -------------------- -------------------- --------------------
  r17080_Day73TI_89_Env -------------------- -------------------- -------------------- -------------------- --------------------
  r17080_Day73TI_90_Env -------------------- -------------------- -------------------- -------------------- --------------------
  r17080_Day73TI_91_Env -------------------- -------------------- -------------------- -------------------- --------------------
  r17080_Day73TI_92_Env -------------------- -------------------- -------------------- -------------------- --------------------
  r17080_Day73TI_93_Env -------------------- -------------------- -------------------- -------------------- --------------------
  r17080_Day73TI_94_Env -------------------- -------------------- -------------------- -------------------- --------------------
  r17080_Day73TI_95_Env -------------------- -------------------- -------------------- -------------------- --------------------
  r17080_Day73TI_96_Env -------------------- -------------------- -------------------- -------------------- --------------------
  r17080_Day73TI_97_Env -------------------- -------------------- -------------------- -------------------- --------------------
  r17080_Day73TI_98_Env -------------------- -------------------- -------------------- -------------------- --------------------
  r17080_Day73TI_99_Env -------------------- -------------------- -------------------- -------------------- --------------------
 r17080_Day73TI_100_Env -------------------- -------------------- -------------------- -------------------- --------------------
 r17080_Day73TI_101_Env -------------------- -------------------- -------------------- -------------------- --------------------
 r17080_Day73TI_102_Env -------------------- -------------------- -------------------- -------------------- --------------------
 r17080_Day73TI_103_Env -------------------- -------------------- -------------------- -------------------- --------------------
 r17080_Day73TI_104_Env -------------------- -------------------- -------------------- -------------------- --------------------
 r17080_Day73TI_105_Env -------------------- -------------------- -------------------- -------------------- --------------------
 r17080_Day73TI_106_Env -------------------- -------------------- -------------------- -------------------- --------------------
 r17080_Day73TI_107_Env -------------------- -------------------- -------------------- -------------------- --------------------
 r17080_Day73TI_108_Env -------------------- -------------------- -------------------- -------------------- --------------------
 r17080_Day73TI_109_Env -------------------- -------------------- -------------------- -------------------- --------------------
 r17080_Day73TI_110_Env -------------------- -------------------- -------------------- -------------------- --------------------
 r17080_Day77TI_111_Env -------------------- -------------------- -------------------- -------------------- --------------------
 r17080_Day77TI_112_Env -------------------- -------------------- -------------------- -----------...------ --------------------
 r17080_Day77TI_113_Env -------------------- -------------------- -------------------- -------------------- --------------------
 r17080_Day77TI_114_Env -------------------- -------------------- -------------------- -------------------- --------------------
 r17080_Day77TI_115_Env -------------------- -------------------- -------------------- -------------------- --------------------
 r17080_Day77TI_116_Env -------------------- -------------------- -------------------- -------------------- --------------------
 r17080_Day77TI_117_Env -------------------- -------------------- -------------------- -------------------- --------------------
 r17080_Day77TI_118_Env -------------------- -------------------- -------------------- -------------------- --------------------
 r17080_Day77TI_119_Env -------------------- -------------------- -------------------- -------------------- --------------------
 r17080_Day77TI_120_Env -------------------- -------------------- -------------------- -------------------- --------------------
 r17080_Day77TI_121_Env -------------------- -------------------- -------------------- -------------------- --------------------
 r17080_Day77TI_122_Env -------------------- -------------------- -------------------- -------------------- --------------------
 r17080_Day77TI_123_Env -------------------- -------------------- -------------------- -------------------- --------------------
 r17080_Day77TI_124_Env -------------------- -------------------- -------------------- -------------------- --------------------
 r17080_Day77TI_125_Env -------------------- -------------------- -------------------- -------------------- --------------------
 r17080_Day77TI_126_Env -------------------- -------------------- -------------------- -------------------- --------------------
 r17080_Day77TI_127_Env -------------------- -------------------- -------------------- -------------------- --------------------
 r17080_Day77TI_128_Env -------------------- -------------------- -------------------- -------------------- --------------------
 r17080_Day77TI_129_Env -------------------- -------------------- -------------------- -------------------- --------------------
 r17080_Day77TI_130_Env -------------------- -------------------- -------------------- -------------------- --------------------
 r17080_Day77TI_131_Env -------------------- -------------------- -------------------- -------------------- --------------------
 r17080_Day77TI_132_Env -------------------- -------------------- -------------------- -------------------- --------------------
 r17080_Day77TI_133_Env -------------------- -------------------- -------------------- -------------------- --------------------
 r17080_Day77TI_134_Env -------------------- -------------------- -------------------- -------------------- --------------------
 r17080_Day77TI_135_Env -------------------- -------------------- -------------------- -------------------- --------------------
 r17080_Day77TI_136_Env -------------------- -------------------- -------------------- -------------------- --------------------
 r17080_Day77TI_137_Env -------------------- -------------------- -------------------- -------------------- --------------------
 r17080_Day77TI_138_Env -------------------- -------------------- -------------------- -------------------- --------------------
 r17080_Day77TI_139_Env -------------------- -------------------- -------------------- -------------------- --------------------
    r16080_Day89_16_Env -------------------- -------------------- -------------------- -------------------- -R------------------
    r16080_Day89_17_Env -------------------- -------------------- -------------------- -------------------- -R------------------
    r16080_Day89_18_Env -------------------- -------------------- -------------------- -------------------- -R------------------
    r16080_Day89_19_Env -------------------- -------------------- -------------------- -------------------- -R------------------
    r16080_Day89_20_Env -------------------- -------------------- -------------------- -------------------- -R------------------
    r16080_Day89_21_Env -------------------- -------------------- -------------------- -------------------- -R------------------
    r16080_Day89_22_Env -------------------- -------------------- -------------------- -------------------- -R------------------
    r16080_Day89_23_Env -------------------- -------------------- -------------------- -------------------- -R------------------
    r16080_Day89_24_Env -------------------- -------------------- -------------------- -------------------- -R------------------
    r16080_Day89_25_Env -------------------- -------------------- -------------------- -------------------- --------------------
    r16080_Day89_26_Env -------------------- -------------------- -------------------- -------------------- -R------------------
    r16080_Day89_27_Env -------------------- -------------------- -------------------- -------------------- -R------------------
    r16080_Day89_28_Env -------------------- -------------------- -------------------- -------------------- -R------------------
    r16080_Day89_29_Env -------------------- -------------------- -------------------- -------------------- --------------------
    r16080_Day89_30_Env -------------------- -------------------- -------------------- -------------------- -R------------------
    r16080_Day89_31_Env -------------------- -------------------- -------------------- -------------------- -R------------------
r17009_Day131TI_175_Env -------------------- -------------------- -------------------- -------------------- --------------------
r17009_Day131TI_176_Env -------------------- -------------------- -------------------- -------------------- --------------------
r17009_Day131TI_177_Env -------------------- -------------------- -------------------- -------------------- --------------------
r17009_Day131TI_178_Env -------------------- -------------------- -------------------- -------------------- --------------------
r17009_Day131TI_179_Env -------------------- -------------------- -------------------- -------------------- --------------------
r17009_Day131TI_180_Env -------------------- -------------------- -------------------- -------------------- --------------------
r17009_Day131TI_181_Env -------------------- -------------------- -------------------- -------------------- --------------------
r17009_Day131TI_182_Env -------------------- -------------------- -------------------- -------------------- --------------------
r17009_Day131TI_183_Env -------------------- -------------------- -------------------- -------------------- --------------------
r17009_Day131TI_184_Env -------------------- -------------------- -------------------- -------------------- --------------------
r17009_Day131TI_185_Env -------------------- -------------------- -------------------- -------------------- --------------------
r17009_Day131TI_186_Env -------------------- -------------------- -------------------- -------------------- --------------------
r17009_Day131TI_187_Env -------------------- -------------------- -------------------- -------------------- --------------------
r17009_Day131TI_188_Env -------------------- -------------------- -------------------- -------------------- --------------------
r17009_Day131TI_189_Env -------------------- -------------------- -------------------- -------------------- --------------------
r17009_Day131TI_190_Env -------------------- -------------------- -------------------- -------------------- --------------------
r17009_Day131TI_191_Env -------------------- -------------------- -------------------- -------------------- --------------------
r17009_Day131TI_192_Env -------------------- -------------------- -------------------- -------------------- --------------------
r17009_Day131TI_193_Env -------------------- -------------------- -------------------- -------------------- --------------------
r17009_Day131TI_194_Env -------------------- -------------------- -------E------------ -------------------- --------------------
r17009_Day131TI_195_Env -------------------- -------------------- -------------------- -------------------- --------------------
r17009_Day131TI_196_Env -------------------- -------------------- -------------------- -------------------- --------------------
r17009_Day131TI_197_Env -------------------- -------------------- -------------------- -------------------- --------------------
r17009_Day131TI_198_Env -------------------- -------------------- -------------------- -------------------- --------------------
r17009_Day131TI_199_Env -------------------- -------------------- -------------------- -------------------- --------------------
r17009_Day131TI_200_Env -------------------- -------------------- -------------------- -------------------- --------------------
r17009_Day131TI_201_Env -------------------- -------------------- -------------------- -------------------- --------------------
r17009_Day131TI_202_Env -------------------- -------------------- -------------------- -------------------- --------------------
 r17058__Day89TI_74_Env -------------------- -------------------R -------------------- -------------------- --------------------
 r17058__Day89TI_75_Env -------------------- -------------------- -------------------- -------------------- --------------------
 r17058__Day89TI_76_Env -------------------- -------------------- -------------------- -------------------- --------------------
 r17058__Day89TI_77_Env -------------------- -------------------- -------------------- -------------------- --------------------
 r17058__Day89TI_78_Env -------------------- -------------------- -------------------- -------------------- --------------------
 r17058__Day89TI_79_Env -------------------- -------------------- -------------------- -------------------- --------------------
 r17058__Day89TI_80_Env -------------------- -------------------- -------------------- -------------------- --------------------
 r17058__Day89TI_81_Env -------------------- -------------------- -------------------- -------------------- --------------------
 r17058__Day89TI_82_Env -------------------- -------------------- -------------------- -------------------- --------------------
 r17058__Day89TI_83_Env -------------------- -------------------- -------------------- -------------------- --------------------
 r17058__Day89TI_84_Env -------------------- -------------------- -------------------- -------------------- --------------------
 r17058__Day89TI_85_Env -------------------- -------------------- -------------------- -------------------- --------------------
 r17058__Day89TI_86_Env -------------------- -------------------- -------------------- -------------------- --------------------
 r17058__Day89TI_87_Env -------------------- -------------------- -------------------- -------------------- --------------------
 r17080_Day80TI_140_Env -------------------- -------------------- -------------------- -------------------- --------------------
 r17080_Day80TI_141_Env -------------------- -------------------- -------------------- -------------------- --------------------
 r17080_Day80TI_142_Env -------------------- -------------------- -------------------- -------------------- --------------------
 r17080_Day80TI_143_Env -------------------- -------------------- -------------------- -------------------- --------------------
 r17080_Day80TI_144_Env -------------------- -------------------- -------------------- -------------------- --------------------
 r17080_Day80TI_145_Env -------------------- -------------------- -------------------- -------------------- --------------------
 r17080_Day80TI_146_Env -------------------- -------------------- -------------------- -------------------- --------------------
 r17080_Day80TI_147_Env -------------------- -------------------- -------------------- -------------------- --------------------
 r17080_Day80TI_148_Env -------------------- -------------------- -------------------- -------------------- --------------------
 r17080_Day80TI_149_Env -------------------- -------------------- -------------------- -------------------- --------------------
 r17080_Day80TI_150_Env -------------------- -------------------- -------------------- -------------------- --------------------
 r17080_Day80TI_151_Env -------------------- -------------------- -------------------- -------------------- --------------------
 r17080_Day80TI_152_Env -------------------- -------------------- -------------------- -------------------- --------------------
 r17080_Day80TI_153_Env -------------------- -------------------- -------------------- -------------------- --------------------
 r17080_Day80TI_154_Env -------------------- -------------------- -------------------- -------------------- --------------------
 r17080_Day80TI_155_Env -------------------- -------------------- -------------------- -------------------- --------------------
 r17080_Day80TI_156_Env -------------------- -------------------- -------------------- -------------------- --------------------
 r17080_Day80TI_157_Env -------------------- -------------------- -------------------- -------------------- --------------------
 r17080_Day80TI_158_Env -------------------- -------------------- -------------------- -------------------- --------------------
 r17080_Day80TI_159_Env -------------------- -------------------- -------------------- -------------------- --------------------
 r17080_Day80TI_160_Env -------------------- -------------------- -------------------- -------------------- --------------------
 r17080_Day80TI_161_Env -------------------- -------------------- -------------------- -------------------- --------------------
 r17080_Day80TI_162_Env -------------------- -------------------- -------------------- -------------------- --------------------
 r17080_Day80TI_163_Env -------------------- -------------------- -------------------- -------------------- --------------------
 r17080_Day80TI_164_Env -------------------- -------------------R -------------------- -------------------- --------------------

             MAC239_Env YCKMNWFLNWVEDRNTANQK PKEQHKRNYVPCHIRQIINT WHKVGKNVYLPPREGDLTCN STVTSLIANIDWIDGNQTNI TMSAEVAELYRLELGDYKLV  500
     r16080_Day84_1_Env -------------------- -------------------- -------------------- ------------------D- --------------------
     r16080_Day84_2_Env -------------------- -R------------------ -------------------- ------------------D- --------------------
     r16080_Day84_3_Env -------------------- -------------------- -------------------- ------------------D- --------------------
     r16080_Day84_4_Env -------------------- -------------------- -------------------- ------------------D- --------------------
     r16080_Day84_5_Env -------------------- -------------------- -------------------- ------------------D- --------------------
     r16080_Day84_6_Env -------------------- -------------------- -------------------- -------------------- --------------------
     r16080_Day84_7_Env -------------------- -------------------- -------------------- -------------------- --------------------
     r16080_Day84_8_Env -------------------- -E------------------ -------------------- ------------------D- --------------------
     r16080_Day84_9_Env -------------------- -------------------- -------------------- -------------------- --------------------
    r16080_Day84_10_Env -------------------- -------------------- -------------------- ------------------D- --------------------
    r16080_Day84_11_Env -------------------- -------------------- -------------------- ------------------D- --------------------
    r16080_Day84_12_Env -------------------- -------------------- -------------------- -------------------- --------------------
    r16080_Day84_13_Env -------------------- -------------------- -------------------- ------------------D- --------------------
    r16080_Day84_14_Env -------------------- -------------------- -------------------- -------------------- --------------------
    r16080_Day84_15_Env -------------------- -------------------- -------------------- -------------------- --------------------
 r17009_Day124TI_32_Env -------------------- -------------------- -------------------- ------------------K- ------------K-------
 r17009_Day124TI_33_Env -------------------- -------------------- -------------------- ------------------K- --------------------
 r17009_Day124TI_34_Env -------------------- -------------------- -------------------- ------------------K- --------------------
 r17009_Day124TI_35_Env -------------------- -------------------- -------------------- ------------------K- --------------------
 r17009_Day124TI_36_Env -------------------- -------------------- -------------------- ------------------K- --------------------
 r17009_Day124TI_37_Env -------------------- -------------------- -------------------- ------------------K- --------------------
 r17009_Day124TI_38_Env -------------------- -------------------- -------------------- ------------------K- --------------------
 r17009_Day124TI_39_Env -------------------- -------------------- -------------------- ------------------K- --------------------
 r17009_Day124TI_40_Env -------------------- -------------------- -------------------- ------------------K- --------------------
 r17009_Day124TI_41_Env -------------------- -------------------- -------------------- ------------------K- --------------------
 r17009_Day124TI_42_Env -------------------- -------------------- -------------------- ------------------K- --------------------
 r17009_Day124TI_43_Env -------------------- -------------------- -------------------- ------------------K- --------------------
 r17009_Day124TI_44_Env -------------------- -------------------- -------------------- ------------------K- --------------------
 r17009_Day124TI_45_Env -------------------- -------------------- -------------------- ------------------K- --------------------
 r17009_Day124TI_46_Env -------------------- -------------------- -------------------- ------------------K- --------------------
 r17009_Day124TI_47_Env -------------------- -------------------- -------------------- ------------------K- --------------------
 r17009_Day124TI_48_Env -------------------- -------------------- -------------------- ------------------K- --------------------
 r17009_Day124TI_49_Env -------------------- -------------------- -------------------- ------------------K- --------------------
 r17009_Day124TI_50_Env -------------------- -------------------- -------------------- ------------------K- --------------------
 r17009_Day124TI_51_Env -------------------- -------------------- -------------------- ------------------K- --------------------
 r17009_Day124TI_52_Env -------------------- -------------------- -------------------- ------------------K- --------------------
 r17009_Day124TI_53_Env -------------------- -------------------- -------------------- ------------------K- --------------------
 r17009_Day124TI_54_Env -------------------- -------------------- -------------------- ------------------K- --------------------
 r17009_Day124TI_55_Env -------------------- -------------------- -------------------- ------------------K- --------------------
 r17009_Day124TI_56_Env -------------------- -------------------- -------------------- ------------------K- --------------------
 r17009_Day124TI_57_Env -------------------- -------------------- -------------------- ------------------K- --------------------
r17009_Day118TI_165_Env -------------------- -------------------- -------------------- ------------------K- --------------------
r17009_Day118TI_166_Env -------------------- -------------------- -------------------- ------------------K- --------------------
r17009_Day118TI_167_Env -------------------- -------------------- -------------------- ------------------K- --------------------
r17009_Day118TI_168_Env -------------------- -------------------- -------------------- ------------------K- --------------------
r17009_Day118TI_169_Env -------------------- -------------------- -------------------- ------------------K- --------------------
r17009_Day118TI_170_Env -------------------- -------------------- -------------------- ------------------K- --------------------
r17009_Day118TI_171_Env -------------------- -------------------- -------------------- ------------------K- --------------------
r17009_Day118TI_172_Env -------------------- -------------------- -------------------- ------------------K- --------------------
r17009_Day118TI_173_Env -------------------- -------------------- -------------------- ------------------K- --------------------
  r17058_Day84TI_59_Env -------------------- -------------------- -------------------- -------------------- --------------------
  r17058_Day84TI_60_Env -------------------- -------------------- -------------------- -------------------- --------------------
  r17058_Day84TI_61_Env -------------------- -------------------- -------------------- -----------R-------- --------------------
  r17058_Day84TI_62_Env -------------------- -------------------- -------------------- -------------------- --------------------
  r17058_Day84TI_63_Env -------------------- -------------------- -------------------- -------------------- --------------------
  r17058_Day84TI_64_Env -------------------- -------------------- -------------------- -------------------- --------------------
  r17058_Day84TI_65_Env -------------------- -------------------- -------------------- -------------------- --------------------
  r17058_Day84TI_66_Env -------------------- -------------------- -------------------- -------------------- --------------------
  r17058_Day84TI_67_Env -------------------- -------------------- -------------------- -------------------- --------------------
  r17058_Day84TI_68_Env -------------------- -------------------- -------------------- -------------------- --------------------
  r17058_Day84TI_69_Env -------------------- -------------------- -------------------- -------------------- --------------------
  r17058_Day84TI_70_Env -------------------- -------------------- -------------------- -------------------- --------------------
  r17058_Day84TI_71_Env -------------------- ----------------V--- -------------------- -------------------- --------------------
  r17058_Day84TI_72_Env -------------------- ----Q--------------- -------------------- -------------------- --------------------
  r17058_Day84TI_73_Env -------------------- -------------------- -------------------- -------------------- --------------------
  r17080_Day73TI_88_Env -------------------- -------------------- -------------------- -------------------- A-------------------
  r17080_Day73TI_89_Env -------------------- -------------------- -------------------- -------------------- A-------------------
  r17080_Day73TI_90_Env -------------------- -------------------- -------------------- -------------------- A-------------------
  r17080_Day73TI_91_Env -------------------- -------------------- -------------------- -------------------- A-------------------
  r17080_Day73TI_92_Env -------------------- -------------------- -------------------- -------------------- A-------------------
  r17080_Day73TI_93_Env -------------------- -------------------- -------------------- -------------------- A-------------------
  r17080_Day73TI_94_Env -------------------- -------------------- -------------------- -------------------- A-------------------
  r17080_Day73TI_95_Env -------------------- -------------------- -------------------- -------------------- A-------------------
  r17080_Day73TI_96_Env -------------------- -------------------- -------------------- -------------------- A-------------------
  r17080_Day73TI_97_Env -------------------- -------------------- -------------------- -------------------- A-------------------
  r17080_Day73TI_98_Env -------------------- -------------------- -------------------- -------------------- A-------------------
  r17080_Day73TI_99_Env -------------------- -------------------- -------------------- -------------------- A-------------------
 r17080_Day73TI_100_Env -------------------- -------------------- -------------------- -------------------- A-------------------
 r17080_Day73TI_101_Env -------------------- -------------------- -------------------- -------------------- A-------------------
 r17080_Day73TI_102_Env -------------------- -------------------- -------------------- -------------------- A-------------------
 r17080_Day73TI_103_Env -------------------- -------------------- -------------------- -------------------- A-------------------
 r17080_Day73TI_104_Env -------------------- -------------------- -------------------- -------------------- A-------------------
 r17080_Day73TI_105_Env -------------------- -------------------- -------------------- -------------------- A-------------------
 r17080_Day73TI_106_Env -------------------- -------------------- -------------------- -------------------- A-------------------
 r17080_Day73TI_107_Env -------------------- -------------------- -------------------- -------------------- A-------------R-----
 r17080_Day73TI_108_Env -------------------- -------------------- -------------------- -------------------- A-------------------
 r17080_Day73TI_109_Env -------------------- -------------------- -------------------- -------------------- A-------------------
 r17080_Day73TI_110_Env -------------------- -------------------- -------------------- -------------------- A-------------------
 r17080_Day77TI_111_Env -------------------- -------------------- -------------------- -------------------- A-------------------
 r17080_Day77TI_112_Env -------------------- -------------------- -------------------- -------------------- A-------------------
 r17080_Day77TI_113_Env -------------------- -------------------- -------------------- -------------------- A-------------------
 r17080_Day77TI_114_Env -------------------- -------------------- -------------------- -------------------- A-------------------
 r17080_Day77TI_115_Env -------------------- -------------------- -------------------- -------------------- A-------------------
 r17080_Day77TI_116_Env -------------------- -------------------- -------------------- -------------------- A-------------------
 r17080_Day77TI_117_Env -------------------- -------------------- -------------------- -------------------- A-------------------
 r17080_Day77TI_118_Env -------------------- -------------------- -------------------- ------------------D- --------------------
 r17080_Day77TI_119_Env -------------------- -------------------- -------------------- -------------------- A-------------------
 r17080_Day77TI_120_Env -------------------- -------------------- -------------------- -------------------- A-------------------
 r17080_Day77TI_121_Env -------------------- -------------------- -------------------- -------------------- A-------------------
 r17080_Day77TI_122_Env -------------------- -------------------- -------------------- -------------------- A-------------------
 r17080_Day77TI_123_Env -------------------- -------------------- ------------K------- -------------------- A-------------------
 r17080_Day77TI_124_Env -------------------- -------------------- -------------------- -------------------- A-------------------
 r17080_Day77TI_125_Env -------------------- -------------------- -------------------- -------------------- A-------------------
 r17080_Day77TI_126_Env -------------------- -------------------- -------------------- -------------------- A-------------------
 r17080_Day77TI_127_Env -------------------- -------------------- -------------------- -------------------- A-------------------
 r17080_Day77TI_128_Env -------------------- -------------------- -------------------- -------------------- A-------------------
 r17080_Day77TI_129_Env -------------------- -------------------- -------------------- -------------------- A-------------------
 r17080_Day77TI_130_Env -------------------- -------------------- -------------------- ------------------D- --------------------
 r17080_Day77TI_131_Env -------------------- -------------------- -------------------- -------------------- A-------------------
 r17080_Day77TI_132_Env -------------------- ----------------.--- -------------------- -------------------- A-------------------
 r17080_Day77TI_133_Env -------------------- -------------------- -------------------- -------------------- A-------------------
 r17080_Day77TI_134_Env -------------------- -------------------- -------------------- -------------------- A-------------------
 r17080_Day77TI_135_Env -------------------- -------------------- -------------------- ------------------D- --------------------
 r17080_Day77TI_136_Env -------------------- -------------------- -------------------- -------------------- A-------------------
 r17080_Day77TI_137_Env -------------------- -------------------- -------------------- -------------------- A-------------------
 r17080_Day77TI_138_Env -------------------- -------------------- -------------------- -------------------- A-------------------
 r17080_Day77TI_139_Env -------------------- -------------------- -------------------- -------------------- A-------------------
    r16080_Day89_16_Env -------------------- -------------------- -------------------- ------------------D- --------------------
    r16080_Day89_17_Env -------------------- -------------------- -------------------- ------------------D- --------------------
    r16080_Day89_18_Env -------------------- -------------------- -------------------- ------------------D- --------------------
    r16080_Day89_19_Env -------------------- -------------------- -------------------- ------------------D- --------------------
    r16080_Day89_20_Env -------------------- -------------------- -------------------- ------------------D- --------------------
    r16080_Day89_21_Env -------------------- -------------------- -------------------- ------------------D- --------------------
    r16080_Day89_22_Env -------------------- -------------------- -------------------- ------------------D- --------------------
    r16080_Day89_23_Env -------------------- -------------------- -------------------- ------------------D- --------------------
    r16080_Day89_24_Env -------------------- -------------------- -------------------- ------------------D- --------------------
    r16080_Day89_25_Env -------------------- -------------------- -------------------- ------------------D- --------------------
    r16080_Day89_26_Env -------------------- -------------------- -------------------- ------------------D- --------------------
    r16080_Day89_27_Env -------------------- -------------------- -------------------- ------------------D- --------------------
    r16080_Day89_28_Env -------------------- -------------------- -----R-------------- ------------------D- --------------------
    r16080_Day89_29_Env -------------------- -E------------------ -------------------- ------------------G- --------------------
    r16080_Day89_30_Env -------------------- -------------------- -------------------- ------------------D- --------------------
    r16080_Day89_31_Env -------------------- -------------------- -------------------- ------------------D- --------------------
r17009_Day131TI_175_Env -------------------- -------------------- *------------------- ------------------K- --------------------
r17009_Day131TI_176_Env -------------------- -------------------- -------------------- ------------------K- --------------------
r17009_Day131TI_177_Env -------------------- -------------------- -------------------- ------------------K- --------------------
r17009_Day131TI_178_Env -------------------- -------------------- -------------------- ------------------K- --------------------
r17009_Day131TI_179_Env -------------------- -------------------- -------------------- ------------------K- --------------------
r17009_Day131TI_180_Env -------------------- -------------------- -------------------- ------------------K- --------------------
r17009_Day131TI_181_Env -------------------- -------------------- -------------------- ------------------K- --------------------
r17009_Day131TI_182_Env -------------------- -------------------- -------------------- ------------------K- --------------------
r17009_Day131TI_183_Env -------------------- -------------------- -------------------- ------------------K- --------------------
r17009_Day131TI_184_Env -------------------- -------------------- -------------------- ------------------K- --------------------
r17009_Day131TI_185_Env -------------------- -------------------- -------------------- ------------------K- --------------------
r17009_Day131TI_186_Env -------------------- -------------------- -------------------- ------------------K- --------------------
r17009_Day131TI_187_Env -------------------- -------------------- -------------------- ------------------K- --------------------
r17009_Day131TI_188_Env -------------------- -------------------- -------------------- ------------------K- --------------------
r17009_Day131TI_189_Env -------------------- -------------------- -------------------- ------------------K- --------------------
r17009_Day131TI_190_Env -------------------- -------------------- -------------------- ------------------K- --------------------
r17009_Day131TI_191_Env -------------------- -------------------- -------------------- ------------------K- --------------------
r17009_Day131TI_192_Env -------------------- -------------------- -------------------- ------------------K- --------------------
r17009_Day131TI_193_Env -------------------- -------------------- -------------------- ------------------K- --------------------
r17009_Day131TI_194_Env -------------------- -------------------- -------------------- ------------------K- --------------------
r17009_Day131TI_195_Env -------------------- -------------------- -------------------- ------------------K- --------------------
r17009_Day131TI_196_Env -------------------- -------------------- -------------------- ------------------K- --------------------
r17009_Day131TI_197_Env -------------------- -------------------- -------------------- ------------------K- --------------------
r17009_Day131TI_198_Env -------------------- -------------------- -------------------- ------------------K- --------------------
r17009_Day131TI_199_Env -------------------- -------------------- -------------------- ------------------K- --------------------
r17009_Day131TI_200_Env -------------------- -------------------- -------------------- ------------------K- --------------------
r17009_Day131TI_201_Env -------------------- -------------------- -------------------- ------------------K- --------------------
r17009_Day131TI_202_Env -------------------- -------------------- -------------------- ------------------K- --------------------
 r17058__Day89TI_74_Env -------------------- -------------------- -------------------- -------------------- --------------------
 r17058__Day89TI_75_Env -------------------- -------------------- -------------------- -------------------- --------------------
 r17058__Day89TI_76_Env -------------------- -------------------- -------------------- -------------------- --------------------
 r17058__Day89TI_77_Env -------------------- -------------------- -------------------- -------------------- --------------------
 r17058__Day89TI_78_Env -------------------- -------------------- -------------------- -------------------- --------------------
 r17058__Day89TI_79_Env -------------------- -------------------- -------------------- -------------------- --------------------
 r17058__Day89TI_80_Env -------------------- -------------------- -------------------- -------------------- --------------------
 r17058__Day89TI_81_Env -------------------- -------------------- -------------------- -------------------- --------------------
 r17058__Day89TI_82_Env -------------------- -------------------- -------------------- -------------------- --------------------
 r17058__Day89TI_83_Env -------------------- -------------------- -------------------- ---S---------------- --------------------
 r17058__Day89TI_84_Env -------------------- -------------------- -------------------- -------------------- --------------------
 r17058__Day89TI_85_Env -------------------- -------------------- -------------------- -------------------- --------------------
 r17058__Day89TI_86_Env -------------------- -------------------- -------------------- -------------------- --------------------
 r17058__Day89TI_87_Env -------------------- -------------------- -------------------- -------------------- --------------------
 r17080_Day80TI_140_Env -------------------- -------------------- -------------------- -------------------- A-------------------
 r17080_Day80TI_141_Env -------------------- -------------------- -------------------- -------------------- A-------------------
 r17080_Day80TI_142_Env -------------------- -------------------- -------------------- -------------------- A-------------------
 r17080_Day80TI_143_Env -------------------- -------------------- -------------------- -------------------- A-------------------
 r17080_Day80TI_144_Env -------------------- -------------------- -------------------- -------------------- A-------------------
 r17080_Day80TI_145_Env -------------------- -------------------- -------------------- -------------------- A-------------------
 r17080_Day80TI_146_Env -------------------- -------------------- -------------------- -------------------- A-------------------
 r17080_Day80TI_147_Env -------------------- -------------------- -------------------- -------------------- A-------------------
 r17080_Day80TI_148_Env -------------------- -------------------- -------------------- -------------------- A-------------------
 r17080_Day80TI_149_Env -------------------- -------------------- -------------------- -------------------- A-------------------
 r17080_Day80TI_150_Env -------------------- -------------------- -------------------- -------------------- A-------------------
 r17080_Day80TI_151_Env -------------------- -------------------- -------------------- -------------------- A-------------------
 r17080_Day80TI_152_Env -------------------- -------------------- -------------------- -------------------- A-------------------
 r17080_Day80TI_153_Env -------------------- -------------------- -------------------- -------------------- A-------------------
 r17080_Day80TI_154_Env ----------------P--- -------------------- -------------------- -------------------- A-------------------
 r17080_Day80TI_155_Env -------------------- -------------------- -------------------- -------------------- A-------------------
 r17080_Day80TI_156_Env -------------------- -------------------- -------------------- -------------------- A-------------------
 r17080_Day80TI_157_Env -------------------- -------------------- -------------------- ------------------D- --------------------
 r17080_Day80TI_158_Env -----------K-------- -------------------- -------------------- -------------------- A-------------------
 r17080_Day80TI_159_Env -------------------- -------------------- -------------------- -------------------- A-------------------
 r17080_Day80TI_160_Env -------------------- -------------------- -------------------- -------------------- A-------------------
 r17080_Day80TI_161_Env -------------------- -------------------- -------------------- -------------------- A-------------------
 r17080_Day80TI_162_Env -------------------- -------------------- -------------------- -------------------- A-------------------
 r17080_Day80TI_163_Env -------------------- -------------------- -------------------- -------------------- A-------------------
 r17080_Day80TI_164_Env -------------------- -------------------- -------------------- -------------------- A-------------------

             MAC239_Env EITPIGLAPTDVKRYTTGGT SRNKRGVFVLGFLGFLATAG SAMGAASLTLTAQSRTLLAG IVQQQQQLLDVVKRQQELLR LTVWGTKNLQTRVTAIEKYL  600
     r16080_Day84_1_Env -------------------- -------------------- -------------------- -------------------- --------------------
     r16080_Day84_2_Env -------------------- -------------------- -------------------- -------------------- --------------------
     r16080_Day84_3_Env -------------------- ------A------------- -------------------- -------------------- --------------------
     r16080_Day84_4_Env -------------------- -------------------- -------------------- -------------------- --------------------
     r16080_Day84_5_Env -------------------- ------A------------- -------------------- -------------------- --------------------
     r16080_Day84_6_Env -------------------- -------------------- -------------------- -------------------- --------------------
     r16080_Day84_7_Env -------------------- -------------------- -------------------- -------------------- --------------------
     r16080_Day84_8_Env -------------------- --------A----------- -------------------- -------------------- --------------------
     r16080_Day84_9_Env -------------------- -------------------- -------------------- -------------------- --------------------
    r16080_Day84_10_Env -------------------- -------------------- -------------------- -------------------- --------------------
    r16080_Day84_11_Env -------------------- -------------------- -------------------- -------------------- --------------------
    r16080_Day84_12_Env -------------------- -------------------- -------------------- -------------------- --------------------
    r16080_Day84_13_Env -------------------- ------A------------- -------------------- -------------------- --------------------
    r16080_Day84_14_Env -------------------- -------------------- -------------------- -------------------- --------------------
    r16080_Day84_15_Env -------------------- -------------------- -------------------- -------------------- --------------------
 r17009_Day124TI_32_Env -------------------- -------------------- -------------------- -------------------- --------------------
 r17009_Day124TI_33_Env -------------------- -------------------- -------------------- -------------------- --------------------
 r17009_Day124TI_34_Env -------------------- -------------------- -------------------- -------------------- --------------------
 r17009_Day124TI_35_Env -------------------- -------------------- -------------------- -------------------- --------------------
 r17009_Day124TI_36_Env -------------------- -------------------- -------------------- -------------------- --------------------
 r17009_Day124TI_37_Env -------------------- -------------------- -------------------- -------------------- --------------------
 r17009_Day124TI_38_Env -------------------- -------------------- -------------------- -------------------- --------------------
 r17009_Day124TI_39_Env -------------------- -------------------- -------------------- -------------------- --------------------
 r17009_Day124TI_40_Env -------------------- -------------------- -------------------- -------------------- --------------------
 r17009_Day124TI_41_Env -------------------- -------------------- -------------------- -------------------- --------------------
 r17009_Day124TI_42_Env -------------------- -------------------- -------------------- -------------------- --------------------
 r17009_Day124TI_43_Env -------------------- -------------------- -------------------- -------------------- --------------------
 r17009_Day124TI_44_Env -------------------- -------------------- -------------------- -------------------- --------------------
 r17009_Day124TI_45_Env -------------------- -------------------- -------------------- -------------------- --------------------
 r17009_Day124TI_46_Env -------------------- -------------------- -------------------- -------------------- --------------------
 r17009_Day124TI_47_Env -------------------- -------------------- -------------------- -------------------- --------------------
 r17009_Day124TI_48_Env -------------------- -------------------- -------------------- -------------------- --------------------
 r17009_Day124TI_49_Env -------------------- -------------------- -------------------- -------------------- --------------------
 r17009_Day124TI_50_Env -------------------- -------------------- -------------------- -------------------- --------------------
 r17009_Day124TI_51_Env -------------------- -------------------- -------------------- -------------------- --------------------
 r17009_Day124TI_52_Env -------------------- -------------------- -------------------- -------------------- --------------------
 r17009_Day124TI_53_Env -------------------- -------------------- -------------------- -------------------- --------------------
 r17009_Day124TI_54_Env -------------------- -------------------- -------------------- -------------------- --------------------
 r17009_Day124TI_55_Env -------------------- -------------------- -------------------- -------------------- --------------------
 r17009_Day124TI_56_Env -------------------- -------------------- -------------------- -------------------- --------------------
 r17009_Day124TI_57_Env -------------------- -------------------- -------------------- -------------------- --------------------
r17009_Day118TI_165_Env -------------------- -------------------- -------------------- -------------------- --------------------
r17009_Day118TI_166_Env -------------------- -------------------- -------------------- -------------------- --------------------
r17009_Day118TI_167_Env -------------------- -------------------- -------------------- -------------------- --------------------
r17009_Day118TI_168_Env -------------------- -------------------- -------------------- -------------------- --------------------
r17009_Day118TI_169_Env -------------------- -------------------- -------------------- -------------------- --------------------
r17009_Day118TI_170_Env -------------------- -------------------- -------------------- -------------------- --------------------
r17009_Day118TI_171_Env -------------------- -------------------- -------------------- -------------------- --------------------
r17009_Day118TI_172_Env -------------------- -------------------- -------------------- -------------------- --------------------
r17009_Day118TI_173_Env -------------------- -------------------- -------------------- -------------------- --------------------
  r17058_Day84TI_59_Env -------------------- -------------------- -------------------- -------------------- --------------------
  r17058_Day84TI_60_Env -------------------- -----R-------------- -------------------- -------------------- --------------------
  r17058_Day84TI_61_Env -------------------- -------------------- -------------------- -------------------- --------------------
  r17058_Day84TI_62_Env -------------------- -------------------- -------------------- -------------------- --------------------
  r17058_Day84TI_63_Env -------------------- --------A----------- -------------------- -------------------- --------------------
  r17058_Day84TI_64_Env -V------------------ -------------------- -------------------- -------------------- --------------------
  r17058_Day84TI_65_Env -------------------- -------------------- -------------------- -------------------- --------------------
  r17058_Day84TI_66_Env -------------------- -------------------- -------------------- -------------------- --------------------
  r17058_Day84TI_67_Env -V------------------ -------------------- -------------------- -------------------- --------------------
  r17058_Day84TI_68_Env -------------------- -------------------- -------------------- -------------------- --------------------
  r17058_Day84TI_69_Env -------------------- -------------------- -------------------- -------------------- --------------------
  r17058_Day84TI_70_Env -------------------- --------A----------- -------------------- -------------------- --------------------
  r17058_Day84TI_71_Env -------------------- -------------------- -------------------- -------------------- --------------------
  r17058_Day84TI_72_Env -------------------- -------------------- -------------------- -------------------- --------------------
  r17058_Day84TI_73_Env -------------------- -------------------- -------------------- -------------------- --------------------
  r17080_Day73TI_88_Env -------------------- -------------------- -------------------- -------------------- --------------------
  r17080_Day73TI_89_Env -------------------- -------------------- -------------------- -------------------- --------------------
  r17080_Day73TI_90_Env -------------------- -------------------- -------------------- -------------------- --------------------
  r17080_Day73TI_91_Env -------------------- -------------------- -------------------- -------------------- --------------------
  r17080_Day73TI_92_Env -------------------- -------------------- -------------------- -------------------- --------------------
  r17080_Day73TI_93_Env -------------------- -------------------- -------------------- -------------------- --------------------
  r17080_Day73TI_94_Env -------------------- -------------------- -------------------- -------------------- --------------------
  r17080_Day73TI_95_Env -------------------- -------------------- -------------------- -------------------- --------------------
  r17080_Day73TI_96_Env -------------------- -------------------- -------------------- -------------------- --------------------
  r17080_Day73TI_97_Env -------------------- -------------------- -------------------- -------------------- --------------------
  r17080_Day73TI_98_Env -------------------- -------------------- -------------------- -------------------- --------------------
  r17080_Day73TI_99_Env -------------------- -------------------- -------------------- -------------------- --------------------
 r17080_Day73TI_100_Env -------------------- -------------------- -------------------- -------------------- --------------------
 r17080_Day73TI_101_Env -------------------- -------------------- -------------------- -------------------- --------------------
 r17080_Day73TI_102_Env -------------------- -------------------- -------------------- -------------------- --------------------
 r17080_Day73TI_103_Env -------------------- -------------------- -------------------- -------------------- --------------------
 r17080_Day73TI_104_Env -------------------- -------------------- -------------------- -------------------- --------------------
 r17080_Day73TI_105_Env -------------------- -------------------- -------------------- -------------------- --------------------
 r17080_Day73TI_106_Env -------------------- -------------------- -------------------- -------------------- --------------------
 r17080_Day73TI_107_Env -------------------- -------------------- -------------------- -------------------- --------------------
 r17080_Day73TI_108_Env -------------------- -------------------- -------------------- -------------------- --------------------
 r17080_Day73TI_109_Env -------------------- -------------------- -------------------- -------------------- --------------------
 r17080_Day73TI_110_Env -------------------- -------------------- -------------------- -------------------- --------------------
 r17080_Day77TI_111_Env -------------------- -------------------- -------------------- -------------------- --------------------
 r17080_Day77TI_112_Env -------------------- -------------------- -------------------- -------------------- --------------------
 r17080_Day77TI_113_Env -------------------- -------------------- -------------------- -------------------- --------------------
 r17080_Day77TI_114_Env -------------------- -------------------- -------------------- -------------------- --------------------
 r17080_Day77TI_115_Env -------------------- -------------------- -------------------- -------------------- --------------------
 r17080_Day77TI_116_Env -------------------- -------------------- -------------------- -------------------- --------------------
 r17080_Day77TI_117_Env -------------------- -------------------- -------------------- -------------------- --------------------
 r17080_Day77TI_118_Env -------------------- -------------------- -------------------- -------------------- --------------------
 r17080_Day77TI_119_Env -------------------- -------------------- -------------------- -------------------- --------------------
 r17080_Day77TI_120_Env -------------------- -------------------- -------------------- -------------------- --------------------
 r17080_Day77TI_121_Env -------------------- -------------------- -------------------- -------------------- --------------------
 r17080_Day77TI_122_Env -------------------- -------------------- -------------------- -------------------- --------------------
 r17080_Day77TI_123_Env -------------------- -------------------- -------------------- -------------------- --------------------
 r17080_Day77TI_124_Env -------------------- -------------------- -------------------- -------------------- --------------------
 r17080_Day77TI_125_Env -------------------- -------------------- -------------------- -------------------- --------------------
 r17080_Day77TI_126_Env -------------------- -------------------- -------------------- -------------------- --------------------
 r17080_Day77TI_127_Env -------------------- -------------------- -------------------- -------------------- --------------------
 r17080_Day77TI_128_Env -------------------- -------------------- -------------------- -------------------- --------------------
 r17080_Day77TI_129_Env -------------------- -------------------- -------------------- -------------------- --------------------
 r17080_Day77TI_130_Env -------------------- -------------------- -------------------- -------------------- --------------------
 r17080_Day77TI_131_Env -------------------- -------------------- -------------------- -------------------- --------------------
 r17080_Day77TI_132_Env -------------------- -------------------- -------------------- -------------------- --------------------
 r17080_Day77TI_133_Env -------------------- -------------------- -------------------- -------------------- --------------------
 r17080_Day77TI_134_Env -------------------- -------------------- -------------------- -------------------- --------------------
 r17080_Day77TI_135_Env -------------------- -------------------- -------------------- -------------------- --------------------
 r17080_Day77TI_136_Env -------------------- -------------------- -------------------- -------------------- --------------------
 r17080_Day77TI_137_Env -------------------- -------------------- -------------------- -------------------- --------------------
 r17080_Day77TI_138_Env -------------------- -------------------- -------------------- -------------------- --------------------
 r17080_Day77TI_139_Env -------------------- -------------------- -------------------- -------------------- --------------------
    r16080_Day89_16_Env -------------------- ------A------------- -------------------- -------------------- --------------------
    r16080_Day89_17_Env -------------------- ------A------------- -------------------- -------------------- --------------------
    r16080_Day89_18_Env -------------------- ------A------------- -------------------- -------------------- --------------------
    r16080_Day89_19_Env -------------------- ------A------------- -------------------- -------------------- --------------------
    r16080_Day89_20_Env -------------------- -------------------- -------------------- -------------------- --------------------
    r16080_Day89_21_Env -------------------- ------A------------- -------------------- -------------------- --------------------
    r16080_Day89_22_Env -------------------- ------A------------- -------------------- -------------------- --------------------
    r16080_Day89_23_Env -------------------- ------A------------- -------------------- -------------------- --------------------
    r16080_Day89_24_Env -------------------- ------A------------- -------------------- -------------------- --------------------
    r16080_Day89_25_Env -------------------- -------------------- -------------------- -------------------- --------------------
    r16080_Day89_26_Env -------------------- ------A------------- -------------------- -------------------- --------------------
    r16080_Day89_27_Env -------------------- ------A------------- -------------------- -------------------- --------------------
    r16080_Day89_28_Env -------------------- -------------------- -------------------- -------------------- --------------------
    r16080_Day89_29_Env -------------------- --------A----------- -------------------- -------------------- --------------------
    r16080_Day89_30_Env -------------------- ------A------------- -------------------- -------------------- --------------------
    r16080_Day89_31_Env -------------------- ------A------------- -------------------- -------------------- --------------------
r17009_Day131TI_175_Env -------------------- -------------------- -------------------- -------------------- --------------------
r17009_Day131TI_176_Env -------------------- -------------------- -------------------- -------------------- --------------------
r17009_Day131TI_177_Env -------------------- -------------------- -------------------- -------------------- --------------------
r17009_Day131TI_178_Env -------------------- -------------------- -------------------- -------------------- --------------------
r17009_Day131TI_179_Env -------------------- -------------------- -------------------- -------------------- --------------------
r17009_Day131TI_180_Env -------------------- -------------------- -------------------- -------------------- --------------------
r17009_Day131TI_181_Env -------------------- -------------------- -------------------- -------------------- --------------------
r17009_Day131TI_182_Env -------------------- -------------------- -------------------- -------------------- --------------------
r17009_Day131TI_183_Env -------------------- -------------------- -------------------- -------------------- --------------------
r17009_Day131TI_184_Env -------------------- -------------------- -------------------- -------------------- --------------------
r17009_Day131TI_185_Env -------------------- -------------------- -------------------- -------------------- --------------------
r17009_Day131TI_186_Env -------------------- -------------------- -------------------- -------------------- --------------------
r17009_Day131TI_187_Env -------------------- -------------------- -------------------- -------------------- --------------------
r17009_Day131TI_188_Env -------------------- -------------------- -------------------- -------------------- --------------------
r17009_Day131TI_189_Env -------------------- -------------------- -------------------- -------------------- --------------------
r17009_Day131TI_190_Env -------------------- -------------------- -------------------- -------------------- --------------------
r17009_Day131TI_191_Env -------------------- -------------------- -------------------- -------------------- --------------------
r17009_Day131TI_192_Env -------------------- -------------------- -------------------- -------------------- --------------------
r17009_Day131TI_193_Env -------------------- -------------------- -------------------- -------------------- --------------------
r17009_Day131TI_194_Env -------------------- -------------------- -------------------- -------------------- --------------------
r17009_Day131TI_195_Env -------------------- -------------------- -------------------- -------------------- --------------------
r17009_Day131TI_196_Env -------------------- -------------------- -------------------- -------------------- --------------------
r17009_Day131TI_197_Env -------------------- -------------------- -------------------- -------------------- --------------------
r17009_Day131TI_198_Env -------------------- -------------------- -------------------- -------------------- --------------------
r17009_Day131TI_199_Env -------------------- -------------------- -------------------- -------------------- --------------------
r17009_Day131TI_200_Env -------------------- -------------------- -------------------- -------------------- --------------------
r17009_Day131TI_201_Env -------------------- -------------------- -------------------- -------------------- --------------------
r17009_Day131TI_202_Env -------------------- -------------------- -------------------- -------------------- --------------------
 r17058__Day89TI_74_Env -------------------- -------------------- -------------------- -------------------- --------------------
 r17058__Day89TI_75_Env -------------------- --------A----------- -------------------- -------------------- --------------------
 r17058__Day89TI_76_Env -------------------- --------A----------- -------------------- -------------------- --------------------
 r17058__Day89TI_77_Env -------------------- --------A----------- -------------------- -------------------- --------------------
 r17058__Day89TI_78_Env -------------------- --------A----------- -------------------- -------------------- --------------------
 r17058__Day89TI_79_Env -------------------- -------------------- -------------------- -------------------- --------------------
 r17058__Day89TI_80_Env -------------------- -------------------- -------------------- -------------------- --------------------
 r17058__Day89TI_81_Env -------------------- -------------------- -------------------- -------------------- --------------------
 r17058__Day89TI_82_Env -V------------------ -------------------- -------------------- -------------------- --------------------
 r17058__Day89TI_83_Env -------------------- --------A----------- -------------------- -------------------- --------------------
 r17058__Day89TI_84_Env ---------------A---- -------------------- -------------------- -------------------- --------------------
 r17058__Day89TI_85_Env -------------------- -------------------- -------------------- -------------------- --------------------
 r17058__Day89TI_86_Env -------------------- -------------------- -------------------- -------------------- --------------------
 r17058__Day89TI_87_Env -------------------- -------------------- ---S---------------- -------------------- --------------------
 r17080_Day80TI_140_Env -------------------- -------------------- -------------------- -------------------- --------------------
 r17080_Day80TI_141_Env -------------------- -------------------- -------------------- -------------------- --------------------
 r17080_Day80TI_142_Env -------------------- -------------------- -------------------- -------------------- --------------------
 r17080_Day80TI_143_Env -------------------- -------------------- -------------------- -------------------- --------------------
 r17080_Day80TI_144_Env -------------------- -------------------- -------------------- -------------------- --------------------
 r17080_Day80TI_145_Env -------------------- -------------------- -------------------- -------------------- --------------------
 r17080_Day80TI_146_Env -------------------- -------------------- -------------------- -------------------- --------------------
 r17080_Day80TI_147_Env -------------------- -------------------- -------------------- -------------------- --------------------
 r17080_Day80TI_148_Env -------------------- -------------------- -------------------- -------------------- --------------------
 r17080_Day80TI_149_Env -------------------- -------------------- -------------------- -------------------- --------------------
 r17080_Day80TI_150_Env -------------------- -------------------- -------------------- -------------------- --------------------
 r17080_Day80TI_151_Env -------------------- -------------------- -------------------- -------------------- --------------------
 r17080_Day80TI_152_Env -------------------- -------------------- -------------------- -------------------- --------------------
 r17080_Day80TI_153_Env -------------------- -------------------- -------------------- -------------------- --------------------
 r17080_Day80TI_154_Env -------------------- -------------------- -------------------- -------------------- --------------------
 r17080_Day80TI_155_Env -------------------- -------------------- -------------------- -------------------- --------------------
 r17080_Day80TI_156_Env -------------------- -------------------- -------------------- -------------------- --------------------
 r17080_Day80TI_157_Env -------------------- -------------------- -------------------- -------------------- --------------------
 r17080_Day80TI_158_Env -------------------- -------------------- -------------------- -------------------- --------------------
 r17080_Day80TI_159_Env -------------------- -------------------- -------------------- -------------------- --------------------
 r17080_Day80TI_160_Env -------------------- -------------------- -------------------- -------------------- --------------------
 r17080_Day80TI_161_Env -------------------- -------------------- -------------------- -------------------- --------------------
 r17080_Day80TI_162_Env -------------------- -------------------- -------------------- -------------------- --------------------
 r17080_Day80TI_163_Env -------------------- -------------------- -------------------- -------------------- --------------------
 r17080_Day80TI_164_Env -------------------- -------------------- -------------------- -------------------- --------------------

             MAC239_Env KDQAQLNAWGCAFRQVCHTT VPWPNASLTPKWNNETWQEW ERKVDFLEENITALLEEAQI QQEKNMYELQKLNSWDVFGN WFDLASWIKYIQYGVYIVVG  700
     r16080_Day84_1_Env -------------------- -------------------- -------------------- -------------------- --------------------
     r16080_Day84_2_Env -------------------- -------------------- -------------------- -------------------- --------------------
     r16080_Day84_3_Env -------------------- -------------------- --------K----------- -------------------- --------------------
     r16080_Day84_4_Env -------------------- -------------------- -------------------- -------------------- --------------------
     r16080_Day84_5_Env -------------------- -------------------- -------------------- -------------------- --------------------
     r16080_Day84_6_Env -------------------- -------------------- -------------------- -------------------- --------------------
     r16080_Day84_7_Env -------------------- -------------------- -------------------- -------------------- --------------------
     r16080_Day84_8_Env -------------------- -------------------- -------------------- -------------------- --------------------
     r16080_Day84_9_Env -------------------- -------------------- -------------------- -------------------- --------------------
    r16080_Day84_10_Env -------------------- -------------------- -------------------- -------------------- --------------------
    r16080_Day84_11_Env -------------------- -------------------- -------------------- -------------------- ---F----------------
    r16080_Day84_12_Env -------------------- -------------------- -------------------- -------------------- --------------------
    r16080_Day84_13_Env -------------------- -------------------- -------------------- -------------------- --------------------
    r16080_Day84_14_Env -------------------- -------------------- -------------------- -------------------- --------------------
    r16080_Day84_15_Env -------------------- -------------------- -------------------- -------------------- --------------------
 r17009_Day124TI_32_Env -------------------- -------------------- -------------------- -------------------- --------------------
 r17009_Day124TI_33_Env -------------------- -------------------- -------------------- -------------------- --------------------
 r17009_Day124TI_34_Env -------------------- -------------------- -------------------- -------------------- --------------------
 r17009_Day124TI_35_Env -------------------- -------------------- -------------------- -------------------- --------------------
 r17009_Day124TI_36_Env -------------------- -------------------- -------------------- -------------------- --------------------
 r17009_Day124TI_37_Env -------------------- -------------------- -------------------- -------------------- --------------------
 r17009_Day124TI_38_Env -------------------- -------------------- -------------------- -------------------- --------------------
 r17009_Day124TI_39_Env -------------------- -------------------- -------------------- -------------------- --------------------
 r17009_Day124TI_40_Env -------------------- -------------------- -------------------- -------------------- --------------------
 r17009_Day124TI_41_Env -------------------- -------------------- -------------------- -------------------- --------------------
 r17009_Day124TI_42_Env -------------------- -------------------- -------------------- -------------------- --------------------
 r17009_Day124TI_43_Env -------------------- -------------------- -------------------- -------------------- --------------------
 r17009_Day124TI_44_Env -------------------- -------------------- -------------------- -------------------- --------------------
 r17009_Day124TI_45_Env -------------------- -------------------- -------------------- -------------------- --------------------
 r17009_Day124TI_46_Env -------------------- -------------------- -------------------- -------------------- --------------------
 r17009_Day124TI_47_Env -------------------- -------------------- -------------------- -------------------- --------------------
 r17009_Day124TI_48_Env -------------------- -------------------- -------------------- -------------------- --------------------
 r17009_Day124TI_49_Env -------------------- -------------------- -------------------- -------------------- --------------------
 r17009_Day124TI_50_Env -------------------- -------------------- -------------------- -------------------- --------------------
 r17009_Day124TI_51_Env -------------------- -------------------- -------------------- -------------------- --------------------
 r17009_Day124TI_52_Env -------------------- -------------------- -------------------- -------------------- --------------------
 r17009_Day124TI_53_Env -------------------- -------------------- -------------------- -------------------- --------------------
 r17009_Day124TI_54_Env -------------------- -------------------- -------------------- -------------------- --------------------
 r17009_Day124TI_55_Env -------------------- -------------------- -------------------- -------------------- --------------------
 r17009_Day124TI_56_Env -------------------- -------------------- -------------------- -------------------- --------------------
 r17009_Day124TI_57_Env -------------------- -------------------- -------------------- -------------------- --------------------
r17009_Day118TI_165_Env -------------------- -------------------- -------------------- -------------------- --------------------
r17009_Day118TI_166_Env -------------------- -------------------- -------------------- -------------------- --------------------
r17009_Day118TI_167_Env -------------------- -------------------- -------------------- -------------------- --------------------
r17009_Day118TI_168_Env -------------------- -------------------- -------------------- -------------------- --------------------
r17009_Day118TI_169_Env -------------------- -------------------- -------------------- -------------------- --------------------
r17009_Day118TI_170_Env -------------------- -------------------- -------------------- -------K------------ --------------------
r17009_Day118TI_171_Env -------------------- -------------------- -------------------- -------------------- --------------------
r17009_Day118TI_172_Env -------------------- -------------------- -------------------- -------------------- --------------------
r17009_Day118TI_173_Env -------------------- -------------------- -------------------- -------------------- --------------------
  r17058_Day84TI_59_Env -------------------- -------------------- -------------------- -------------------- --------------------
  r17058_Day84TI_60_Env -------------------- ------N------------- -------------------- -------------------- ----------V---------
  r17058_Day84TI_61_Env -------------------- ------N------------- -------------------- -------------------- --------------------
  r17058_Day84TI_62_Env -------------------- -------------------- -------------------- -------------------- --------------------
  r17058_Day84TI_63_Env -------------------- -------------------- -------------------- -------------------- --------------------
  r17058_Day84TI_64_Env -------------------- -------------------- -------------------- -------------------- --------------------
  r17058_Day84TI_65_Env -------------------- -------------------- -------------------- -------------------- --------------------
  r17058_Day84TI_66_Env -------------------- ------N------------- -------------------- -------------------- ----------V---------
  r17058_Day84TI_67_Env -------------------- -------------------- -------------------- -------------------- --------------------
  r17058_Day84TI_68_Env -------------------- -------------------- -------------------- -------------------- --------------------
  r17058_Day84TI_69_Env -------------------- ------N------------- -------------------- -------------------- --------------------
  r17058_Day84TI_70_Env -------------------- ----------E--------- -------------------- -------------------- --------------------
  r17058_Day84TI_71_Env -------------------- --------I----------- -------------------- -------------------- --------------------
  r17058_Day84TI_72_Env -------------------- -------------------- -------------------- -------------------- --------------------
  r17058_Day84TI_73_Env -------------------- ------N------------- -------------------- -------------------- ----------V---------
  r17080_Day73TI_88_Env -------------------- -------------------- -------------------- -------------------- --------------------
  r17080_Day73TI_89_Env -------------------- -------------------- -------------------- -------------------- --------------------
  r17080_Day73TI_90_Env -------------------- -------------------- -------------------- -------------------- --------------------
  r17080_Day73TI_91_Env -------------------- -------------------- -------------------- -------------------- --------------------
  r17080_Day73TI_92_Env -------------------- -------------------- -------------------- -------------------- --------------------
  r17080_Day73TI_93_Env -------------------- -------------------- -------------------- -------------------- --------------------
  r17080_Day73TI_94_Env -------------------- -------------------- -------------------- -------------------- --------------------
  r17080_Day73TI_95_Env -------------------- -------------------- -------------------- -------------------- --------------------
  r17080_Day73TI_96_Env -------------------- -------------------- -------------------- -------------------- --------------------
  r17080_Day73TI_97_Env -------------------- -------------------- -------------------- -------------------- --------------------
  r17080_Day73TI_98_Env -------------------- -------------------- -------------------- -------------------- --------------------
  r17080_Day73TI_99_Env -------------------- -------------------- -------------------- -------------------- --------------------
 r17080_Day73TI_100_Env -------------------- -------------------- -------------------- -------------------- --------------------
 r17080_Day73TI_101_Env -------------------- -------------------- -------------------- -------------------- --------------------
 r17080_Day73TI_102_Env -------------------- -------------------- -------------------- -------------------- --------------------
 r17080_Day73TI_103_Env -------------------- -------------------- -------------------- -------------------- --------------------
 r17080_Day73TI_104_Env -------------------- -------------------- -------------------- -------------------- --------------------
 r17080_Day73TI_105_Env -------------------- -------------------- -------------------- -------------------- --------------------
 r17080_Day73TI_106_Env -------------------- -------------------- -------------------- -------------------- --------------------
 r17080_Day73TI_107_Env -------------------- -------------------- -------------------- -------------------- --------------------
 r17080_Day73TI_108_Env -------------------- -------------------- -------------------- -------------------- --------------------
 r17080_Day73TI_109_Env -------------------- -------------------- -------------------- -------------------- --------------------
 r17080_Day73TI_110_Env -------------------- -------------------- -------------------- -------------------- --------------------
 r17080_Day77TI_111_Env -------------------- -------------------- -------------------- -------------------- --------------------
 r17080_Day77TI_112_Env -------------------- -------------------- -------------------- -------------------- --------------------
 r17080_Day77TI_113_Env -------------------- -------------------- -------------------- -------------------- --------------------
 r17080_Day77TI_114_Env -------------------- -------------------- -------------------- -------------------- --------------------
 r17080_Day77TI_115_Env -------------------- -------------------- -------------------- -------------------- --------------------
 r17080_Day77TI_116_Env -------------------- -------------------- -------------------- -------------------- --------------------
 r17080_Day77TI_117_Env -------------------- -------------------- -------------------- -------------------- --------------------
 r17080_Day77TI_118_Env -------------------- -------------------- -------------------- -------------------- --------------------
 r17080_Day77TI_119_Env -------------------- -------------------- -------------------- -------------------- --------------------
 r17080_Day77TI_120_Env -------------------- -------------------- -------------------- -------------------- --------------------
 r17080_Day77TI_121_Env -------------------- -------------------- -------------------- -------------------- --------------------
 r17080_Day77TI_122_Env -------------------- -------------------- -------------------- -------------------- --------------------
 r17080_Day77TI_123_Env -------------------- -------------------- -------------------- -------------------- --------------------
 r17080_Day77TI_124_Env -------------------- -------------------- -------------------- -------------------- --------------------
 r17080_Day77TI_125_Env -------------------- -------------------- -------------------- -------------------- --------------------
 r17080_Day77TI_126_Env -------------------- -------------------- -------------------- -------------------- --------------------
 r17080_Day77TI_127_Env -------------------- -------------------- -------------------- -------------------- --------------------
 r17080_Day77TI_128_Env -------------------- -------------------- -------------------- -------------------- --------------------
 r17080_Day77TI_129_Env -------------------- -------------------- -------------------- -------------------- --------------------
 r17080_Day77TI_130_Env -------------------- -------------------- -------------------- -------------------- --------------------
 r17080_Day77TI_131_Env -------------------- -------------------- -------------------- -------------------- --------------------
 r17080_Day77TI_132_Env -------------------- -------------------- -------------------- -------------------- --------------------
 r17080_Day77TI_133_Env -------------------- -------------------- -------------------- -------------------- --------------------
 r17080_Day77TI_134_Env -------------------- -------------------- -------------------- -------------------- --------------------
 r17080_Day77TI_135_Env -------------------- -------------------- -------------------- -------------------- --------------------
 r17080_Day77TI_136_Env -------------------- -------------------- -------------------- -------------------- --------------------
 r17080_Day77TI_137_Env -------------------- -------------------- -------------------- -------------------- --------------------
 r17080_Day77TI_138_Env -------------------- -------------------- -------------------- -------------------- --------------------
 r17080_Day77TI_139_Env -------------------- -------------------- -------------------- -------------------- --------------------
    r16080_Day89_16_Env -------------------- -------------------- -------------------- -------------------- --------------------
    r16080_Day89_17_Env -------------------- -------------------- -------------------- -------------------- --------------------
    r16080_Day89_18_Env -------------------- -------------------- -------------------- -------------------- --------------------
    r16080_Day89_19_Env -------------------- -------------------- -------------------- -------------------- --------------------
    r16080_Day89_20_Env -------------------- -------------------- -------------------- -------------------- --------------------
    r16080_Day89_21_Env -------------------- -------------------- -------------------- -------------------- --------------------
    r16080_Day89_22_Env -------------------- -------------------- -------------------- -------------------- --------------------
    r16080_Day89_23_Env -------------------- -------------------- -------------------- -------------------- --------------------
    r16080_Day89_24_Env -------------------- -------------------- -------------------- -------------------- --------------------
    r16080_Day89_25_Env -------------------- -------------------- -------------------- -------------------- --------------------
    r16080_Day89_26_Env -------------------- -------------------- -------------------- -------------------- --------------------
    r16080_Day89_27_Env -------------------- -------------------- -------------------- -------------------- --------------------
    r16080_Day89_28_Env -------------------- -------------------- -------------------- -------------------- --------------------
    r16080_Day89_29_Env -------------------- -------------------- -------------------- -------------------- --------------------
    r16080_Day89_30_Env -------------------- -------------------- -------------------- -------------------- --------------------
    r16080_Day89_31_Env -------------------- -------------------- -------------------- -------------------- --------------------
r17009_Day131TI_175_Env -------------------- -------------------- -------------------- -------------------- --------------------
r17009_Day131TI_176_Env -------------------- -------------------- -------------------- -------------------- --------------------
r17009_Day131TI_177_Env -------------------- -------------------- -------------------- -------------------- --------------------
r17009_Day131TI_178_Env -------------------- -------------------- -------------------- -------------------- --------------------
r17009_Day131TI_179_Env -------------------- -------------------- -------------------- -------------------- --------------------
r17009_Day131TI_180_Env -------------------- -------------------- -------------------- -------------------- --------------------
r17009_Day131TI_181_Env -------------------- -------------------- -------------------- -------------------- --------------------
r17009_Day131TI_182_Env -------------------- -------------------- -------------------- -------------------- --------------------
r17009_Day131TI_183_Env -------------------- -------------------- -------------------- -------------------- --------------------
r17009_Day131TI_184_Env -------------------- -------------------- -------------------- -------------------- --------------------
r17009_Day131TI_185_Env -------------------- -------------------- -------------------- -------------------- --------------------
r17009_Day131TI_186_Env -------------------- -------------------- -------------------- -------------------- --------------------
r17009_Day131TI_187_Env -------------------- -------------------- -------------------- -------------------- --------------------
r17009_Day131TI_188_Env -------------------- -------------------- -------------------- -------------------- --------------------
r17009_Day131TI_189_Env -------------------- -------------------- -------------------- -------------------- --------------------
r17009_Day131TI_190_Env -------------------- -------------------- -------------------- -------------------- --------------------
r17009_Day131TI_191_Env -------------------- -------------------- -------------------- -------------------- --------------------
r17009_Day131TI_192_Env -------------------- -------------------- -------------------- -------------------- --------------------
r17009_Day131TI_193_Env -------------------- -------------------- -------------------- -------------------- --------------------
r17009_Day131TI_194_Env -------------------- -------------------- -------------------- -------------------- --------------------
r17009_Day131TI_195_Env -------------------- -------------------- -------------------- -------------------- --------------------
r17009_Day131TI_196_Env -------------------- -------------------- -------------------- -------------------- --------------------
r17009_Day131TI_197_Env -------------------- -------------------- -------------------- -------------------- --------------------
r17009_Day131TI_198_Env -------------------- -------------------- -------------------- -------------------- --------------------
r17009_Day131TI_199_Env -------------------- -------------------- -------------------- -------------------- --------------------
r17009_Day131TI_200_Env -------------------- -------------------- -------------------- -------------------- --------------------
r17009_Day131TI_201_Env -------------------- -------------------- -------------------- -------------------- --------------------
r17009_Day131TI_202_Env -------------------- -------------------- -------------------- -------------------- --------------------
 r17058__Day89TI_74_Env -------------------- -------------------- -------------------- -------------------- --------------------
 r17058__Day89TI_75_Env -------------------- ----------E--------- -------------------- -------------------- --------------------
 r17058__Day89TI_76_Env -------------------- ----------E--------- -------------------- -------------------- --------------------
 r17058__Day89TI_77_Env -------------------- ----------E--------- -------------------- -------------------- --------------------
 r17058__Day89TI_78_Env -------------------- ----------E--------- -------------------- -------------------- --------------------
 r17058__Day89TI_79_Env -------------------- ------N------------- -------------------- -------------------- ----------V---------
 r17058__Day89TI_80_Env -------------------- -------------------- -------------------- -------------------- --------------------
 r17058__Day89TI_81_Env -------------------- ------N------------- -------------------- -------------------- ----------V---------
 r17058__Day89TI_82_Env -------------------- -----T-------------- -------------------- -------------------- --------------------
 r17058__Day89TI_83_Env -------------------- -------I--E--------- -------------------- -------------------- --------------------
 r17058__Day89TI_84_Env -------------------- -------------------- -------------------- -------------------- --------------------
 r17058__Day89TI_85_Env -------------------- ------N------------- -------------------- -------------------- --------------------
 r17058__Day89TI_86_Env -------------------- -------------------- -------------------- -------------------- --------------------
 r17058__Day89TI_87_Env -------------------- ----------E--------- -------------------- -------------------- --------------------
 r17080_Day80TI_140_Env -------------------- -------------------- -------------------- -------------------- --------------------
 r17080_Day80TI_141_Env -------------------- -------------------- -------------------- -------------------- --------------------
 r17080_Day80TI_142_Env -------------------- -------------------- -------------------- -------------------- --------------------
 r17080_Day80TI_143_Env -------------------- -------------------- -------------------- -------------------- --------------------
 r17080_Day80TI_144_Env -------------------- -------------------- -------------------- -------------------- --------------------
 r17080_Day80TI_145_Env -------------------- -------------------- -------------------- -------------------- --------------------
 r17080_Day80TI_146_Env -------------------- -------------------- -------------------- -------------------- --------------------
 r17080_Day80TI_147_Env -------------------- -------------------- -------------------- -------------------- --------------------
 r17080_Day80TI_148_Env -------------------- -------------------- -------------------- -------------------- --------------------
 r17080_Day80TI_149_Env -------------------- -------------------- -------------------- -------------------- --------------------
 r17080_Day80TI_150_Env -------------------- -------------------- -------------------- -------------------- --------------------
 r17080_Day80TI_151_Env -------------------- -------------------- -------------------- -------------------- --------------------
 r17080_Day80TI_152_Env -------------------- -------------------- -------------------- -------------------- --------------------
 r17080_Day80TI_153_Env -------------------- -------------------- -------------------- -------------------- --------------------
 r17080_Day80TI_154_Env -------------------- -------------------- -------------------- -------------------- --------------------
 r17080_Day80TI_155_Env -------------------- -------------------- -------------------- -------------------- --------------------
 r17080_Day80TI_156_Env -------------------- -------------------- -------------------- -------------------- --------------------
 r17080_Day80TI_157_Env -------------------- -------------------- -------------------- -------------------- --------------------
 r17080_Day80TI_158_Env -------------------- -------------------- -------------------- -------------------- --------------------
 r17080_Day80TI_159_Env -------------------- -------------------- -------------------- -------------------- --------------------
 r17080_Day80TI_160_Env -------------------- -------------------- -------------------- -------------------- --------------------
 r17080_Day80TI_161_Env -------------------- -------------------- -------------------- -------------------- --------------------
 r17080_Day80TI_162_Env -------------------- -------------------- -------------------- -------------------- --------------------
 r17080_Day80TI_163_Env -------------------- -------------------- -------------------- -------------------- --------------------
 r17080_Day80TI_164_Env -------------------- -------------------- -------------------- -------------------- --------------------

             MAC239_Env VILLRIVIYIVQMLAKLRQG YRPVFSSPPSYFQQTHIQQD PALPTREGKERDGGEGGGNS SWPWQIEYIHFLIRQLIRLL TWLFSNCRTLLSRVYQILQP  800
     r16080_Day84_1_Env -------------------- -------------------- ----------G--------- -------------------- --------------------
     r16080_Day84_2_Env -------------------- -----F-------------- ----------G--------- -------------------- --------------------
     r16080_Day84_3_Env -------------------- -------------------- ----------G--------- -------------------- --------------------
     r16080_Day84_4_Env -------------------- -------------------- ----------G--------G -------------------- --------------------
     r16080_Day84_5_Env -------------------- -------------------- ----------G--------- -------------------- --------------------
     r16080_Day84_6_Env -------------------- -------------------- ----------G--------- -------------------- ----------------T---
     r16080_Day84_7_Env -------------------- -------------------- -----G----G---K----- -------------------- --------------------
     r16080_Day84_8_Env -------------------- -------------------- ----------G--------- -------------------- --------------------
     r16080_Day84_9_Env -------------------- -------------------- ----------G--------- -------------------- --------------------
    r16080_Day84_10_Env -------------------- -------------------- ----------G--------- -------------------- --------------------
    r16080_Day84_11_Env -------------------- -------------------- ----------G--------- -------------------- --------------------
    r16080_Day84_12_Env -------------------- -------------------- ----------G--------- -------------------- --------------------
    r16080_Day84_13_Env -------------------- -------------------- ----------G--------- -------------------- --------------------
    r16080_Day84_14_Env -------------------- -------------------- ----------G--------- -------------------- --------------------
    r16080_Day84_15_Env -------------------- -------------------- ----------G--------- -------------------- --------------------
 r17009_Day124TI_32_Env -------------------- -------------------- -------------------- -------------------- --------------------
 r17009_Day124TI_33_Env -------------------- -------------------- -------------------- -------------------- --------------------
 r17009_Day124TI_34_Env -------------------- -------------------- -------------------- -------------------- --------------------
 r17009_Day124TI_35_Env -------------------- -------------------- -------------------- -------------------- --------------------
 r17009_Day124TI_36_Env -------------------- -------------------- -------------------- -------------------- --------------------
 r17009_Day124TI_37_Env -------------------- -------------------- -------------------- -------------------- --------------------
 r17009_Day124TI_38_Env -------------------- -------------------- -------------------- -------------------- --------------------
 r17009_Day124TI_39_Env -------------------- -------------------- -------------------- -------------------- --------------------
 r17009_Day124TI_40_Env -------------------- -------------------- -------------------- -------------------- --------------------
 r17009_Day124TI_41_Env -------------------- -------------------- -------------------- -------------------- --------------------
 r17009_Day124TI_42_Env -------------------- -------------------- -------------------- -------------------- --------------------
 r17009_Day124TI_43_Env -------------------- -------------------- -------------------- -------------------- --------------------
 r17009_Day124TI_44_Env -------------------- -------------------- -------------------- -------------------- --------------------
 r17009_Day124TI_45_Env -------------------- -------------------- -------------------- -------------------- --------------------
 r17009_Day124TI_46_Env -------------------- -------------------- -------------------- -------------------- --------------------
 r17009_Day124TI_47_Env -------------------- -------------------- -------------------- -------------------- --------------------
 r17009_Day124TI_48_Env -------------------- -------------------- -------------------- -------------------- --------------------
 r17009_Day124TI_49_Env -------------------- -------------------- -------------------- -------------------- --------------------
 r17009_Day124TI_50_Env -------------------- -------------------- -------------------- -------------------- --------------------
 r17009_Day124TI_51_Env -------------------- -------------------- -------------------- -------------------- --------------------
 r17009_Day124TI_52_Env -------------------- -------------------- -------------------- -------------------- --------------------
 r17009_Day124TI_53_Env -------------------- -------------------- -------------------- -------------------- --------------------
 r17009_Day124TI_54_Env -------------------- -------------------- -------------------- -------------------- --------------------
 r17009_Day124TI_55_Env -------------------- -------------------- -------------------- -------------------- --------------------
 r17009_Day124TI_56_Env -------------------- -------------------- -------------------- -------------------- --------------------
 r17009_Day124TI_57_Env -------------------- -------------------- -------------------- -------------------- --------------------
r17009_Day118TI_165_Env -------------------- -------------------- -------------------- -------------------- --------------------
r17009_Day118TI_166_Env -------------------- -------------------- -------------------- -------------------- --------------------
r17009_Day118TI_167_Env -------------------- -------------------- -------------------- -------------------- --------------------
r17009_Day118TI_168_Env -------------------- -------------------- -------------------- -------------------- --------------------
r17009_Day118TI_169_Env -------------------- -------------------- -------------------- -------------------- --------------------
r17009_Day118TI_170_Env -------------------- -------------------- -------------------- -------------------- --------------------
r17009_Day118TI_171_Env -------------------- -------------------- -------------------- -------------------- --------------------
r17009_Day118TI_172_Env -------------------- -------------------- -------------------- -------------------- --------------------
r17009_Day118TI_173_Env -------------------- -------------------- -------------------- -------------------- --------------------
  r17058_Day84TI_59_Env -------------------- -------------------- ----------G--------- -------------------- --------------------
  r17058_Day84TI_60_Env -------------------- -------------------- ----------G--------- -------------------- --------------------
  r17058_Day84TI_61_Env -----------------G-- -------------------- ----------G--------- -------------------- --------------------
  r17058_Day84TI_62_Env ------------V------- -------------------- ----------G--------- -------------------- --------------------
  r17058_Day84TI_63_Env -------------------- -------------------- ----------G--------- -------------------- --------------------
  r17058_Day84TI_64_Env -------------------- -------------------- ----------G--------- -------------------- --------------------
  r17058_Day84TI_65_Env -------------------- -------------------- ----------G--------- -------------------- --------------------
  r17058_Day84TI_66_Env -------------------- -------------------- ----------G--------- -------------------- --------------------
  r17058_Day84TI_67_Env -------------------- -------------------- ----------G--------- -------------------- --------------------
  r17058_Day84TI_68_Env -------------------- -------------------- ----------G--------- -------------------- --------------------
  r17058_Day84TI_69_Env -------------------- -------------------- ----------G--------- -------------------- --------------------
  r17058_Day84TI_70_Env -------------------- -------------------- ----------G--------- -------------C------ --------------------
  r17058_Day84TI_71_Env -------------------- -------------------- ----------G--------- -------------------- -----------------F--
  r17058_Day84TI_72_Env -------------------- -------------------- ----------G--------- -------------------- --------------------
  r17058_Day84TI_73_Env -------------------- -------------------- ----------G--------- -------------------- --------------------
  r17080_Day73TI_88_Env -------------------- -------------------- -------------------- -------------------- --------------------
  r17080_Day73TI_89_Env -------------------- -------------------- -------------------- -------------------- --------------------
  r17080_Day73TI_90_Env -------------------- -------------------- -------------------- -------------------- --------------------
  r17080_Day73TI_91_Env -------------------- -------------------- -------------------- -------------------- --------------------
  r17080_Day73TI_92_Env -------------------- -------------------- -------------------- -------------------- --------------------
  r17080_Day73TI_93_Env -------------------- -------------------- -------------------- -------------------- --------------------
  r17080_Day73TI_94_Env -------------------- -------------------- -------------------- -------------------- --------------------
  r17080_Day73TI_95_Env -------------------- -------------------- -------------------- -------------------- --------------------
  r17080_Day73TI_96_Env -------------------- -------------------- -------------------- -------------------- --------------------
  r17080_Day73TI_97_Env -------------------- -------------------- -------------------- -------------------- --------------------
  r17080_Day73TI_98_Env -------------------- -------------------- -------------------- -------------------- --------------------
  r17080_Day73TI_99_Env -------------------- -------------------- -------------------- -------------------- --------------------
 r17080_Day73TI_100_Env -------------------- -------------------- -------------------- -------------------- --------------------
 r17080_Day73TI_101_Env -------------------- -------------------- -------------------- -------------------- --------------------
 r17080_Day73TI_102_Env -------------------- -------------------- -------------------- -------------------- --------------------
 r17080_Day73TI_103_Env -------------------- -------------------- -------------------- -------------------- --------------------
 r17080_Day73TI_104_Env -------------------- -------------------- -------------------- -------------------- --------------------
 r17080_Day73TI_105_Env -------------------- -------------------- -------------------- -------------------- --------------------
 r17080_Day73TI_106_Env -------------------- -------------------- -------------------- -------------------- --------------------
 r17080_Day73TI_107_Env -------------------- -------------------- -------------------- -------------------- --------------------
 r17080_Day73TI_108_Env -------------------- -------------------- -------------------- -------------------- --------------------
 r17080_Day73TI_109_Env -------------------- -------------------- -------------------- -------------------- --------------------
 r17080_Day73TI_110_Env -------------------- -------------------- -------------------- -------------------- --------------------
 r17080_Day77TI_111_Env -------------------- -------------------- -------------------- -------------------- --------------------
 r17080_Day77TI_112_Env -------------------- -------------------- -------------------- -------------------- --------------------
 r17080_Day77TI_113_Env -------------------- -------------------- -------------------- -------------------- --------------------
 r17080_Day77TI_114_Env -------------------- -------------------- -------------------- -------------------- --------------------
 r17080_Day77TI_115_Env -------------------- -------------------- -------------------- -------------------- --------------------
 r17080_Day77TI_116_Env -------------------- -------------------- -------------------- -------------------- --------------------
 r17080_Day77TI_117_Env -------------------- -------------------- -------------------- -------------------- --------------------
 r17080_Day77TI_118_Env -------------------- -------------------- -------------------- -------------------- --------------------
 r17080_Day77TI_119_Env -------------------- -------------------- -------------------- -------------------- --------------------
 r17080_Day77TI_120_Env -------------------- -------------------- -------------------- -------------------- --------------------
 r17080_Day77TI_121_Env -------------------- -------------------- -------------------- -------------------- --------------------
 r17080_Day77TI_122_Env -------------------- -------------------- -------------------- -------------------- --------------------
 r17080_Day77TI_123_Env -------------------- -------------------- -------------------- -------------------- --------------------
 r17080_Day77TI_124_Env -------------------- -------------------- -------------------- -------------------- --------------------
 r17080_Day77TI_125_Env -------------------- -------------------- -------------------- -------------------- --------------------
 r17080_Day77TI_126_Env -------------------- -------------------- -------------------- -------------------- --------------------
 r17080_Day77TI_127_Env -------------------- -------------------- -------------------- -------------------- --------------------
 r17080_Day77TI_128_Env -------------------- -------------------- -------------------- -------------------- --------------------
 r17080_Day77TI_129_Env -------------------- -------------------- -------------------- -------------------- --------------------
 r17080_Day77TI_130_Env -------------------- -------------------- -------------------- -------------------- --------------------
 r17080_Day77TI_131_Env -------------------- -------------------- -------------------- -------------------- --------------------
 r17080_Day77TI_132_Env -------------------- -------------------- -------------------- -------------------- --------------------
 r17080_Day77TI_133_Env -------------------- -------------------- -------------------- -------------------- --------------------
 r17080_Day77TI_134_Env -------------------- -------------------- -------------------- -------------------- --------------------
 r17080_Day77TI_135_Env -------------------- -------------------- -------------------- -------------------- --------------------
 r17080_Day77TI_136_Env -------------------- -------------------- -------------------- -------------------- --------------------
 r17080_Day77TI_137_Env -------------------- -------------------- -------------------- -------------------- --------------------
 r17080_Day77TI_138_Env -------------------- -------------------- -----G-------------- -------------------- --------------------
 r17080_Day77TI_139_Env -------------------- -------------------- -------------------- -------------------- --------------------
    r16080_Day89_16_Env -------------------- -------------------- ----------G--------- -------------------- --------------------
    r16080_Day89_17_Env -------------------- -------------------- ----------G--------- -------------------- --------------------
    r16080_Day89_18_Env -------------------- -------------------- ----------G--------- -------------------- --------------------
    r16080_Day89_19_Env -------------------- -------------------- ----------G--------- -------------------- --------------------
    r16080_Day89_20_Env -------------------- -------------------- ----------G--------- -------------------- --------------------
    r16080_Day89_21_Env -------------------- -------------------- ----------G--------- -------------------- --------------------
    r16080_Day89_22_Env -------------------- -------------------- ----------G--------- -------------------- --------------------
    r16080_Day89_23_Env -------------------- -------------------- ----------G--------- -------------------- --------------------
    r16080_Day89_24_Env -------------------- -------------------- ----------G--------- -------------------- --------------------
    r16080_Day89_25_Env -------------------- ------P------------- ----------G--------- -------------------- --------------------
    r16080_Day89_26_Env -------------------- -------------------- ----------G--------- -------------------- --------------------
    r16080_Day89_27_Env -------------------- -------------------- ----------G--------- -------------------- --------------------
    r16080_Day89_28_Env -------------------- -------------------- ----------G--------- -------------------- --------------------
    r16080_Day89_29_Env -------------------- -------------------- ----------G--------- -------------------- --------------------
    r16080_Day89_30_Env -------------------- -------------------- ----------G--------- -------------------- --------------------
    r16080_Day89_31_Env -------------------- -------------------- ----------G--------- ------------T------- --------------------
r17009_Day131TI_175_Env -------------------- -------------------- -------------------- -------------------- --------------------
r17009_Day131TI_176_Env -------------------- -------------------- -------------------- -------------------- --------------------
r17009_Day131TI_177_Env -------------------- -------------------- -------------------- -------------------- --------------------
r17009_Day131TI_178_Env -------------------- -------------------- -------------------- -------------------- --------------------
r17009_Day131TI_179_Env -------------------- -------------------- -------------------- -------------------- --------------------
r17009_Day131TI_180_Env -------------------- -------------------- -------------------- -------------------- --------------------
r17009_Day131TI_181_Env -------------------- -------------------- -------------------- -------------------- --------------------
r17009_Day131TI_182_Env -------------------- -------------------- -------------------- -------------------- --------------------
r17009_Day131TI_183_Env -------------------- -------------------- -------------------- -------------------- --------------------
r17009_Day131TI_184_Env -------------------- -------------------- -------------------- -------------------- --------------------
r17009_Day131TI_185_Env -------------------- -------------------- -------------------- -------------------- --------------------
r17009_Day131TI_186_Env -------------------- -------------------- -------------------- -------------------- --------------------
r17009_Day131TI_187_Env -------------------- -------------------- -------------------- -------------------- --------------------
r17009_Day131TI_188_Env -------------------- -------------------- -------------------- -------------------- --------------------
r17009_Day131TI_189_Env -------------------- -------------------- -------------------- -------------------- --------------------
r17009_Day131TI_190_Env -------------------- -------------------- -------------------- -------------------- --------------------
r17009_Day131TI_191_Env -------------------- -------------------- -------------------- -------------------- --------------------
r17009_Day131TI_192_Env -------------------- -------------------- -------------------- -------------------- --------------------
r17009_Day131TI_193_Env -------------------- -------------------- -------------------- -------------------- --------------------
r17009_Day131TI_194_Env -------------------- --------------.....- -------------------- -------------------- --------------------
r17009_Day131TI_195_Env -------------------- -------------------- -------------------- -------------------- --------------------
r17009_Day131TI_196_Env -------------------- -------------------- -------------------- -------------------- --------------------
r17009_Day131TI_197_Env -------------------- -------------------- -------------------- -------------------- --------------------
r17009_Day131TI_198_Env -------------------- -------------------- -------------------- -------------------- --------------------
r17009_Day131TI_199_Env -------------------- -------------------- -------------------- -------------------- --------------------
r17009_Day131TI_200_Env -------------------- -------------------- -------------------- -------------------- --------------------
r17009_Day131TI_201_Env -------------------- -------------------- -------------------- -------------------- --------------------
r17009_Day131TI_202_Env -------------------- -------------------- -------------------- -------------------- --------------------
 r17058__Day89TI_74_Env -------------------- -------------------- ----------G--------- -------------------- --------------------
 r17058__Day89TI_75_Env -------------------- -------------------- ----------G--------- -------------C------ --------------------
 r17058__Day89TI_76_Env -------------------- -------------------- ----------G--------- -------------C------ --------------------
 r17058__Day89TI_77_Env -------------------- -------------------- ----------G--------- -------------C------ --------------------
 r17058__Day89TI_78_Env -------------------- -------------------- ----------G--------- -------------C------ --------------------
 r17058__Day89TI_79_Env -------------------- -------------------- ----------G--------- -------------------- --------------------
 r17058__Day89TI_80_Env -------------------- -------------------- ----------G--------- -------------------- --------------------
 r17058__Day89TI_81_Env -------------------- -------------------- ----------G--------- -------------------- --------------------
 r17058__Day89TI_82_Env -------------------- -------------------- ----------G--------- -------------------- --------------------
 r17058__Day89TI_83_Env -------------------- -------------------- ----------G--------- -------------C------ --------------------
 r17058__Day89TI_84_Env -------------------- -------------------- ----------G--------- -------------------- --------------------
 r17058__Day89TI_85_Env -------------------- -------------------- ----------G--------- -------------------- --------------------
 r17058__Day89TI_86_Env -------------------- -------------------- ----------G--------- -------------------- --------------------
 r17058__Day89TI_87_Env -------------------- -------------------- ----------G--------- -------------C------ --------------------
 r17080_Day80TI_140_Env -------------------- -------------------- -------------------- -------------------- --------------------
 r17080_Day80TI_141_Env -------------------- -------------------- -------------------- -------------------- --------------------
 r17080_Day80TI_142_Env -------------------- -------------------- -------------------- -------------------- --------------------
 r17080_Day80TI_143_Env -------------------- -------------------- -------------------- -------------------- --------------------
 r17080_Day80TI_144_Env -------------------- -------------------- -------------------- -------------------- --------------------
 r17080_Day80TI_145_Env -------------------- -------------------- -------------------- -------------------- --------------------
 r17080_Day80TI_146_Env -------------------- -------------------- -------------------- -------------------- --------------------
 r17080_Day80TI_147_Env -------------------- -------------------- -------------------- -------------------- --------------------
 r17080_Day80TI_148_Env -------------------- -------------------- -------------------- -------------------- --------------------
 r17080_Day80TI_149_Env -------------------- -------------------- -------------------- -------------------- --------------------
 r17080_Day80TI_150_Env -------------------- -------------------- -------------------- -------------------- --------------------
 r17080_Day80TI_151_Env -------------------- -------------------- -------------------- -------------------- --------------------
 r17080_Day80TI_152_Env -------------------- -------------------- -------------------- -------------------- --------------------
 r17080_Day80TI_153_Env -------------------- -------------------- -------------------- -------------------- --------------------
 r17080_Day80TI_154_Env -------------------- -------------------- -------------------- -------------------- --------------------
 r17080_Day80TI_155_Env -------------------- -------------------- -------------------- -------------------- --------------------
 r17080_Day80TI_156_Env -------------------- -------------------- -------------------- -------------------- --------------------
 r17080_Day80TI_157_Env -------------------- -------------------- -------------------- -------------------- --------------------
 r17080_Day80TI_158_Env -------------------- -------------------- -------------------- -------------------- --------------------
 r17080_Day80TI_159_Env -------------------- -------------------- -------------------- -------------------- --------------------
 r17080_Day80TI_160_Env -------------------- -------------------- -------------------- -------------------- --------------------
 r17080_Day80TI_161_Env -------------------- -------------------- -------------------- -------------------- --------------------
 r17080_Day80TI_162_Env -------------------- -------------------- -------------------- -------------------- --------------------
 r17080_Day80TI_163_Env -------------------- -------------------- -------------------- -------------------- --------------------
 r17080_Day80TI_164_Env -------------------- -------------------- -------------------- -------------------- --------------------

             MAC239_Env ILQRLSATLQRIREVLRTEL TYLQYGWSYFHEAVQAVWRS ATETLAGAWGDLWETLRRGG RWILAIPRRIRQGLELTLL*  880
     r16080_Day84_1_Env -------------------- -------------------- -------------------- --------------------
     r16080_Day84_2_Env -------------------- -------------------- -------------------- --------------------
     r16080_Day84_3_Env -------------------- -------------------- -------------------- --------------------
     r16080_Day84_4_Env -------------------- -------------------- -------------------- --------------------
     r16080_Day84_5_Env -------------------- -------------------- -------------------- --------------------
     r16080_Day84_6_Env -------------------- -------------------- -------------------- --------------------
     r16080_Day84_7_Env -------------------- -------------------- -------------------- --------------------
     r16080_Day84_8_Env -------------------- -------------------- -------------------- --------------------
     r16080_Day84_9_Env -------------------- -------------------- -------------------- --------------------
    r16080_Day84_10_Env -------------------- -------------------- -------------------- --------------------
    r16080_Day84_11_Env -------------------- -------------------- -------------------- --------------------
    r16080_Day84_12_Env -------------------- -------------------- -------------------- --------------------
    r16080_Day84_13_Env -------------------- -------------------- -------------------- --------------------
    r16080_Day84_14_Env -------------------- -------------------- -------------------- --------------------
    r16080_Day84_15_Env -------------------- -------------------- -------------------- --------------------
 r17009_Day124TI_32_Env -------------------- -------------------- -------------------- --------------------
 r17009_Day124TI_33_Env -------------------- -------------------- -------------------- --------------------
 r17009_Day124TI_34_Env -------------------- -------------------- -------------------- --------------------
 r17009_Day124TI_35_Env -------------------- -------------------- -------------------- --------------------
 r17009_Day124TI_36_Env -------------------- -------------------- -------------------- --------------------
 r17009_Day124TI_37_Env -------------------- -------------------- -------------------- --------------------
 r17009_Day124TI_38_Env -------------------- -------------------- -------------------- --------------------
 r17009_Day124TI_39_Env -------------------- -------------------- -------------------- --------------------
 r17009_Day124TI_40_Env -------------------- -------------------- -------------------- --------------------
 r17009_Day124TI_41_Env -------------------- -------------------- -------------------- --------------------
 r17009_Day124TI_42_Env -------------------- -------------------- -------------------- --------------------
 r17009_Day124TI_43_Env -------------------- -------------------- -------------------- --------------------
 r17009_Day124TI_44_Env -------------------- -------------------- -------------------- --------------------
 r17009_Day124TI_45_Env -------------------- -------------------- -------------------- --------------------
 r17009_Day124TI_46_Env -------------------- -------------------- -------------------- --------------------
 r17009_Day124TI_47_Env -------------------- -------------------- -------------------- --------------------
 r17009_Day124TI_48_Env -------------------- -------------------- -------------------- --------------------
 r17009_Day124TI_49_Env -------------------- -------------------- -------------------- --------------------
 r17009_Day124TI_50_Env -------------------- -------------------- -------------------- --------------------
 r17009_Day124TI_51_Env -------------------- -------------------- -------------------- --------------------
 r17009_Day124TI_52_Env -------------------- -------------------- -------------------- --------------------
 r17009_Day124TI_53_Env -------------------- -------------------- -------------------- --------------------
 r17009_Day124TI_54_Env -----F-------------- -------------------- -------------------- --------------------
 r17009_Day124TI_55_Env -------------------- -------------------- -------------------- --------------------
 r17009_Day124TI_56_Env -------------------- -------------------- -------------------- --------------------
 r17009_Day124TI_57_Env -------------------- -------------------- -------------------- --------------------
r17009_Day118TI_165_Env -------------------- -------------------- -------------------- --------------------
r17009_Day118TI_166_Env -------------------- -------------------- -------------------- --------------------
r17009_Day118TI_167_Env -------------------- -------------------- -------------------- --------------------
r17009_Day118TI_168_Env -------------------- -------------------- -------------------- --------------------
r17009_Day118TI_169_Env -------------------- -------------------- -------------------- --------------------
r17009_Day118TI_170_Env -------------------- -------------------- -------------------- --------------------
r17009_Day118TI_171_Env -------------------- -------------------- -------------------- --------------------
r17009_Day118TI_172_Env -------------------- -------------------- -------------------- --------------------
r17009_Day118TI_173_Env -------------------- -------------------- -------------------- --------------------
  r17058_Day84TI_59_Env -------------------- -------------------- -------------------- --------------------
  r17058_Day84TI_60_Env -------------------- -------------------- -------------------- --------------------
  r17058_Day84TI_61_Env --------------I----- -------------------- -------------------- --------------------
  r17058_Day84TI_62_Env -------------------- -------------------- -------------------- --------------------
  r17058_Day84TI_63_Env -------------------- -------------------- -------------------- --------------------
  r17058_Day84TI_64_Env -------------------- -------------------- -------------------- --------------------
  r17058_Day84TI_65_Env -------------------- -------------------- -------------------- --------------------
  r17058_Day84TI_66_Env -------------------- -------------------- -------------------- --------------------
  r17058_Day84TI_67_Env -------------------- -------------------- -------------------- --------------------
  r17058_Day84TI_68_Env -------------------- -------------------- -------------------- --------------------
  r17058_Day84TI_69_Env -------------------- -------------------- -------------------- --------------------
  r17058_Day84TI_70_Env -------------------- -------------------- -------------------- --------------------
  r17058_Day84TI_71_Env -------------------- -------------------- -------------------- --------------------
  r17058_Day84TI_72_Env -------------------- -------------------- -------------------- --------------------
  r17058_Day84TI_73_Env -------------------- -------------------- -------------------- --------------------
  r17080_Day73TI_88_Env -------------------- ------------V------- -------------------- -------------------
  r17080_Day73TI_89_Env -------------------- ------------V------- -------------------- -------------------
  r17080_Day73TI_90_Env -------------------- -------------------- -------------------- -------------------
  r17080_Day73TI_91_Env -------------------- -------------------- -------------------- -------------------
  r17080_Day73TI_92_Env -------------------- -------------------- -------------------- -------------------
  r17080_Day73TI_93_Env -------------------- ------------V------- -------------------- -------------------
  r17080_Day73TI_94_Env -------------------- -------------------- -------------------- -------------------
  r17080_Day73TI_95_Env -------------------- -------------------- -------------------- -------------------
  r17080_Day73TI_96_Env -------------------- -------------------- -------------------- -------------------
  r17080_Day73TI_97_Env -------------------- -------------------- -------------------- -------------------
  r17080_Day73TI_98_Env -------------------- -------------------- -------------------- -------------------
  r17080_Day73TI_99_Env -------------------- ------------V------- -------------------- -------------------
 r17080_Day73TI_100_Env -------------------- ------------V------- -------------------- -------------------
 r17080_Day73TI_101_Env -------------------- -------------------- -------------------- -------------------
 r17080_Day73TI_102_Env -------------------- ------------V------- -------------------- -------------------
 r17080_Day73TI_103_Env -------------------- -------------------- -------------------- -------------------
 r17080_Day73TI_104_Env -------------------- -------------------- -------------------- -------------------
 r17080_Day73TI_105_Env -------------------- -------------------- -------------------- -------------------
 r17080_Day73TI_106_Env -------------------- -------------------- -------------------- -------------------
 r17080_Day73TI_107_Env -------------------- ------------V------- -------------------- -------------------
 r17080_Day73TI_108_Env -------------------- -------------------- -------------------- -------------------
 r17080_Day73TI_109_Env -------------------- ------------V------- -------------------- -------------------
 r17080_Day73TI_110_Env -------------------- -------------------- -------------------- -------------------
 r17080_Day77TI_111_Env -------------------- ------------V------- -------------------- -------------------
 r17080_Day77TI_112_Env -------------------- -------------------- -------------------- -------------------
 r17080_Day77TI_113_Env -------------------- -------------------- -------------------- -------------------
 r17080_Day77TI_114_Env -------------------- ------------V------- -------------------- -------------------
 r17080_Day77TI_115_Env -------------------- -------------------- -------------------- -------------------
 r17080_Day77TI_116_Env -------------------- -------------------- -------------------- -------------------
 r17080_Day77TI_117_Env -------------------- -------------------- -------------------- -------------------
 r17080_Day77TI_118_Env -------------------- -------------------- -------------------- -------------------
 r17080_Day77TI_119_Env -------------------- ------------V------- -------------------- -------------------
 r17080_Day77TI_120_Env -------------------- -------------------- -------------------- -------------------
 r17080_Day77TI_121_Env -------------------- -------------------- -------------------- -------------------
 r17080_Day77TI_122_Env -------------------- -------------------- -------------------- -------------------
 r17080_Day77TI_123_Env -------------------- -------------------- -------------------- -------------------
 r17080_Day77TI_124_Env -------------------- ------------V------- -------------------- -------------------
 r17080_Day77TI_125_Env -------------------- -------------------- -------------------- -------------------
 r17080_Day77TI_126_Env -------------------- -------------------- -------------------- -------------------
 r17080_Day77TI_127_Env -------------------- ------------V------- -------------------- -------------------
 r17080_Day77TI_128_Env -------------------- -------------------- -------------------- -------------------
 r17080_Day77TI_129_Env -------------------- ------------V------- -------------------- -------------------
 r17080_Day77TI_130_Env --------------I----- -------------------- -------------------- -------------------
 r17080_Day77TI_131_Env -------------------- -------------------- -------------------- -------------------
 r17080_Day77TI_132_Env -------------------- -------------------- -------------------- -------------------
 r17080_Day77TI_133_Env -------------------- -------------------- -------------------- -------------------
 r17080_Day77TI_134_Env -------------------- ------------V------- -------------------- -------------------
 r17080_Day77TI_135_Env -------------------- -------------------- -------------------- -------------------
 r17080_Day77TI_136_Env -------------------- ------------V------- -------------------- -------------------
 r17080_Day77TI_137_Env -------------------- -------------------- -------------------- -------------------
 r17080_Day77TI_138_Env -------------------- -------------------- -------------------- -------------------
 r17080_Day77TI_139_Env -------------------- ------------V------- -------------------- -------------------
    r16080_Day89_16_Env -------------------- -------------------- -------------------- --------------------
    r16080_Day89_17_Env -------------------- -------------------- -------------------- --------------------
    r16080_Day89_18_Env -------------------- -------------------- -------------------- --------------------
    r16080_Day89_19_Env -------------------- -------------------- -------------------- --------------------
    r16080_Day89_20_Env -------------------- -------------------- -------------------- --------------------
    r16080_Day89_21_Env -------------------- -------------------- -------------------- --------------------
    r16080_Day89_22_Env -------------------- -------------------- -------------------- --------------------
    r16080_Day89_23_Env -------------------- -------------------- -------------------- --------------------
    r16080_Day89_24_Env -------------------- -------------------- -------------------- --------------------
    r16080_Day89_25_Env -------------------- -------------------- -------------------- --------------------
    r16080_Day89_26_Env -------------------- -------------------- -------------------- --------------------
    r16080_Day89_27_Env -------------------- -------------------- -------------------- --------------------
    r16080_Day89_28_Env -------------------- -------------------- -------------------- --------------------
    r16080_Day89_29_Env -------------------- -------------------- -------------------- --------------------
    r16080_Day89_30_Env -------------------- -------------------- -------------------- --------------------
    r16080_Day89_31_Env -------------------- -------------------- -------------------- --------------------
r17009_Day131TI_175_Env -------------------- -------------------- -------------------- --------------------
r17009_Day131TI_176_Env -------------------- -------------------- -------------------- --------------------
r17009_Day131TI_177_Env -------------------- -------------------- -------------------- --------------------
r17009_Day131TI_178_Env -------------------- -------------------- -------------------- --------------------
r17009_Day131TI_179_Env -------------------- -------------------- -------------------- --------------------
r17009_Day131TI_180_Env -------------------- -------------------- -------------------- --------------------
r17009_Day131TI_181_Env -------------------- -------------------- -------------------- --------------------
r17009_Day131TI_182_Env -------------------- -------------------- -------------------- --------------------
r17009_Day131TI_183_Env -------------------- -------------------- -------------------- --------------------
r17009_Day131TI_184_Env -------------------- -------------------- -------------------- --------------------
r17009_Day131TI_185_Env -------------------- -------------------- -------------K------ --------------------
r17009_Day131TI_186_Env -------------------- -------------------- -------------------- --------------------
r17009_Day131TI_187_Env -------------------- -------------------- -------------------- --------------------
r17009_Day131TI_188_Env -------------------- -------------------- -------------------- --------------------
r17009_Day131TI_189_Env -------------------- -------------------- -------------------- --------------------
r17009_Day131TI_190_Env -------------------- -------------------- -------------------- --------------------
r17009_Day131TI_191_Env -------------------- -------------------- -------------------- --------------------
r17009_Day131TI_192_Env -------------------- -------------------- -------------------- --------------------
r17009_Day131TI_193_Env -------------------- -------------------- -------------------- --------------------
r17009_Day131TI_194_Env -------------------- -------------------- -------------------- --------------------
r17009_Day131TI_195_Env -------------------- -------------------- -------------------- --------------------
r17009_Day131TI_196_Env -------------------- -------------------- -------------------- --------------------
r17009_Day131TI_197_Env -------------------- -------------------- -------------------- --------------------
r17009_Day131TI_198_Env -------------------- -------------------- -------------------- --------------------
r17009_Day131TI_199_Env -------------------- -------------------- -------------------- --------------------
r17009_Day131TI_200_Env -------------------- -------------------- --------------I----- --------------------
r17009_Day131TI_201_Env -------------------- -------------------- -------------------- --------------------
r17009_Day131TI_202_Env -------------------- -------------------- -------------------- --------------------
 r17058__Day89TI_74_Env -------------------- -------------------- -------------------- --------------------
 r17058__Day89TI_75_Env -------------------- -------------------- -------------------- --------------------
 r17058__Day89TI_76_Env -------------------- -------------------- -------------------- --------------------
 r17058__Day89TI_77_Env -------------------- -------------------- -------------------- --------------------
 r17058__Day89TI_78_Env -------------------- -------------------- -------------------- --------------------
 r17058__Day89TI_79_Env -------------------- -------------------- -------------------- --------------------
 r17058__Day89TI_80_Env -------------------- -------------------- -------------------- --------------------
 r17058__Day89TI_81_Env -------------------- -------------------- -------------------- --------------------
 r17058__Day89TI_82_Env -------------------- -------------------- -------------------- --------------------
 r17058__Day89TI_83_Env -------------------- -------------------- -------------------- --------------------
 r17058__Day89TI_84_Env -------------------- -------------------- -------------------- --------------------
 r17058__Day89TI_85_Env -------------------- -------------------- -------------------- --------------------
 r17058__Day89TI_86_Env -------------------- -------------------- -------------------- --------------------
 r17058__Day89TI_87_Env -------------------- -------------------- -------------------- --------------------
 r17080_Day80TI_140_Env -------------------- -------------------- -------------------- -------------------
 r17080_Day80TI_141_Env -------------------- -------------------- -------------------- -------------------
 r17080_Day80TI_142_Env -------------------- -------------------- -------------------- -------------------
 r17080_Day80TI_143_Env -------------------- ------------V------- -------------------- -------------------
 r17080_Day80TI_144_Env -------------------- -------------------- -------------------- -------------------
 r17080_Day80TI_145_Env -------------------- ------------V------- -------------------- -------------------
 r17080_Day80TI_146_Env -------------------- ------------V------- -------------------- -------------------
 r17080_Day80TI_147_Env -------------------- ------------V------- -------------------- -------------------
 r17080_Day80TI_148_Env -------------------- -------------------- -------------------- -------------------
 r17080_Day80TI_149_Env -------------------- -------------------- -------------------- -------------------
 r17080_Day80TI_150_Env -------------------- ------------V------- -------------------- -------------------
 r17080_Day80TI_151_Env -------------------- -------------------- -------------------- -------------------
 r17080_Day80TI_152_Env -------------------- -------------------- -------------------- -------------------
 r17080_Day80TI_153_Env -------------------- ------------V------- -------------------- -------------------
 r17080_Day80TI_154_Env -------------------- -------------------- -------------------- -------------------
 r17080_Day80TI_155_Env -------------------- ------------V------- -------------------- -------------------
 r17080_Day80TI_156_Env -------------------- -------------------- -------------------- -------------------
 r17080_Day80TI_157_Env -------------------- -------------------- -------------------- -------------------
 r17080_Day80TI_158_Env -------------------- ------------V------- -------------------- -------------------
 r17080_Day80TI_159_Env -------------------- ------------V------- -------------------- -------------------
 r17080_Day80TI_160_Env -------------------- -------------------- -------------------- -------------------
 r17080_Day80TI_161_Env -------------------- ------------V------- -------------------- -------------------
 r17080_Day80TI_162_Env -------------------- ------------V------- -------------------- -------------------
 r17080_Day80TI_163_Env -------------------- -------------------- -------------------- -------------------
 r17080_Day80TI_164_Env -------------------- -------------------- -------------------- -------------------
```
